# Anaerobic sulfur oxidation underlies adaptation of a chemosynthetic symbiont to oxic-anoxic interfaces

**DOI:** 10.1101/2020.03.17.994798

**Authors:** Gabriela F. Paredes, Tobias Viehboeck, Raymond Lee, Marton Palatinszky, Michaela A. Mausz, Siegfried Reipert, Arno Schintlmeister, Andreas Maier, Jean-Marie Volland, Claudia Hirschfeld, Michael Wagner, David Berry, Stephanie Markert, Silvia Bulgheresi, Lena König

## Abstract

Chemosynthetic symbioses occur worldwide in marine habitats, but comprehensive physiological studies of chemoautotrophic bacteria thriving on animals are scarce. Stilbonematinae are coated by monocultures of thiotrophic Gammaproteobacteria. As these nematodes migrate through the redox zone, their ectosymbionts experience varying oxygen concentrations. However, nothing is known about how these variations affect their physiology or metabolism. Here, by applying omics, Raman microspectroscopy and stable isotope labelling, we investigated the effect of oxygen on *Candidatus* Thiosymbion oneisti. Unexpectedly, sulfur oxidation genes were upregulated in anoxic relative to oxic conditions, but carbon fixation genes and incorporation of ^13^C-labeled bicarbonate were not. Instead, several genes involved in carbon fixation were upregulated in oxic conditions, together with genes involved in organic carbon assimilation, polyhydroxyalkanoate (PHA) biosynthesis, nitrogen fixation and urea utilization. Furthermore, in the presence of oxygen, stress-related genes were upregulated together with vitamin biosynthesis genes likely necessary to withstand its deleterious effects, and the symbiont appeared to proliferate less. Based on its physiological response to oxygen, we propose that *Ca.* T. oneisti may exploit anaerobic sulfur oxidation coupled to denitrification to proliferate in anoxic sand. However, the ectosymbiont would still profit from the oxygen available in superficial sand, as the energy-efficient aerobic respiration would facilitate carbon and nitrogen assimilation by the ectosymbiont.

**IMPORTANCE:** Chemoautotrophic endosymbionts are famous for exploiting sulfur oxidization to feed marine organisms with fixed carbon. However, the physiology of thiotrophic bacteria thriving on the surface of animals (ectosymbionts) is less understood. One long standing hypothesis posits that attachment to animals that migrate between reduced and oxic environments would boost sulfur oxidation, as the ectosymbionts would alternatively access sulfide and oxygen, the most favorable electron acceptor. Here, we investigated the effect of oxygen on the physiology of *Candidatus* Thiosymbion oneisti, a Gammaproteobacterium which lives attached to marine nematodes inhabiting shallow water sand. Surprisingly, sulfur oxidation genes were upregulated in anoxic relative to oxic conditions. Furthermore, under anoxia, the ectosymbiont appeared to be less stressed and to proliferate more. We propose that animal-mediated access to oxygen, rather than enhancing sulfur oxidation, would facilitate assimilation of carbon and nitrogen by the ectosymbiont.

## INTRODUCTION

At least six animal phyla and numerous lineages of bacterial symbionts belonging to Alphaproteobacteria, Gammaproteobacteria, and Campylobacteria (formerly Epsilonproteobacteria) (1) engage in chemosynthetic symbioses, rendering the evolutionary success of these associations incontestable (2, 3). Many of these mutualistic associations rely on sulfur-oxidizing (thiotrophic), chemoautotrophic bacterial symbionts. These oxidize reduced sulfur compounds for energy generation in order to fix inorganic carbon (CO_2_) for biomass build-up. Particularly in binary symbioses involving thiotrophic endosymbionts, it is generally accepted that the bacterial chemosynthetic metabolism serves to provide organic carbon for feeding the animal host (reviewed in 2–4). In addition, some chemosynthetic symbionts have been found capable to fix atmospheric nitrogen, albeit symbiont-to-host transfer of fixed nitrogen remains unproven (5, 6). As for the rarer chemosynthetic bacterial-animal associations in which symbionts colonize exterior surfaces (ectosymbioses), fixation of inorganic carbon and transfer of organic carbon to the host has only been unequivocally shown for the microbial community colonizing the gill chamber of the hydrothermal vent shrimp *Rimicaris exoculata* (7).

The majority of thioautotrophic symbioses have been described to rely on reduced sulfur compounds as electron donors and oxygen as terminal electron acceptor (3, 4). However, given that sulfidic and oxic zones are often spatially separated, also owing to abiotic sulfide oxidation (8, 9), chemosynthetic symbioses (1) are typically found where sulfide and oxygen occur in close proximity (e.g. hydrothermal vents, shallow-water sediments) and/or (2) exhibit host behavioral, physiological and anatomical adaptations enabling the symbionts to access both substrates. Among the former adaptations, host-mediated migration across oxygen and sulfide gradients was proposed for shallow water interstitial invertebrates and *Kentrophoros* ciliates (reviewed in 2, 3). The symbionts of Stilbonematinae have also long been hypothesized to associate with their motile nematode hosts to maximize sulfur oxidation-fueled chemosynthesis, by alternatively accessing oxygen in upper sand layers and sulfide in deeper, anoxic sand. This hypothesis was based upon the distribution pattern of Stilbonematinae in sediment cores together with their migration patterns observed in agar columns (10, 11, 12). However, several chemosynthetic symbionts were subsequently shown to use nitrate as an alternative electron acceptor, and nitrate respiration was stimulated by sulfide, suggesting that symbionts may gain energy by respiring nitrate instead of oxygen (13–16). Furthermore, although physiological studies on chemosynthetic symbioses are available (e.g. 17-19), the impact of oxygen on the symbiont central metabolism has not been investigated (remarkably, not even in those that are directly exposed to fluctuating concentrations thereof as in the case of ectosymbionts).

Here, to understand how oxygen affects symbiont physiology, we focused on *Candidatus* Thiosymbion oneisti, a Gammaproteobacterium belonging to the basal family of *Chromatiaceae* (also known as purple sulfur bacteria), which colonizes the surface of the marine nematode *Laxus oneistus* (Stilbonematinae). This group of free-living roundworms represents the only known animals engaging in monospecific ectosymbioses, i.e. each nematode species is ensheathed by a single *Ca.* Thiosymbion phylotype, and, in the case of *Ca*. T. oneisti, the bacteria form a single layer on the cuticle of its host (20–24). Moreover, the rod-shaped representatives of this bacterial genus, including *Ca*. T. oneisti, divide by FtsZ-based longitudinal fission, a unique reproductive strategy which ensures continuous and transgenerational host-attachment (25–27).

Like other chemosynthetic symbionts, *Ca*. Thiosymbion have been considered chemoautotrophic sulfur oxidizers based on several lines of evidence: stable carbon isotope ratios of symbiotic nematodes are comparable to those found in other chemosynthetic symbioses (10); the key enzyme for carbon fixation via the Calvin-Benson-Bassham cycle (RuBisCO) along with elemental sulfur and enzymes involved in sulfur oxidation have been detected (28–30); reduced sulfur compounds (sulfide, thiosulfate) have been shown to be taken up from the environment by the ectosymbionts, to be used as energy source, and to be responsible for the white appearance of the symbiotic nematodes (11, 13, 30); the animals often occur in the sulfidic zone of marine shallow-water sands (12). More recently, the phylogenetic placement and genetic repertoire of *Ca*. Thiosymbion species have equally been supporting the chemosynthetic nature of the symbiosis (6, 24).

In this study, we incubated nematodes associated with *Ca*. T. oneisti under conditions resembling those encountered in their natural environment, and subsequently examined the ectosymbiont transcriptional responses via RNA sequencing (RNA-Seq). In combination with complementary methods such as stable isotope-labeling, proteomics, Raman microspectroscopy and lipidomics, we show that the ectosymbiont exhibits specific metabolic responses to oxygen. Most strikingly, sulfur oxidation but not carbon fixation was upregulated in anoxia. Such a response in their natural environment would challenge the current opinion that sulfur oxidation requires oxygen and drives carbon fixation in chemosynthetic symbioses. We finally present a metabolic scheme of a thiotrophic ectosymbiont experiencing ever-changing oxygen concentrations, in which anaerobic sulfur oxidation coupled to denitrification may represent the preferred metabolism for growth.

## MATERIALS AND METHODS

### Sample collection

*Laxus oneistus* individuals were collected on multiple field trips (2016–2019) at approximately 1 m depth from sand bars off the Smithsonian Field Station, Carrie Bow Cay in Belize (16°48′11.01″N, 88°4′54.42″W). The nematodes were extracted from the sediment by gently stirring the sand and pouring the supernatant seawater through a 212 μm mesh sieve. The retained meiofauna was collected in a petri dish, and single worms of similar size (1-2 mm length, representing adult *L. oneistus*) were handpicked by forceps (Dumont 3, Fine Science Tools, Canada) under a dissecting microscope. *L. oneistus* nematodes were identified based on morphological characteristics (31). Approximately 4 h after collection, the nematodes were used for various incubations as described below. Replicate incubations were started simultaneously.

The spatial distribution of *L. oneistus* as well as concentrations of sulfide (∑H_2_S, i.e. the sum of H_2_S, HS^−^ and S^2−^), dissolved inorganic nitrogen (DIN: nitrate, nitrite, and ammonia) and dissolved organic carbon (DOC) were determined in sediment cores at various depths (Supplementary Materials & Methods, Figure S1A, and Table S1).

### Incubations for RNA sequencing (RNA-Seq)

Batches of 50 *L. oneistus* individuals per incubation and replicate were collected and incubated at two field trips. They were incubated for 24 h in the dark, in either the presence (oxic) or absence (anoxic) of oxygen, in 13 ml-exetainers (Labco, Lampeter, Wales, UK) fully filled with 0.2 µm filtered seawater (Figure S1B). The oxic incubations consisted of two separate experiments of low (hypoxic; three replicates in July 2017) and high (oxic; three replicates in July 2017, three replicates in March 2019) oxygen concentrations. Here, all exetainers were kept open, but only the samples with high oxygen concentrations were submerged in an aquarium constantly bubbled with air (air pump plus; Sera, Heinsberg, Germany). Oxic incubations started with around 195 µM of O_2_ and reached an average of 188 µM after 24 h. Hypoxic incubations started with around 115 µM O_2_ but reached less than 60 µM O_2_ after 24 h. This likely occurred due to nematode oxygen consumption. On the other hand, the anoxic treatments comprised incubations to which either 11 µM of sodium sulfide (Na_2_S*9H_2_O; Sigma-Aldrich, St. Louis, MS, USA) was added (anoxic-sulfidic; three replicates in July 2017), or no sulfide was supplied (anoxic; three replicates in July 2017, two replicates in March 2019) (Figure S1B), and ∑H_2_S concentrations were checked at the beginning and at the end (24 h) of each incubation by spectrophotometric determination following the protocol of Cline (Supplementary Materials & Methods). Anoxic incubations were achieved with the aid of a polyethylene glove bag (AtmosBag; Sigma-Aldrich) that was flushed with N_2_ gas (Fabrigas, Belize City, Belize), together with incubation media and all vials, for at least 1 h before closing. Dissolved oxygen inside the bag, and of every incubation was monitored throughout the 24 h incubation time using a PreSens Fibox 3 trace fiber-optic oxygen meter and non-invasive trace oxygen sensor spots attached to the exetainers (PSt6 and PSt3; PreSens, Regensburg, Germany). For exact measurements of ∑H_2_S and oxygen see Table S2. The seawater used for all incubations had an initial concentration of nitrate and nitrite of 4.2 µM and 0.31 µM, respectively (Supplementary Materials & Methods). Temperature and salinity remained constant throughout all incubations measuring 27-28°C and 33-34 ‰, respectively. After the 24 h incubations, each set of 50 worms was quickly transferred into 2 ml RNA storage solution (13.3 mM EDTA disodium dihydrate pH 8.0, 16.6 mM sodium citrate dihydrate, 3.5 M ammonium sulfate, pH 5.2), kept at 4°C overnight, and finally stored in liquid nitrogen until RNA extraction.

### RNA extraction, library preparation and RNA-Seq

RNA from symbiotic *L. oneistus* was extracted using the NucleoSpin RNA XS Kit (Macherey-Nagel, Düren, Germany). Briefly, sets of 50 worms in RNA storage solution were thawed and the worms were transferred into 90 µl lysis buffer RA1 containing Tris (2-carboxyethyl) phosphine (TCEP) according to the manufacturer’s instructions. The remaining RNA storage solution was centrifuged to collect any detached bacterial cells (10 min, 4°C, 16 100 x g), pellets were resuspended in 10 µl lysis buffer RA1 (plus TCEP) and then added to the worms in lysis buffer. To further disrupt cells, suspensions were vortexed for two minutes followed by three cycles of freeze (−80°C) and thaw (37°C), and homogenization using a pellet pestle (Sigma-Aldrich) for 60 s with a 15 s break after 30 s. Any remaining biological material on the pestle tips was collected by rinsing the tip with 100 µl lysis buffer RA1 (plus TCEP). Lysates were applied to NucleoSpin Filters and samples were processed according to the manufacturer’s instructions, including an on-filter DNA digest. RNA was eluted in 20 µl RNase-free water. To remove any residual DNA, a second DNase treatment was performed using the Turbo DNA-free Kit (Thermo Fisher Scientific, Waltham, MA, USA), RNA was then dissolved in 17 µl RNase-free water, and the RNA quality was assessed using a Bioanalyzer (Agilent, Santa Clara, CA, USA). To check whether all DNA was digested, real-time quantitative PCR using the GoTaq qPCR Master Mix (Promega, Madison, WI, USA) was performed targeting a 158 bp stretch of the *sodB* gene using primers specific for the symbiont (sodB-F: GTGAAGGGTAAGGACGGTTC, sodB-R: AATCCCAGTTGACGATCTCC, 10 µM per primer). Different concentrations of genomic *Ca.* T. oneisti DNA were used as positive controls. The program was as follows: 1x 95°C for 2 min, 40x 95°C for 15 s and 60°C for 1 min, 1x 95°C for 15 s, 55°C to 95°C for 20 min. Next, bacterial and eukaryotic rRNA was removed using the Ribo-Zero Gold rRNA Removal Kit (Epidemiology) (Illumina, San Diego, CA, USA) following the manufacturer’s instructions, but volumes were adjusted for low input RNA (32). In short, 125 µl of Magnetic Beads Solution, 32.5 µl Magnetic Bead Resuspension Solution, 2 µl Ribo-Zero Reaction Buffer and 4 µl Ribo-Zero Removal Solution were used per sample. RNA was cleaned up via ethanol precipitation, dissolved in 9 µl RNase-free water, and rRNA removal was evaluated using the Bioanalyzer RNA Pico Kit (Agilent, Santa Clara, CA, USA). Strand-specific, indexed cDNA libraries were prepared using the SMARTer Stranded RNA-Seq Kit (Takara Bio USA, Mountain View, CA, USA). Library preparation was performed according to the instructions, with 8 µl of RNA per sample as input, 3 min fragmentation time, two rounds of AMPure XP Beads (Beckman Coulter, Brea, CA, USA) clean-up before amplification, and 18 PCR cycles for library amplification. The quality of the libraries was assessed via the Bioanalyzer DNA High Sensitivity Kit (Agilent). Libraries were sequenced on an Illumina HiSeq 2500 instrument (single-read, 100 nt) at the next-generation sequencing facility of the Vienna BioCenter Core Facilities (VBCF, https://www.viennabiocenter.org/facilities/).

### Genome sequencing, assembly and functional annotation

The genome draft of *Ca.* T. oneisti was obtained by performing a hybrid assembly using reads from Oxford Nanopore Technologies (ONT) sequencing and Illumina sequencing. To extract DNA for ONT sequencing and dissociate the ectosymbionts from the host, approximately 800 *Laxus oneistus* individuals were incubated three times for 5 min each in TE-Buffer (10 mM Tris-HCl pH 8.0, 1 mM disodium EDTA pH 8.0). Dissociated symbionts were collected by 10 min centrifugation at 7 000 x g and subsequent removal of the supernatant. DNA was extracted from this pellet using the Blood and Tissue Kit (Qiagen, Hilden, Germany) according to the manufacturer’s instructions. The eluant was further purified using the DNA Clean & Concentrator-5 kit (Zymo Research, Irvine, CA, USA), and the DNA was eluted twice with 10 µl nuclease-free water.

The library for ONT sequencing was prepared using the ONT Rapid Sequencing kit (SQK-RAD002) and sequenced on an R9.4 flow cell (FLO-MIN106) on a MinION for 48 h. Base calling was performed locally with ONT’s Metrichor Agent v1.4.2, and resulting fastq-files were trimmed using Porechop v0.2.1 (https://github.com/rrwick/Porechop). Illumina sequencing reads from a previous study (6), were made available by Harald Gruber-Vodicka (MPI Bremen). Raw reads were filtered: adapters removed and trimmed using bbduk (BBMap v37.22, https://sourceforge.net/projects/bbmap/), with a minimum length of 36 and a minimum phred score of 2. To only keep reads derived from the symbiont, trimmed reads were mapped onto the available genome draft (NCBI Accession FLUZ00000000.1) using BWA-mem v0.7.16a-r1181 (33). Reads that did not map were discarded. The hybrid assembly was performed using SPAdes v3.11 (34) with flags --careful and the ONT reads supplied as --nanopore. Contigs smaller than 200 bp and a coverage lower than 5X were filtered out with a custom python script. The genome completeness was assessed using CheckM v1.0.18 (35) with the gammaproteobacterial marker gene set using the taxonomy workflow. The genome was estimated to be 96.63% complete, contain 1.12% contamination, and was 4.35 Mb in length on 401 contigs with a GC content of 58.7% and N50 value of 27,060 bp.

The genome of *Ca.* T. oneisti was annotated using the MicroScope platform (36), which predicted 5 169 protein-coding genes. To expand the functional annotation provided by MicroScope, predicted proteins were assigned to KEGG pathway maps using BlastKOALA and KEGG Mapper-Reconstruct Pathway (37), gene ontology (GO) terms using Blast2GO v5 (38) and searched for Pfam domains using the hmmscan algorithm of HMMER 3.0 (39, 40). All proteins and pathways mentioned in the manuscript were manually curated. Functional annotations can be found in Data S1.

### Gene expression analyses

Based on quality assessment of raw sequencing reads using FastQC v0.11.8 (41) and prinseq-lite v0.20.4 (42), reads were trimmed and filtered using Trimmomatic v0.39 (43) and prinseq-lite as follows: 18 nucleotides were removed from the 5-prime end (HEADCROP), Illumina adapters were removed (ILLUMINACLIP:TruSeq3-SE.fa:2:30:10), reads were trimmed when the average quality of a five-base sliding window dropped below a phred score of 20 (SLIDINGWINDOW:5:20), 3-prime poly-A tails were trimmed (-trim_tail_right 1), and only reads longer than 24 nucleotides were kept (MINLEN:25). Mapping and expression analysis were done as previously described (44). Briefly, reads were mapped to the *Ca.* T. oneisti genome draft using BWA-backtrack (33) with default settings, only uniquely mapped reads were kept using SAMtools (45), and the number of strand-specific reads per gene was counted using HTSeq in the union mode of counting overlaps (46). On average, 1.4 x 10^6^ (4.4%) reads uniquely mapped to the *Ca.* T. oneisti genome. For detailed read and mapping statistics see Figure S2A.

Gene and differential expression analyses were conducted using the R software environment and the Bioconductor package edgeR v3.28.1 (47–49). Genes were considered expressed if at least two reads in at least two replicates of one of the four conditions could be assigned. Including all four conditions, we found 92.8% of all predicted symbiont protein-encoding genes to be expressed (4 797 genes out of 5 169, Data S1). Log_2_TPM (transcripts per kilobase million) values were calculated by log-transforming TPMs to which library size-adjusted, positive prior counts were added in order to avoid zero TPMs (edgeR function addPriorCount, prior.count=4). Log_2_TPM values were used to assess sample similarities via multidimensional scaling based on Euclidean distances (R Stats package) (49) (Figure S2B), and the average of replicate log_2_TPM values per expressed gene and condition was used to estimate expression strength. Median gene expression of entire metabolic processes and pathways per condition was determined from average log_2_TPMs. A Wilcoxon signed-rank Test was applied to test for significantly different median gene expression between metabolic processes and pathways (R Stats package).

For differential expression analysis, raw data were normalized by the trimmed mean of M-values (TMM) normalization method (edgeR function calcNormFactors) (50), and gene-specific biological variation was estimated (edgeR function estimateDisp). Differential expression was determined using the quasi-likelihood F-test (edgeR functions glmQLFit and glmQLFTest) for pairwise comparisons (between all four conditions) and comparing both anoxic conditions against the average of both oxic conditions. Expression of genes was considered significantly different if their expression changed two-fold between two treatments with a false-discovery rate (FDR) ≤ 0.05 (51). Throughout the paper, all genes meeting these thresholds are either termed differentially expressed or up- or downregulated. However, most follow-up analyses were conducted only considering differentially expressed genes between the anoxic-sulfidic (AS) condition and both oxygenated conditions combined (O, see Results and Figure S2C). For the differential expression analyses between all four conditions see Data S1. Heatmaps show mean-centered expression values to highlight gene expression change.

### Bulk δ^13^C isotopic analysis by Isoprime isotope ratio mass spectrometry (EA-IRMS)

To further analyze the carbon metabolism of *Ca.* T. oneisti, specifically the assimilation of carbon dioxide (CO_2_) by the symbionts in the presence or absence of oxygen, batches of 50 freshly collected, live worms were incubated for 24 h in 150 ml of 0.2 µm-filtered seawater, supplemented with 2 mM (final concentration) of either ^12^C-(natural isotope abundance control) or ^13^C-labeled sodium bicarbonate (Sigma-Aldrich, St. Louis, MS, USA). In addition, a second control was set up (dead control), in which prior to the 24 h incubation with ^13^C-labeled sodium bicarbonate, 50 freshly collected worms were sacrificed by incubating them in a 2% paraformaldehyde/water solution for 12 h.

All three incubations were performed in biological triplicates or quadruplets and set up under anoxic-sulfidic and oxic conditions. Like the RNA-Seq experiment, the oxic incubations consisted of two separate experiments of low (hypoxic) and high (oxic) oxygen concentrations. To prevent isotope dilution through exchange with the atmosphere, both the oxic and anoxic incubations remained closed throughout the 24 h. The procedure was as follows: 0.2 µm-filtered anoxic seawater was prepared as described above and was subsequently used for both oxic and anoxic incubations. Then, compressed air (DAN oxygen kit, Divers Alert Network, USA) and 25 µM of sodium sulfide (Na_2_S*9H_2_O; Sigma-Aldrich, St. Louis, MS, USA) were injected into the oxic and anoxic incubations, respectively, to obtain concentrations resembling the conditions applied in incubations for the RNA-Seq experiment (see Table S2 for details about the incubation conditions and a compilation of the measurement data).

At the end of each incubation (24 h), the nematodes were weighed (0.3-0.7 mg dry weight) into tin capsules (Elemental Microanalysis, Devon, United Kingdom) and dried at 70°C for at least 24 h. Samples were analyzed using a Costech (Valencia, CA USA) elemental analyzer interfaced with a continuous flow Micromass (Manchester, UK) Isoprime isotope ratio mass spectrometer (EA-IRMS) for determination of ^13^C/^12^C isotope ratios. Measurement values are displayed in δ notation (per mil (‰)). A protein hydrolysate, calibrated against NIST reference materials, was used as a standard in sample runs. The achieved precision for δ^13^C was ± 0.2 ‰ (one standard deviation of 10 replicate measurements on the standard).

### Raman microspectroscopy

Three individual nematodes per EA-IRMS incubation were fixed and stored in 0.1 M Trump’s fixative solution (0.1 M sodium cacodylate buffer, 2.5% GA, 2% PFA, pH 7.2, 1 000 mOsm L^-1^) (52), and washed three times for 10 min in 1x PBS (137 mM NaCl, 2.7 mM KCl, 10 mM Na_2_HPO_4_, 1.8 mM KH_2_PO_4_, pH 7.4) before their ectosymbionts were dissociated by sonication for 40 s in 10 µl 1x PBS. 1 µl of each bacterial suspension was spotted on an aluminum-coated glass slide and measured with a LabRAM HR Evolution Raman microspectroscope (Horiba, Kyoto, Japan). 50 individual single-cell spectra were measured from each sample. All spectra were aligned by the phenylalanine peak, baselined using the Sensitive Nonlinear Iterative Peak (SNIP) algorithm of the R package “Peaks” (https://www.rdocumentation.org/packages/Peaks/versions/0.2), and normalized by total spectrum intensity. For calculating the relative sulfur content, the average intensity value for 212-229 wavenumbers (S_8_ peak) was divided by the average intensity of the adjacent flat region of 231-248 wavenumbers (19). For calculating the relative polyhydroxyalkanoate (PHA) content, the average intensity value for 1 723-1 758 wavenumbers (PHA peak) was divided by the average intensity of the adjacent flat region of 1 759-1 793 wavenumbers (53). Median relative sulfur and PHA content (shown as relative Raman intensities) were calculated treating all individual symbiont cells per condition as replicates. Statistically significant differences were determined by applying the non-parametric Kruskal Wallis test, followed by pairwise comparisons.

### Assessment of the percentage of dividing cells

Samples were fixed and ectosymbionts were dissociated from their hosts as described above for Raman microspectroscopy. 1.5 µl of each bacterial suspension per condition were applied to a 1% agarose covered slide (54) and cells were imaged using a Nikon Eclipse NI-U microscope equipped with an MFCool camera (Jenoptik). Images were obtained using the ProgRes Capture Pro 2.8.8 software (Jenoptik) and processed with ImageJ (55). Bacterial cells were manually counted (> 600 per sample) and grouped into constricted (dividing) and non-constricted (non-dividing) cells based on visual inspection (27). The percentage of dividing cells was calculated by counting the total number of dividing cells and the total amount of cells per condition. Statistical tests were performed using Fisher’s exact test.

### Data availability

The assembled and annotated genome of *Ca*. T. oneisti has been deposited at DDBJ/ENA/GenBank under the accession JAAEFD000000000. RNA-Seq data are available at the Gene Expression Omnibus (GEO) database and are accessible through accession number GSE146081.

## RESULTS

### Hypoxic and oxic conditions induce similar expression profiles

To understand how the movement of the animal host across the chemocline affects symbiont physiology and metabolism, we exposed symbiotic worms to sulfide (thereafter used for ∑H_2_S) and oxygen concentrations resembling the ones encountered by *Ca.* T. oneisti in its natural habitat. Previous studies showed that Stilbonematinae live predominantly in anoxic sediment zones with sulfide concentrations < 50 µM (12). To assess whether this applies to *Laxus oneistus* (i.e. *Ca.* T. oneisti’s host) at our collection site (Carrie Bow Cay, Belize), we determined the nematode abundance relative to the sampling depth and sulfide concentration. We found all *L. oneistus* individuals in pore water containing ≤ 25 µM sulfide and only 1.3% of them living in non-sulfidic (0 µM sulfide) surface layers (Figure S1A, Table S1). Therefore, we chose anoxic seawater supplemented with ≤ 25 μM sulfide as the incubation medium (AS condition). To assess the effect of oxygen on symbiont physiology, we additionally incubated the nematodes in hypoxic (< 60 µM oxygen after 24 h) and oxic (> 100 µM after 24 h) seawater. Nitrate, nitrite, ammonium and DOC could be detected throughout the sediment core including the surface layer (Figure S1A, Table S1).

Differential gene expression analysis comparing hypoxic and oxic incubations revealed that only 2.9% of all expressed protein-coding genes differed significantly in their expression. Crucially, this gene set comprised several hypothetical proteins but did not show any significantly enriched metabolic pathways, processes or protein families (Figure S2, Table S3, Data S1). Because the presence of oxygen, irrespective of its concentration, resulted in a similar metabolic response, we treated the samples derived from hypoxic and oxic incubations as biological replicates, and we will hereafter refer to them as the O condition.

Differential gene expression analysis between the AS and O conditions revealed that 20.7% of all expressed protein-coding genes exhibited significantly different expression between these two conditions (Figure S2C), and we will present their expression analysis below.

### Sulfur oxidation genes are upregulated in anoxia

*Ca.* T. oneisti genes encoding for a sulfur oxidation pathway similar to that of the related but free-living purple sulfur bacterium *Allochromatium vinosum* (Figure S3) were highly expressed in both AS and O conditions compared with other central metabolic processes, albeit median gene expression was significantly higher under the AS condition (Figure 1). Consistently, 24 out of the 26 differentially expressed genes involved in sulfur oxidation, were upregulated under AS (Figure 2A). These mostly included genes involved in the cytoplasmic branch of sulfur oxidation, i.e. genes associated with sulfur transfer from sulfur storage globules (*rhd, tusA, dsrE2*), genes encoding for the reverse-acting *dsr* (dissimilatory sulfite reductase) system involved in the oxidation of stored elemental sulfur (S^0^) to sulfite and, finally, also the genes required for further oxidation of sulfite to sulfate in the cytoplasm by two sets of adenylylsulfate (APS) reductase (*aprAB*) along with their membrane anchor (*aprM*) and sulfate adenylyltransferase (*sat*) (56). Genes encoding a quinone-interacting membrane-bound oxidoreductase (*qmoABC*) exhibited the same expression pattern. This is noteworthy, as AprM and QmoABC are hypothesized to have an analogous function, and their co-occurrence is rare among sulfur-oxidizing bacteria (56, 57).

**Figure 1.**
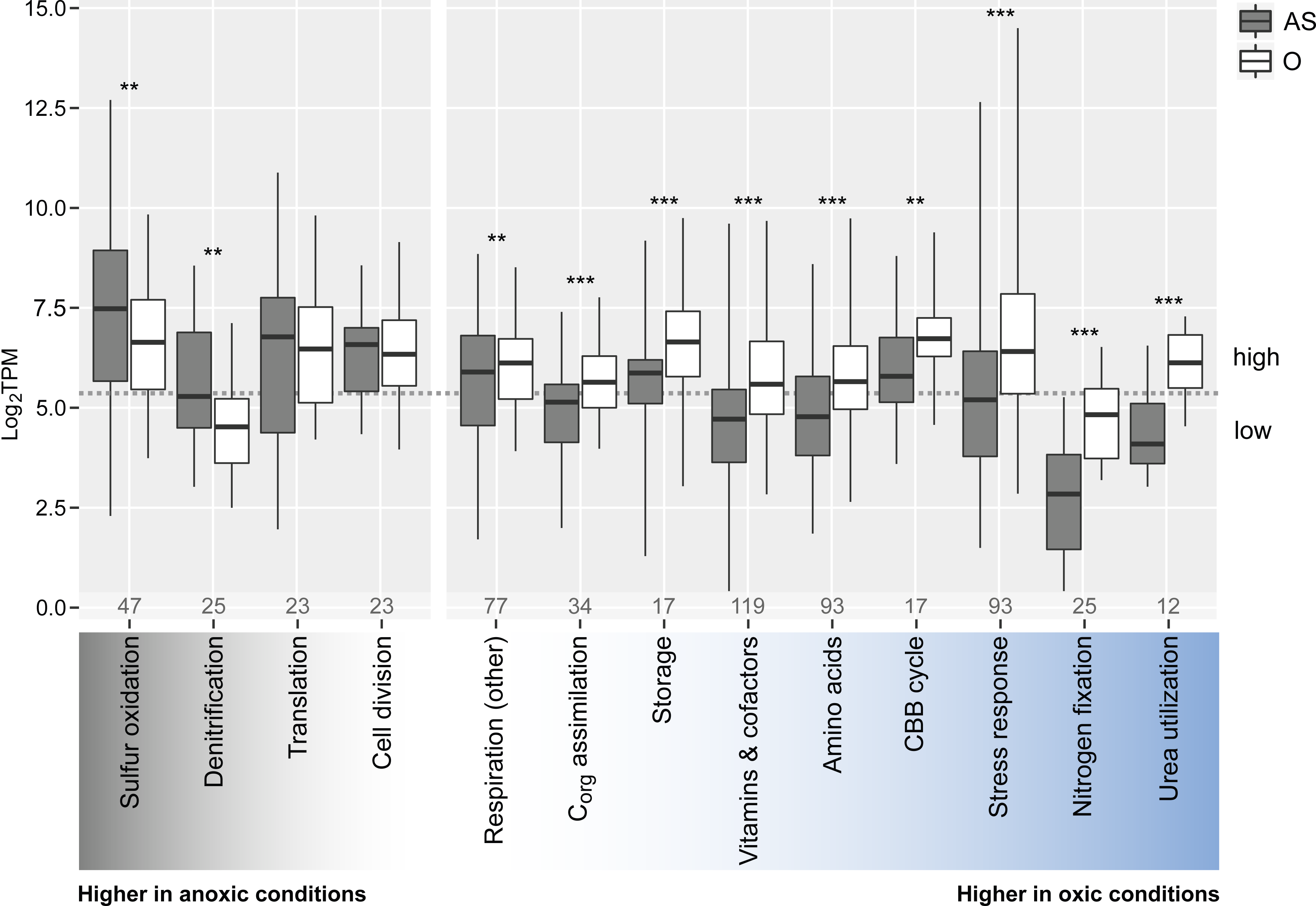
Median gene expression levels of selected *Ca*. T. oneisti metabolic processes under anoxic-sulfidic (AS) versus oxic (O) conditions. All genes involved in a particular process were manually collected and median expression levels (log_2_TPMs, transcripts per kilobase million) per condition and process are shown (horizontal bold lines). Importantly, metabolic processes include both differentially and constitutively expressed genes, and the total number of genes considered are indicated at the bottom of each process. For the specific assignment of genes see Data S1. Note that for the processes designated as ‘Amino acids’, ‘Storage’ and ‘Vitamins & cofactors’ only the expression of the biosynthesis genes was considered. Boxes indicate interquartile ranges (25%-75%), whiskers refer to the minimum and maximum expression values, respectively. The individual processes are ordered according to the difference in median expression between AS and O conditions, i.e. sulfur oxidation (far left) had the largest difference in median expression between the two conditions, with higher median expression in the AS condition, whereas urea utilization (far right) had the largest difference in median expression, with higher median expression in the O condition. Metabolic processes were considered highly expressed when their median expression level was above 5.2 log_2_TPM (dashed grey line), which represents the median expression of all expressed protein-coding genes (n = 4 747) under both conditions. A Wilcoxon Signed-Rank Test was used to test for significantly different median gene expression between conditions (**, p < 0.01; ***, p < 0.001). CBB cycle: Calvin-Benson-Bassham cycle.

**Figure 2.**
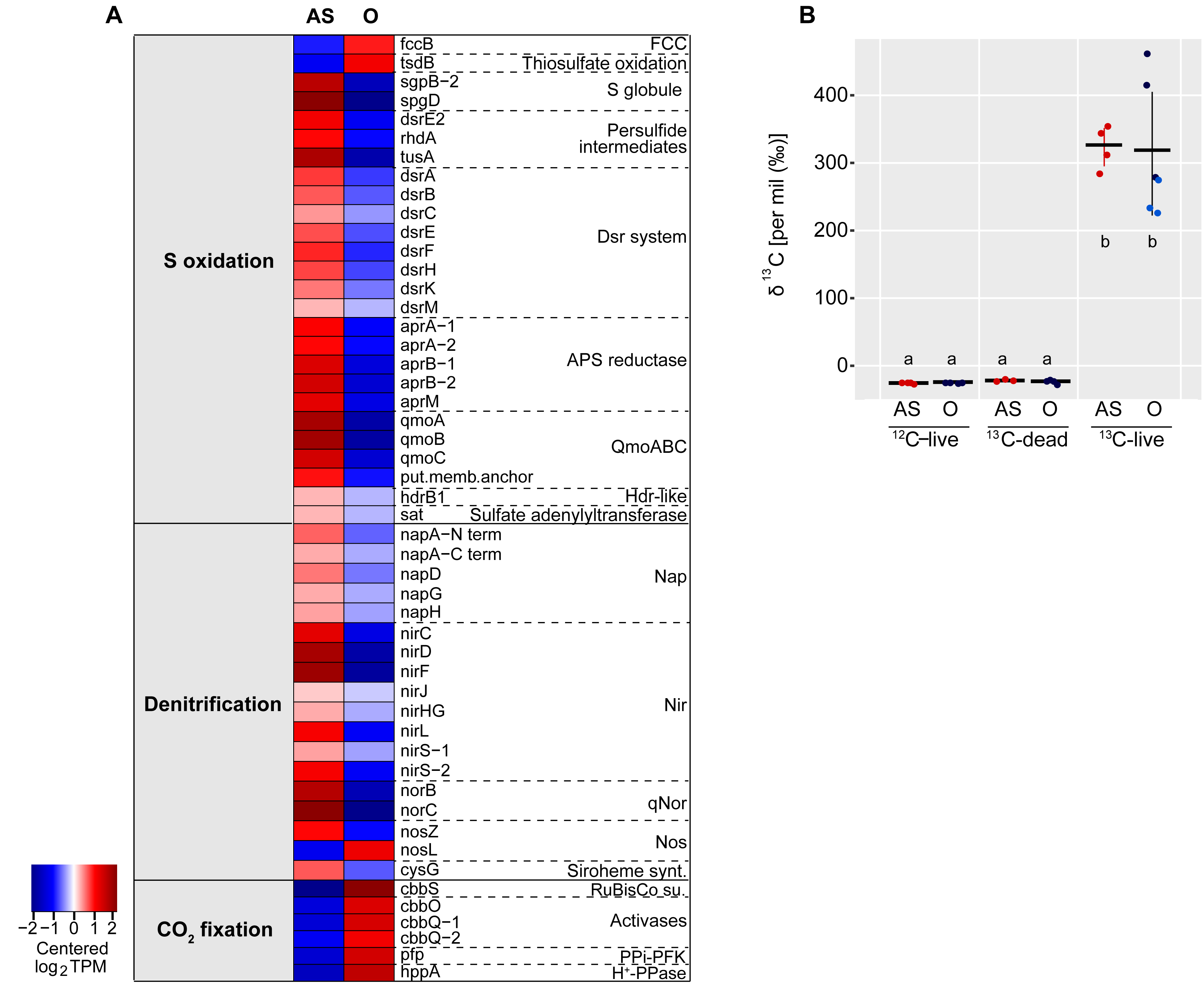
Oxidation of stored sulfur is coupled to denitrification but loosely coupled to CO_2_ fixation under anoxic conditions. (A) Heatmap visualizing only differentially expressed genes (2-fold change, FDR ≤ 0.05) involved in sulfur oxidation, denitrification and CO_2_ fixation via the Calvin-Benson-Bassham cycle between the anoxic-sulfidic (AS) and oxic (O) conditions after 24 h of incubation. Expression levels are visualized by displaying mean-centered log_2_TPMs (transcripts per kilobase million). Upregulation is indicated in red, downregulation in blue. Genes are ordered by function in the respective metabolic pathways. FCC: flavocytochrome c, S: sulfur, Dsr: dissimilatory sulfite reductase, APS: adenylylsulfate, Qmo: quinone-interacting membrane-bound oxidoreductase, Hdr: heterodisulfide reductase, Nap: periplasmic nitrate reductase, Nir: cd1 nitrite reductase, qNor: quinol-dependent nitric oxide reductase, Nos: nitrous oxide reductase, Siroheme synt.: siroheme biosynthesis (heme d precursor), RuBisCo su.: ribulose-1,5-bisphosphate carboxylase/oxygenase small subunit, PPi-PFK: PPi-dependent phosphofructokinase, H^+^-PPase: proton-translocating pyrophosphatase. (B) Relative ^13^C isotope content of symbiotic *L. oneistus* as determined by EA-IRMS after 24 h incubations with ^13^C-labeled bicarbonate under anoxic-sulfidic conditions (AS; red dots), or in the presence of oxygen (O; dark blue: hypoxic, light blue: oxic). Dots refer to the values determined in individual measurements (comprising 50 worms per measurement, for further details see Table S2). Horizontal lines indicate means, error bars correspond to standard deviations. The categories ^12^C-live and ^13^C-dead refer to the natural isotope abundance control and the dead control, respectively. Different lower-case letters indicate significant differences among conditions (one-way ANOVA, Tukey’s post-hoc test, p < 0.05). Importantly, the hypoxic and oxic incubations were not pooled for the EA-IRMS analysis.

Concerning genes involved in the periplasmic branch of sulfur oxidation such as the two types of sulfide-quinone reductases (*sqrA*, s*qrF*; oxidation of sulfide), the Sox system (*soxKAXB*, *soxYZ*) and the thiosulfate dehydrogenase (*tsdA*) both involved in the oxidation of thiosulfate, transcript levels were unchanged between both conditions (Data S1). Only the flavoprotein subunit of the periplasmic flavocytochrome c sulphide dehydrogenase (*fccB*) as well as a cytochrome *c* family protein (*tsdB*) thought to cooperate with TsdA, were downregulated in the absence of oxygen (Figure 2A).

To assess whether the upregulation of sulfur oxidation genes under anoxia was due to the absence of oxygen (and not to the presence of supplemented sulfide in the medium), we performed an additional anoxic incubation where sulfide was not provided (A condition). Differential expression analysis between the anoxic conditions with and without sulfide revealed that transcript levels of 6.3% of all expressed protein-coding genes differed significantly between the two anoxic treatments (Figure S2C). Among them, we found eight genes involved in sulfur oxidation to be upregulated in the presence of sulfide (Data S1). Importantly, however, irrespective of sulfide supplementation, most sulfur oxidation genes were similarly upregulated relative to the O incubation (Figure S4A, Table S3). In addition, proteome data derived from incubations with and without oxygen, but no added sulfide, showed that one copy of AprA and AprM were among the top expressed proteins under anoxia (Table S4, Supplementary Materials & Methods). Raman microspectroscopy revealed that levels of elemental stored sulfur (S^0^) were highest under AS and hypoxic conditions and low or below detection limit under oxic and anoxic without sulfide conditions at the end of the incubations (Figure S4B).

Collectively, sulfur oxidation genes were upregulated under both anoxic conditions irrespective of sulfur storage content and were, conversely, downregulated under hypoxic conditions in spite of elemental sulfur availability in most of the symbiont cells.

### Upregulation of anaerobic respiratory enzymes under AS conditions

Given that sulfur oxidation was upregulated under anoxic conditions, we expected this process to be coupled to the reduction of anaerobic electron acceptors, and nitrate respiration has been shown for symbiotic *L. oneistus* (13). Consistently, genes encoding for components of the four specific enzyme complexes active in denitrification (*nap*, *nir*, *nor*, *nos*), as well as two subunits of the respiratory chain complex III (*petA* and *petB* of the cytochrome bc1 complex, which is known for being involved in denitrification and in the aerobic respiratory chain; 58) were upregulated under AS conditions (Figures 2A and S5).

Besides nitrate respiration, *Ca*. T. oneisti may also utilize polysulfide or thiosulfate as terminal electron acceptors under AS conditions, since we observed an upregulation of all genes encoding either a respiratory polysulfide reductase or thiosulfate reductase (*psrA/phsA*, *psrB/phsB*, *prsC/phsC;* DMSO reductase family, classification based on 59). Concerning other anaerobic electron acceptors, the symbiont has the genetic potential to carry out fumarate reduction (*frdABCD* genes, Figure S3), and the fumarate reductase flavoprotein subunit (*frdA*) was indeed upregulated under AS conditions (Figure S5). We also identified a gene potentially responsible for the biosynthesis of rhodoquinone (*rquA*, Data S1, Figure S3), which acts as an electron carrier in anaerobic respiration in a few other prokaryotic and eukaryotic organisms (60, 61), and could thus replace the missing menaquinone during anaerobic respiration in *Ca*. T. oneisti.

Intriguingly, lipid profiles of the symbiont revealed a change in lipid composition, as well as significantly higher relative abundances of several lyso-phospholipids under anoxia (Figure S6A, Supplementary Materials & Methods), possibly resulting in altered uptake behavior and higher membrane permeability for electron donors and acceptors (62–65). Notably, we also detected lyso-phosphatidylcholine to be significantly more abundant in anoxia (Figure S6B). As the symbiont does not possess any known genes for biosynthesis of this lipid, it may be host-derived. Incorporation of host lipids into symbiont membranes was reported previously (66, 67).

Furthermore, upregulation of the respiratory enzyme glycerol 3-phosphate (G3P) dehydrogenase gene (*glpD*, Figure S5), as well as the substrate-binding subunit of a putative G3P transporter gene (*ugpABCD* genes, Data S1), suggests that putatively host lipid-derived G3P serves as carbon and energy source for the symbiont under anoxia.

Taken together, our data indicate that under AS conditions, the ectosymbiont gains energy by coupling sulfur oxidation to the complete reduction of nitrate to dinitrogen gas. Moreover, the symbiont appears to exploit oxygen-depleted environments for energy generation by utilizing G3P as an additional electron donor, and nitrate, polysulfide or thiosulfate, and fumarate as electron acceptors.

### Upregulation of sulfur oxidation genes is not accompanied by increased expression of carbon fixation genes

Several thioautotrophic symbionts have been shown to use the energy generated by sulfur oxidation for the fixation of inorganic carbon (7, 18, 19, 68–70). Previous studies strongly support that *Ca.* T. oneisti is capable of fixing carbon via an energy-efficient Calvin-Benson-Bassham (CBB) cycle (6, 10, 11, 29,, 71) (Figure S3). In this study, bulk isotope-ratio mass spectrometry (IRMS) conducted with symbiotic nematodes confirmed that they incorporate isotopically-labeled inorganic carbon, and we detected no significant difference in incorporation between any two incubations in the course of 24 h (Figure 2B). To localize the sites of carbon incorporation, we subjected symbiotic nematodes incubated with ^13^C-bicarbonate to nanoscale secondary ion mass spectrometry (NanoSIMS) and detected ^13^C enrichment predominantly within the ectosymbiont (Figure S7, Supplementary Materials & Methods).

Consistent with the evidence for carbon fixation by the ectosymbiont, all genes related to the CBB cycle were detected, both on the transcriptome and the proteome level, with high transcript levels under both AS and O conditions (Figure 1, Data S1). However, the upregulation of sulfur oxidation genes observed under AS did not coincide with an upregulation of carbon fixation genes. On the contrary, the median expression level of all CBB cycle genes was significantly higher in the presence of oxygen (Figure 1). In particular, the transcripts encoding for the small subunit of the key autotrophic carbon fixation enzyme ribulose-1,5-bisphosphate carboxylase/oxygenase RuBisCO (*cbbS*) together with the transcripts encoding its activases (*cbbQ* and *cbbO;* 72), the PPi-dependent 6-phosphofructokinase (PPi-PFK; 73, 74) and the neighboring PPi-energized proton pump (*hppA*) thought to be involved in energy conservation during autotrophic carbon fixation (73, 74) were upregulated under O conditions (Figure 2A). The large subunit of the RuBisCO protein (CbbL, type I-A group, Figure S8) was among the top expressed proteins under anoxic and oxic conditions (Table S4).

In conclusion, (1) upregulation of carbon fixation genes occurred in the presence of oxygen when sulfur oxidation genes were downregulated, while (2) incorporation of inorganic carbon was detected to a similar extent in the presence and absence of oxygen.

### Genes involved in the utilization of organic carbon and PHA storage build-up are upregulated in the presence of oxygen

As anticipated, the nematode ectosymbiont may exploit additional reduced compounds besides sulfide for energy generation. Indeed, *Ca*. T. oneisti possesses the genomic potential to assimilate glyoxylate, acetate and propionate via the partial 3-hydroxypropionate cycle (like the closely related *Olavius algarvensis* γ1-symbiont; 73), and furthermore encodes genes for utilizing additional small organic carbon compounds such as G3P, glycolate, ethanol and lactate (Figure S3, Data S1). With the exception of G3P utilization genes (see above), the expression of genes involved in the assimilation of organic carbon including their putative transporters was significantly higher under O conditions (Figure 1). Among the upregulated genes were *lutB* (involved in the oxidation of lactate to pyruvate; 75), propionyl-CoA synthetase (*prpE,* propanoate assimilation; 76) and two components of a TRAP transporter which most commonly transports carboxylates (77) (Data S1).

These gene expression data imply that the nematode ectosymbiont uses organic carbon compounds in addition to CO_2_ under O conditions, thereby increasing the supply of carbon. Consistent with high carbon availability, genes necessary to synthesize storage compounds such as polyhydroxyalkanoates (PHA), glycogen and trehalose showed an overall higher median transcript level under O conditions (Figure 1). In particular, two key genes involved in the biosynthesis of the PHA compound polyhydroxybutyrate (PHB) – acetyl-CoA acetyltransferase (*phaA*) and a class III PHA synthase subunit (*phaC-2*) – were upregulated in the presence of oxygen. Conversely, we observed upregulation of both PHB depolymerases involved in PHB degradation under AS, and Raman microspectroscopy showed that the median PHA content was slightly lower in symbiont cells under AS relative to both oxic conditions after the incubation period (Figure S9).

We propose that in the presence of oxygen, enhanced mixotrophy (i.e., simultaneous assimilation of inorganic and organic carbon) would result in higher carbon availability reflected by PHA storage build-up and facilitating facultative chemolithoautotrophic synthesis of ATP via the aerobic respiratory chain.

### Upregulation of nitrogen assimilation in the presence of oxygen

It has been shown that high carbon availability is accompanied by high nitrogen assimilation (78–80). Indeed, despite the sensitivity of nitrogenase toward oxygen (81), its key catalytic MoFe enzymes (*nifD, nifK*; 82) and several other genes involved in nitrogen fixation were drastically upregulated in the presence of oxygen (Figures 1 and 3). Moreover, in accordance with a recent study showing the importance of sulfur assimilation for nitrogen fixation (83), genes involved in the assimilation of sulfate, i.e. the sulfate transporters *sulP* and *cys*Z as well as genes encoding two enzymes responsible for cysteine biosynthesis (*cysM*, *cysE*) were also upregulated in the presence of oxygen (Data S1).

**Figure 3.**
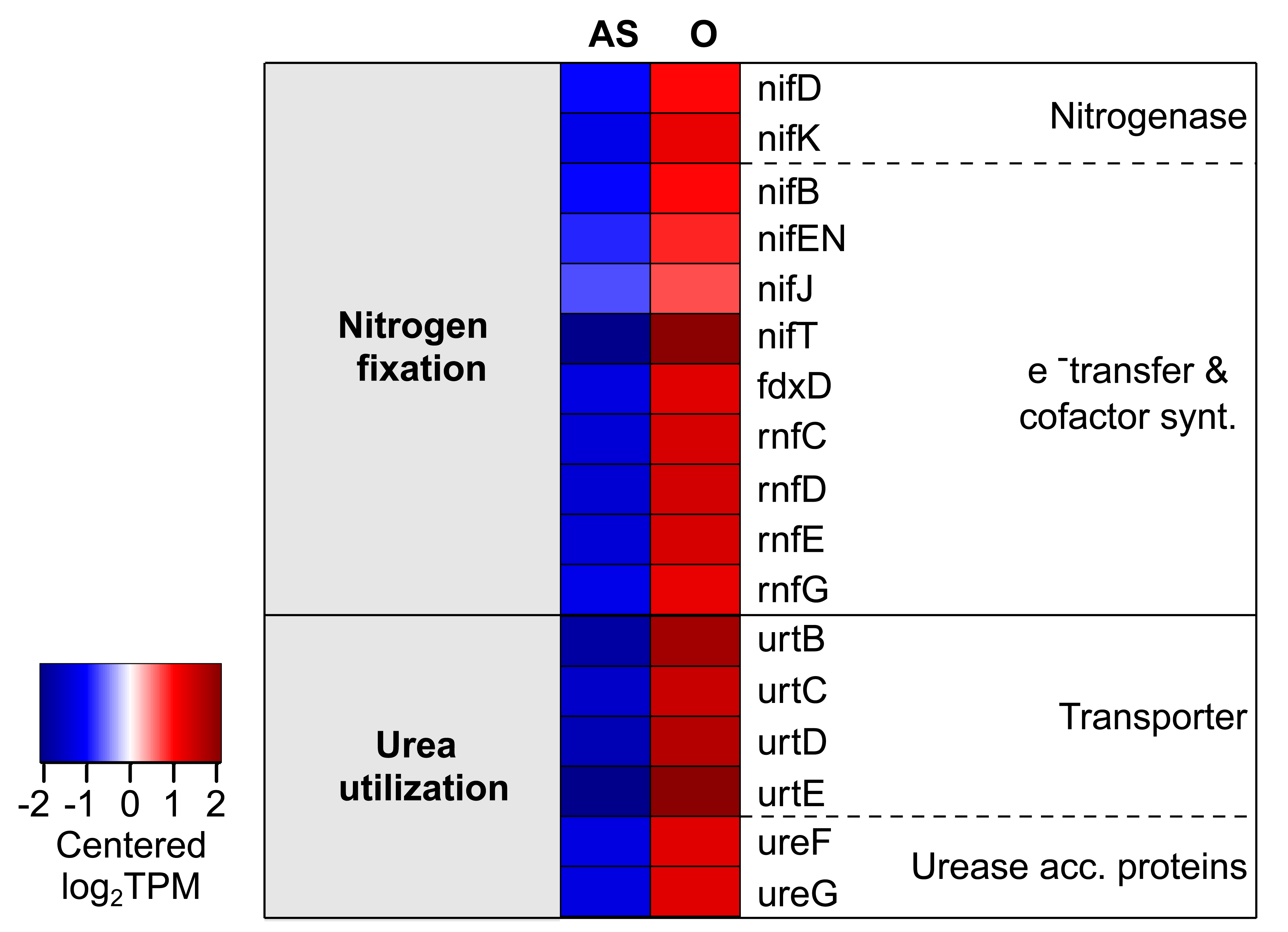
Nitrogen fixation and urea utilization genes are upregulated under oxic conditions. The heatmap only shows genes that were differentially expressed (2-fold change, FDR ≤ 0.05) between anoxic-sulfidic (AS) and oxygenated (O) conditions after 24 h of incubation. Expression levels are displayed as mean-centered log_2_TPMs (transcripts per kilobase million). Upregulation is indicated in red, downregulation in blue. Genes are ordered by function in the respective metabolic pathway. Cofactor synt.: Cofactor biosynthesis, Urease acc. proteins: urease accessory proteins.

Besides nitrogen fixation, genes involved in urea uptake (transporters, *urtCBDE*) and utilization (urease, *ureF* and *ureG*) were also significantly higher transcribed under O conditions (Figures 1 and 3).

In conclusion, genes involved in nitrogen assimilation (from N_2_ or urea) were consistently upregulated in the presence of oxygen, when (1) carbon assimilation was likely higher, and when (2) higher demand for nitrogen is expected due to stress-induced synthesis of vitamins (see section below).

### Upregulation of biosynthesis of cofactors and vitamins and global stress response in the presence of oxygen

Multiple transcripts and proteins associated with diverse bacterial stress responses were among the top expressed in the presence of oxygen (Figure 1 and Table S4). More specifically, heat-shock proteins Hsp70 and Hsp90 were highly abundant (Table S4), and transcripts of heat-shock proteins (Hsp15, Hsp20, Hsp40 and Hsp90) were upregulated (Figure 4A). Besides chaperones, we also detected upregulation of a transcription factor which induces synthesis of Fe-S clusters under oxidative stress (*iscR*; 84) along with several other genes involved in Fe-S cluster formation (Figure 4B, 86), and regulators for redox homeostasis, like thioredoxins, glutaredoxins and peroxiredoxins (86). Furthermore, we observed upregulation of protease genes (*lon*, *ftsH*, *rseP*, *htpX*, *hspQ*; 88–92), genes required for repair of double-strand DNA breaks (such as *radA*, *recB*, *mutSY*, *mfd*; 93–95), and *relA*, known to initiate the stringent response when cells are starved for amino acids (95) (Figure 4A). Amino acid starvation could be caused by a high demand for stress-related proteins under O conditions, and could also explain the upregulation of amino acid biosynthesis pathways under O conditions (96) (Figure 1).

**Figure 4.**
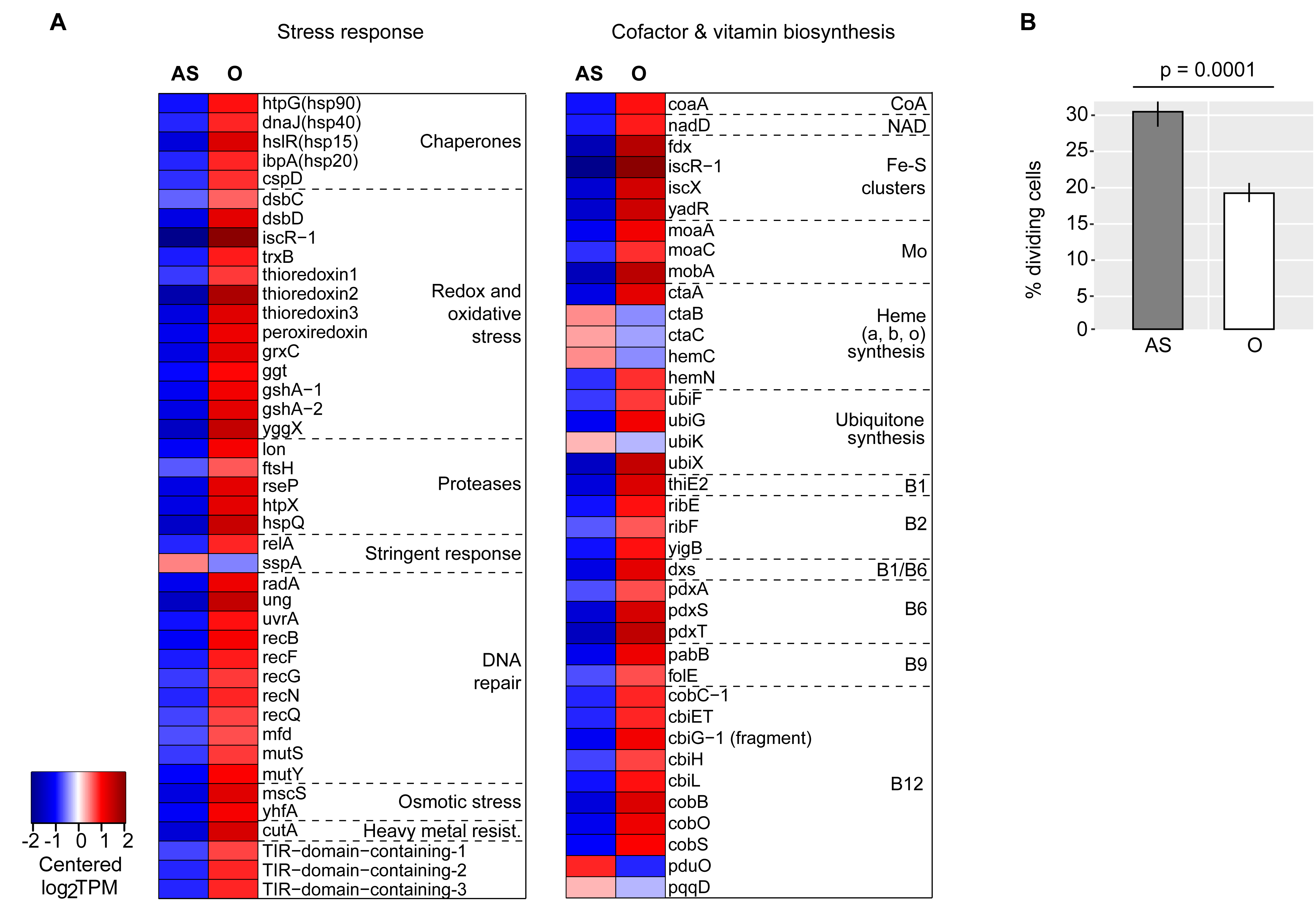
Stress response and vitamin biosynthesis genes are upregulated, and fewer symbiont cells divide in the presence of oxygen. (A) Heatmaps displaying transcript levels of genes involved in stress response, as well as in the biosynthesis of vitamins and cofactors. Expression levels are visualized by displaying mean-centered log_2_TPMs (transcripts per kilobase million). Upregulation is indicated in red, and downregulation in blue. Genes are ordered by function in the respective metabolic pathways. Heavy metal rest.: heavy metal resistance. (B) Bars show the mean percentage of dividing *Ca*. T. oneisti cells upon 24 h incubations under anoxic-sulfidic (AS) and in the presence of oxygen (O). The latter condition includes cells incubated under hypoxic and oxic conditions. A total of 658, 1 009, and 1 923 cells were counted for the AS, hypoxic and oxic condition, respectively. Error bars indicate 95% confidence intervals (Fisher’s exact test).

*SspA*, a gene shown to be important for survival under various stress conditions (97–99), was the only stress-related gene upregulated under AS (Figure 4A).

We hypothesized that the drastic upregulation of stress-related genes observed under oxic conditions would entail an increase in the biosynthesis of vitamins (100–102). Indeed, genes involved in biosynthesis of vitamins such as vitamins B_2_, B_6_, B_9_ and B_12_ were upregulated in the presence of oxygen (Figures 1 and 4A). Notably, the proposed upregulation of nitrogen fixation and urea utilization (see previous section) would support the synthesis of these nitrogen-rich molecules.

The upregulation of stress-related genes under oxic conditions was accompanied by significantly fewer dividing symbiont cells, i.e. 19.2% (under O conditions) versus 30.1% (under AS conditions) (Figure 4B) and downregulation of early (*ftsE, ftsX*) and late (*damX*, *ftsN*) cell division genes (103) (Figure 1, Data S1). Oxygen may therefore elicit a stress response that hampers symbiont proliferation.

## DISCUSSION

This is the first study reporting on the global transcriptional response to oxygen of a thiotrophic animal ectosymbiont, *Ca.* T. oneisti. To the best of our knowledge, the only comparable study is the one conducted on a vent snail endosymbiont subjected to different concentrations of reduced sulfur species (19). Here, we detected a strong transcriptional response of *Ca*. T. oneisti key metabolic processes to oxygen, as well as shifts in protein abundance and lipid composition. Although ongoing comparative host transcriptomics suggests that nematode physiology responds to oxygen (L.K., unpublished data) and although host response likely affects that of *Ca.* T. oneisti, this study exclusively focused on the effect of oxygen on symbiont physiology.

### Experimental design

The concentrations of oxygen and sulfide to which symbiotic nematodes were exposed in our study were chosen based on those measured in the sediment they inhabit at so-called “cool spots”, i.e. sand areas where sulfide concentrations increase gradually with depth but are <50 µM (Figure S1 and 22). Given that in this sand oxygen and sulfide hardly co-occur (104, 105), we did not add any sulfide when nematodes were incubated in the presence of oxygen. While this is reflected in largely depleted symbiont sulfur stores after 24 h under the oxic condition, the symbiont’s sulfur stores remained high under hypoxic and anoxic-sulfidic (AS) condition (Figure S4B). Although we would expect the presence or absence of stored sulfur to affect the symbiont’s metabolism if no external sulfide is present, our data indicate that the gene expression profiles of the sulfur oxidation pathway and other central processes solely followed the presence and absence of oxygen irrespective of sulfide supplementation (Figures 2A and S4A). Residual pre-incubation sulfur stores and other reduced sulfur compounds, such as thiosulfate, present in the seawater may have rendered the symbiont resilient toward the absence of external sulfide.

### Anaerobic sulfur oxidation

Genes involved in sulfur oxidation showed high overall expression compared to other central metabolic processes, indicating that thiotrophy is the predominant energy-generating process for *Ca.* T. oneisti in both oxic and anoxic conditions (Figure 5). Thus, our data strongly support previous observations of Stilbonematinae ectosymbionts performing aerobic and anaerobic sulfur oxidation (11, 13). As the majority of genes involved in denitrification were upregulated under AS conditions (Figures 2A), nitrate likely serves as terminal electron acceptor for anaerobic sulfur oxidation. Importantly, we detected nitrate in the incubation medium, as well as in all sediment layers (Figure S1A, Table S1) at concentrations typical of oligotrophic sediment, in which also the *O. algarvensis* γ3-symbiont is predicted to couple sulfur oxidation to denitrification (73).

**Figure 5.**
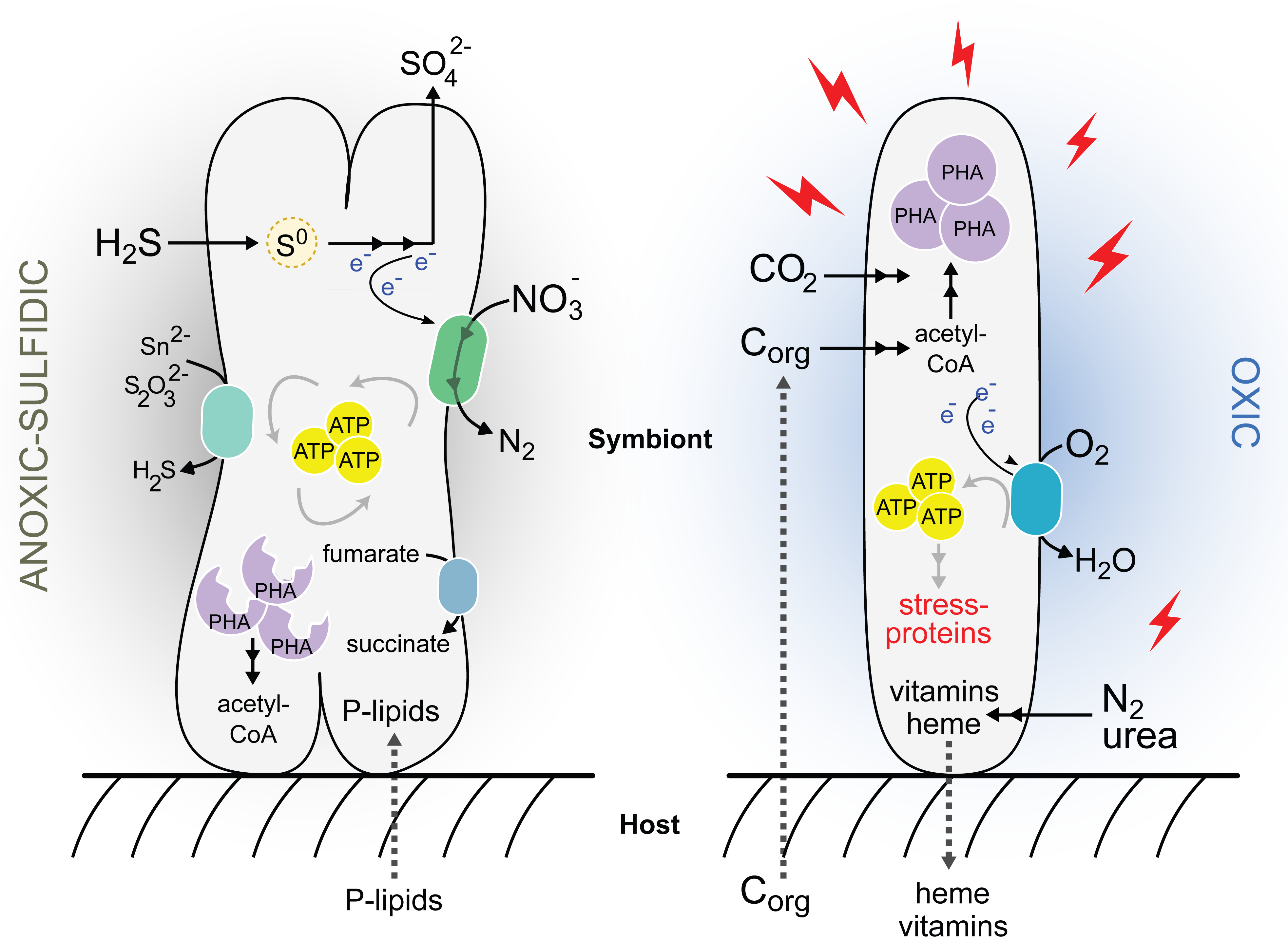
Schematic representation of *Ca*. T. oneisti’s metabolism in deep anoxic and upper oxygenated sand. Our study suggests that in anoxic-sulfidic sediment zones (left) the ectosymbiont performs enhanced anaerobic sulfur oxidation coupled to nitrate reduction to nitrogen gas (denitrification). Additional electron acceptors such as fumarate, polysulfide (Sn^2-^) or thiosulfate (S_2_O_3_^2-^) may also be reduced. The storage compound PHA may serve as a carbon source (in addition to CO_2_) and an additional electron donor. Host-derived phospholipids (P-lipids) may be incorporated into the ectosymbiont’s membrane to increase permeability. In superficial, oxygenated zones (right), oxygen triggers a global stress response that may not only consume energy and dampen proliferation, but may also require vitamin biosynthesis thereby increasing the demand for nitrogen. Small organic carbon compounds (C_org_) putatively excreted by the host and incorporated by the ectosymbiont may contribute to energy generation (and carbon) via aerobic respiration, by the conversion of acetyl CoA via the TCA cycle (not depicted in the figure). Together with autotrophic CO_2_ fixation, C_org_ may increase carbon availability, which would enable *Ca.* T. oneisti to synthesize PHA. Heme and other essential nutrients may be directly or indirectly transferred to the nematode host. Only processes predicted to dominate in one condition over the other are depicted in this model, although they likely occur under both conditions.

Sulfur oxidation in chemosynthetic symbioses is commonly described as an aerobic process required for host survival (3). However, many of these symbiotic organisms likely experience periods of oxygen depletion as would be expected from life at the interface of oxidized and reduced marine environments. Together with previous reports demonstrating nitrate reduction (14–16) and studies showing the genomic potential for using nitrate as terminal electron acceptor (6, 73, 106–108), this study substantiates that nitrate respiration during temporary anoxia could represent a more important strategy for energy conservation among thiotrophic symbionts than currently acknowledged.

While upregulation of sulfur oxidation and denitrification genes in anoxia represents no proof for preferential anaerobic sulfur oxidation, we hypothesize that oxidation of reduced sulfur compounds to sulfate is more pronounced when oxygen is absent. Among the upregulated sulfur oxidation genes, we identified *aprM* and the *qmoABC* complex, both of which are thought to act as electron-accepting units for APS reductase, and therefore rarely co-occur in thiotrophic bacteria (56). The presence and expression of the QmoABC complex could provide a substantial energetic advantage to the ectosymbiont by mediating electron bifurcation (56), in which the additional reduction of a low-potential electron acceptor (e.g. ferredoxin, NAD^+^) could result in optimized energy conservation under anoxic conditions. The maximization of sulfur oxidation under anoxia might even represent a temporary advantage for the host. Indeed, this would be shielded from sulfide poisoning while crawling in a sediment which is free of predators but rich in decomposed organic matter (detritus) (13, 109–111). Due to the dispensability of oxygen for sulfur oxidation, the ectosymbiont may not need to be shuttled to superficial sand by their nematode hosts to oxidize sulfur. Host migration into upper zones of the sediment may therefore primarily reflect the oxygen dependence of the animal host.

In addition to anaerobic sulfur oxidation, the nematode ectosymbiont’s phylogenetic affiliation with facultative anaerobic, anoxygenic phototrophic sulfur oxidizers such as *Allochromatium vinosum* (6, 112), and the presence and expression of yet other anaerobic respiratory complexes (DMSO reductase family enzyme and fumarate reductase) collectively suggest that *Ca*. T. oneisti might be well-adapted to anoxic-sulfidic sediment zones.

### Symbiont proliferation in anoxia

Although a few studies shed light on the molecular cell biology of *Ca*. T. oneisti reproduction (25, 27, 113), up to this study, we did not know how this is influenced by environmental changes. Here, we observed significantly higher numbers of dividing cells under AS conditions (Figure 4B) and therefore, sulfur oxidation coupled to denitrification might represent the ectosymbiont’s preferred strategy to generate energy for growth. We hypothesize that aside from sulfur oxidation, the mobilization of PHA could represent an additional source of ATP (and carbon) supporting symbiont proliferation under AS (Figure S9, Figure 5). Of note, PHA mobilization in anoxia was also shown for *Beggiatoa* spp. (114). On the other hand, several lines of research have shown that stress – experienced by *Ca*. T. oneisti in the presence of oxygen (Figure 4A) – can inhibit bacterial growth (115–121). Importantly, studies on growth preferences of thiotrophic symbionts are scarce, and increased proliferation of a thiotroph with anaerobic electron acceptors (such as nitrate) with respect to using oxygen has never been observed before (122–125).

### Loose coupling of sulfur oxidation and carbon fixation

Reduced sulfur compounds stimulate carbon fixation in thioautotrophic symbionts (7, 11, 18, 19, 68–70, 126, 127). Our bulk isotope-ratio mass spectrometry (EA-IRMS) analysis indicates that, even though expression of the sulfur oxidation pathway was stimulated (Figures 1 and 2), fixation of ^13^C-bicarbonate-derived carbon was not the highest under AS conditions. Instead, carbon fixation appeared unaffected by the presence/absence of oxygen.

Even though, based on EA-IRMS, oxygen did not affect carbon fixation, CBB cycle transcripts in general, and RuBisCO-associated transcripts in particular, were significantly more abundant when oxygen was present (Figures 1 and 2). Upregulation of these genes could be a mechanism to counteract an increased oxygenase activity of RuBisCO in the presence of oxygen, as competition between its two substrates (CO_2_ and O_2_) has been reported to constrain the carbon fixation efficiency of the enzyme (128, 129). Phylogenetic analysis of the ectosymbiont RuBisCO large subunit protein (CbbL) placed it within the type I-A group (Figure S8), whose characterized representatives are adapted to oxic environments (129, 130). The discrepancy between carbon incorporation and transcriptome data could thus reflect a tradeoff between the carboxylase and oxygenase activity of RuBisCo. Of note, fixation of CO_2_ by other carboxylating enzymes may not significantly contribute to the observed pattern of inorganic carbon incorporation. Indeed, acetyl-CoA carboxylase (*acc* genes) is predicted to only act as a biosynthetic carboxylase, whereas the constitutively expressed propionyl-CoA carboxylase (*pccB*) takes part in the partial 3-hydroxypropionate cycle thought to mainly function in assimilation of organic substrates in some thiotrophic symbionts (71, 73, 131). No other known carboxylases are found in the symbiont genome.

Altogether, both lines of evidence point toward a loose coupling between sulfur oxidation and autotrophic carbon fixation. Notably, sulfide oxidation without matching CO_2_ fixation has been described before for the symbiont of *Riftia pachyptila* (132, 133) and an example of extreme decoupling of sulfur oxidation and carbon fixation was recently reported for *Kentrophoros* ectosymbionts. Strikingly, these lack genes for autotrophic carbon fixation altogether and thus represent the first heterotrophic sulfur-oxidizing symbionts (71).

### Oxic mixotrophy

Several chemosynthetic symbionts may engage in mixotrophy (6, 19, 73, 74, 134), and also the nematode ectosymbiont possesses genes for transport of small organic carbon compounds, their assimilation and further metabolization (TCA cycle, glyoxylate shunt). Some of the organic carbon compounds represent typical host waste products (acetate, lactate, propionate) and could therefore be host-derived (73).

The expression of genes involved in transport and assimilation pathways was significantly more pronounced under O than under AS conditions (Figure 1). In addition to assimilating inorganic carbon autotrophically, the ectosymbiont may thus assimilate more organic carbon in the presence of oxygen and, consequently, experience higher carbon availability (Figure 5).

While repression of RuBisCO biosynthesis by organic carbon has been demonstrated (135, 136), simultaneous incorporation of organic and inorganic carbon has been described for several facultative autotrophic bacteria (137–143). Concomitant mixotrophy is thought to be an advantage in oligotrophic environments where nutrients are limiting (140, 144), and CO_2_ derived from the breakdown of organic carbon through decarboxylation can subsequently be reutilized via the CBB cycle (141).

The metabolization of these organic carbon compounds ultimately yields acetyl-CoA, which, in turn, can be further oxidized in the TCA cycle and/or utilized for fatty acid and PHA biosynthesis (Figures S3 and 5). Our transcriptome and Raman microspectroscopy data suggest that *Ca*. T. oneisti favors PHA build-up over its degradation under O conditions (Figure S9). Higher carbon availability in the presence of oxygen resulting in a surplus of acetyl-CoA may cause a nutrient imbalance that could facilitate PHA accumulation as previously shown (145–147). Moreover, it might play a role in resilience against cellular stress, as there is increasing evidence that PHA biosynthesis is enhanced under unfavorable growth conditions such as extreme temperatures, UV radiation, osmotic shock and oxidative stress (148–156). Similar findings have been obtained for pathogenic (157) and symbiotic bacteria of the genus *Burkholderia* (158), with the latter study reporting upregulation of stress response genes and PHA biosynthesis in the presence of oxygen. Finally, oxic biosynthesis of PHA might also prevent excessive accumulation and breakdown of sugars by glycolysis and oxidative phosphorylation, which, in turn, would exacerbate oxidative stress (159).

### Oxic nitrogen assimilation

Despite the oxygen-sensitive nature of nitrogenase (81), we observed a drastic upregulation of nitrogen fixation genes under O conditions (Figures 1 and 3). Besides ammonia production, nitrogen fixation can act as an electron sink during heterotrophic conditions (160, 161). The ectosymbiont may therefore use the nitrogenase to maintain redox balance in the cell when organic carbon is metabolized under oxic conditions. Urea utilization and uptake genes were also upregulated. Although the nematode host likely lacks the urea biosynthetic pathway (L.K., unpublished data), this compound is one of the most abundant organic nitrogen substrates in marine ecosystems, as well as in animal-inhabited (oxygenated) sand (162, 163). The apparent increase in nitrogen assimilation in the presence of oxygen could thus be a result of an increased demand for nitrogen driven by the biosynthesis of nitrogen-rich compounds such as vitamins and cofactors potentially required to survive oxidative stress (Figures 1, 4 and 5). Indeed, the upregulation of the urea uptake system and urease accessory proteins as well as the aforementioned stress-related *relA* gene has been shown to be a response to nitrogen limitation in other systems (164, 165); nitrogen imbalance may have also induced PHA accumulation under oxic conditions (145–147). The role of vitamins in protecting cells against the deleterious effects of oxygen has been shown for animals (166, 167), and the importance of riboflavin for bacterial survival under oxidative stress has previously been reported (100, 102). Along this line of thought, oxygen-exposed *Ca.* T. oneisti upregulated glutathione and thioredoxin, which are known to play a pivotal role in scavenging reactive oxygen species (ROS) (168). Their function directly (or indirectly) requires vitamin B_2_, B_6_ and B_12_ as cofactors. More specifically, thioredoxin reductase (*trxB*) requires riboflavin (vitamin B_2_) in the form of flavin adenine dinucleotide (FAD) (169); cysteine synthase (*cysM*) and glutamate synthases (two-subunit *gltB*/*gltD,* one-subunit *gltS*) involved in the biosynthesis of the glutathione precursors L-cysteine and L-glutamate depend on vitamin B_6_, FAD and riboflavin in the form of flavin mononucleotide (FMN) (170, 171). As for cobalamin, it was thought that this vitamin only played an indirect role in oxidative stress resistance (172), by being a precursor of S-adenosyl methionine (SAM), a substrate involved in the synthesis of glutathione via the methionine metabolism (and the transulfuration pathway), and in preventing the Fenton reaction (173, 174). However, its direct involvement in the protection of chemolithoautotrophic bacteria against oxidative stress has also been illustrated (101).

In summary, in the presence of oxygen, the upregulation of genes involved in biosynthesis of vitamins B_2_, B_6_ and B_12_ along with antioxidant systems and their key precursor genes *cysM* and B_12_-dependent-methionine synthase *metH*, suggests that the ectosymbiont requires increased levels of these vitamins to cope with oxidative stress (Figure 5).

### Evolutionary considerations

Anaerobic sulfur oxidation, increased symbiont proliferation and downregulation of stress-related genes lead us to hypothesize that *Ca.* T. oneisti evolved from a free-living bacterium that mostly, if not exclusively, inhabited anoxic sand zones. In support of this, the closest relatives of the nematode ectosymbionts are free-living sulfur oxidizers thriving under anoxic conditions (i.e. *Allochromatium vinosum*, *Thioflavicoccus mobilis*, *Marichromatium purpuratum*) (6, 175). Eventually, advantages such as protection from predators or utilization of host waste products (e.g. fermentation products, ammonia) may have been driving forces that led to the *Ca*. Thiosymbion*-*Stilbonematinae symbioses. As the association became more and more stable, the symbiont optimized (or acquired) mechanisms to resist oxidative stress, as well as metabolic pathways to most efficiently exploit the metabolic potential of oxygenated sand zones (mixotrophy, nitrogen assimilation, vitamin and cofactor biosynthesis). From the *L. oneistus* nematode perspective, the acquired “symbiotic skin” enabled it to tolerate the otherwise poisonous sulfide and to thrive in sand virtually devoid of predators, but rich in decomposed organic matter.

## ACKNOWLEDGEMENTS

This work was supported by the Austrian Science Fund (FWF) grant P28743 (T. V., S. and L. K.) and the DK plus: Microbial Nitrogen Cycling (G. F. P.). We are indebted to Florian Goldenberg, Patrick Hyden and Thomas Rattei (Division of Computational Systems Biology, University of Vienna) for providing and maintaining the Life Science Compute Cluster (LiSC) and help in preparing the MinION sequencing library for *Ca.* T. oneisti at the University of Vienna. Harald Gruber-Vodicka from the MPI Bremen generously provided Illumina raw reads to aid the assemblies of ectosymbiont genomes. We are very grateful to Wiebke Mohr, Nikolaus Leisch and Nicole Dubilier from the MPI for Marine Microbiology (Bremen) for continuous technical support with the stable isotope-based techniques. We appreciate Tjorven Hinzke’s advice on metaproteome statistics and Carolina Reyes for her input on the nitrogen metabolism. We thank Yin Chen for providing the facilities for lipidomic analysis and Eleonora Silvano for assistance with lipid extractions, and Jana Matulla’s and Sebastian Grund’s excellent technical work during protein sample preparation and MS analysis, respectively. Also, our sincere gratitude to the Carrie Bow Cay Laboratory, Caribbean Coral Reef Ecosystem Program and Station Manager Zach Foltz for his continuous help during field work. Finally, we were very much helped and inspired by insightful discussions with Monika Bright, Jörg A. Ott, Christa Schleper, Simon K.-M.R. Rittmann, Filipa Sousa and Jillian Petersen.

## SUPPLEMENTAL MATERIAL

### Supplemental Figures

**Figure S1. Natural versus experimental conditions.** (A) The left panel shows *Laxus oneistus* total counts per 6 cm core subsection from all 8 sandbars (horizontal beige bars) and corresponding mean sulfide (ΣH_2_S) concentrations (μM, grey line). The right panel shows mean nitrate and nitrate concentrations through depth in the same sandbars of Carrie Bow Cay, Belize. Error bars represent the standard error of the mean. Note that a few nitrate and nitrite data points are derived from only two or one sandbars (Table S1). (B) Experimental set up of incubations for RNA-Seq, EA-IRMS and Raman microspectroscopy. Batches of 50 *L. oneistus* were incubated under different oxygen concentrations: anoxic with sulfide (0 μM O_2_, ≤ 25 μM sodium sulfide added), anoxic without sulfide (0 μM O_2_), hypoxic (< 60 μM O_2_ after 24 h), and oxic (> 100 μM O_2_ after 24 h). The box around the anoxic incubation vials indicates that these incubations were carried out in a polyethylene isolation chamber. Given the similarity of gene expression profiles between the hypoxic and oxic samples (see Figure S2), most of the follow-up analyses were conducted by treating the hypoxic and oxic samples as biological replicates (O), and comparing the O condition to the anoxic-sulfidic (AS) condition. All incubations were performed in 0.2 μM-filtered seawater and in biological triplicates.

**Figure S2. RNA sequencing statistics, sample similarity and differential gene expression.** (A) RNA sequencing and mapping statistics. Sequencing reads were mapped to the symbiont genome assembly consisting of 401 contigs and 5 169 protein-coding genes. The total number of reads refers to the number of reads after quality filtering and trimming, and the number of reads mapped to the genome (i.e. genes, intergenic regions and antisense regions) only includes uniquely mapped reads. (B) Similarity between samples based on Euclidean distances between expression values (log2TPMs), visualized by means of multidimensional scaling (MDS). A total of 4 797 protein-coding genes (92.8%) were detected to be expressed. Most of the follow-up analyses were conducted comparing the anoxic-sulfidic conditions (AS, red circle) to conditions in which oxygen was present (O, blue circle). (C) Differential expression (DE) analysis between hypoxic and oxic samples revealed that the number of DE genes was low (2.9% of all expressed genes), and thus hypoxic and oxic samples were treated as biological replicates. 20.7% of all expressed genes were differentially expressed between AS and O conditions. Genes were considered differentially expressed if their expression changed twofold with a false-discovery rate (FDR) ≤ 0.05.

**Figure S3. Schematic representation of central metabolic pathways present in the *Ca*. T. oneisti genome.** All gene names are indicated in Data S1. Organic carbon compounds such as acetate, lactate, propionate and glycerol 3-phosphate (glycerol-3P) could be host-derived. The respiratory chain of oxygen respiration (O_2_ resp., blue) consists of NADH dehydrogenase (*nuo* genes, complex I), succinate dehydrogenase (*sdh* genes, complex II), the cytochrome bc1 complex (*pet* genes, complex III) and an aa3-type cytochrome c oxidase (*cta* genes, complex IV). Note that *Ca*. T. oneisti only encodes a single complex IV enzyme. Grey: enzymes, brown: transporters, orange: storage compounds.

**Figure S4. Gene expression heatmaps for sulfur oxidation and denitrification including the anoxic condition without sulfide.** (A) Centered expression values of all genes that were differentially expressed between at least two conditions, are shown, with genes in bold that were both differentially expressed between both AS versus O and anoxic without sulfide versus O (twofold change, FDR ≤ 0.05). Genes are ordered by function in the respective metabolic pathways. Note that the expression of many of the genes follows a clear pattern depending on whether oxygen is present or not. (B) Relative elemental sulfur (S^0^) content in ectosymbionts as determined by Raman microspectroscopy after 24 h incubations under anoxic-sulfidic (AS; red dots), anoxic without added sulfide (A, black dots) or in the presence of oxygen (O; dark blue: hypoxic, light blue: oxic). Each dot refers to the value obtained from measuring an individual ectosymbiont cell. 50 cells were measured per condition. Horizontal lines display medians, boxes show the interquartile ranges (25-75%), whiskers indicate minimum and maximum values, and different lower-case letters indicate significant differences among conditions (p < 0.05, Kruskal-Wallis test and Dunn post-hoc test for multiple pairwise comparisons; p (AS vs A) = 3.5E-17, p (AS vs hypoxic) = 0.011, p (AS vs oxic) = 7.4E-13, p (A vs hypoxic) = 3.2E-09, p (A vs oxic) = 0.188, p (hypoxic vs oxic) = 3.6e-06). Relative intensities below 1 (grey dashed line), indicate that elemental sulfur could not be detected. Percentage of cells with sulfur detected: 92 % (AS), 14% (A), 76% (hypoxic), and 22% (oxic).

**Figure S5. Expression of genes involved in energy conservation other than sulfur oxidation and denitrification.** Only genes that were differentially expressed between AS and O are shown (twofold change, FDR ≤ 0.05). Genes are ordered by function. Glycerol-3P-DH: glycerol-3-phosphate dehydrogenase, Fumarate red.: fumarate reductase, Polysulfide/thiosulfate red.: Polysulfide/thiosulfate reductase.

**Figure S6. Lipid composition of ectosymbionts after incubation of symbiotic nematodes in anoxic and oxic conditions.** (A) Major lipid classes and their abundance relative to all lipids detected. (B) Relative abundance of significantly changed glycerophospholipids. Lipid class, fatty acid chain length and saturation are depicted on the x-axis. Note that PG is composed of two fatty acids, while lyso-phospholipids (LPG, LPE and LPC) only contain one fatty acid. Bars show mean abundances relative to total lipids (%) and their standard deviations derived from three analytical replicates. The number of asterisks refers to the significance level (Student’s t-test; * P < 0.05, ** P < 0.01, *** P < 0.001). Note that *Ca*. T. oneisti does not encode for any known phosphatidylcholine biosynthesis genes. PG: phosphatidylglycerol, LPG: lyso-phosphatidylglycerol, LPE: lyso-phosphatidylethanolamine, LPC: lyso-phosphatidylcholine. For details on methodology see Supplementary Materials & Methods.

**Figure S7. NanoSIMS analysis of ^13^C isotope incorporation in *L. oneistus* and its ectosymbiont after incubation in ^13^C-labeled bicarbonate for 24 h under anoxic conditions without sulfide.** The ^13^C content is displayed as ^13^C/(^12^C + ^13^C) isotope fraction, given in at%. (A) NanoSIMS images showing cellular ultrastructure, as displayed by the ^12^C^14^N-secondary ion signal intensity (left), and isotope label distribution (right) in cross sections of *L. oneistus* after incubation of living worms in isotopically labeled (^13^C-live, top row) and unlabeled (^12^C-live, bottom row) bicarbonate. Incubation of 2% PFA-fixed worms under identical conditions in isotopically labeled bicarbonate (^13^C-dead, central row) served as a control for exclusion of unspecific (non-metabolic) label uptake. Scale bars: 5 μm. (B) Region of interest (ROI)-specific evaluation of the isotopic label content, revealing significant ^13^C enrichment both in the ectosymbiont cells and, within particular regions, also in the host tissue. For details on methodology see Supplementary Materials & Methods.

**Figure S8. Phylogenetic tree of the large subunit protein of the ribulose-1,5-bisphosphate carboxylase/oxygenase (RuBisCO).** Unrooted phylogenetic tree illustrating the four forms of RuBisCO from diverse organisms, such as plants, free-living and symbiotic bacteria. The CbbL protein of *Ca*. T. oneisti is highlighted in red. Type I (IA, IB, IC, ID): CbbL, type II: CbbM, type III, type IV: RuBisCO-like. The analysis is based on a MAFFT alignment of full-length amino acid sequences (Table S5) and was estimated under the LG+I+G4 model using Maximum Likelihood phylogeny (IQ-TREE) with node support calculated by SH-aLRT. The scale bar represents 0.5% estimated sequence divergence. SH-aLRT values at the nodes are based on 10 000 replicates. For details on methodology see Supplementary Materials & Methods.

**Figure S9. Differentially expressed genes involved in biosynthesis and utilization of storage compounds and detection of PHA via Raman spectroscopy.** (A) Only differentially expressed genes involved in PHA and trehalose metabolism are shown (2-fold change, FDR ≤ 0.05). (B) Relative PHA content measurement of 50 ectosymbiont cells per condition after 24 h of incubation analyzed by Raman spectroscopy. Horizontal lines indicate medians, boxes show interquartile ranges (25-75%), and whiskers denote minimum and maximum measurements. Each dot represents a single ectosymbiont cell. Cells incubated in the presence of oxygen are depicted in one column (O) but using different colors to indicate the two different oxygenated conditions (light blue: oxic, dark blue: hypoxic). Measurements represented by red dots were obtained from anoxic-sulfidic (AS) incubations. PHA was detected in all conditions, although more PHA was detected in cells incubated in hypoxic incubations (p < 0.05, Kruskal Wallis test, pairwise comparisons; p (AS vs hypoxic) = 0.022, p (AS vs oxic) = 0.057, p (hypoxic vs oxic) = 0.054.

### Supplemental Tables

**Table S1.** Sediment core nematode counts and chemical measurements

**Table S2.** RNA-Seq and EA-IRMS incubation measurements. Note that RNA-Seq and EA-IRMS incubations were performed separately. Samples 1-3 were collected in July 2017, whereas samples 4-6 in March 2019.

**Table S3.** Functional enrichments of selected gene sets. Statistical enrichment of functional categories was tested for GO terms (GO), Pfam domains (PF), KEGG metabolic maps (map), COG category (COG) and COG general category (uppercase letter) using the Bioconductor software package GOseq (M. D. Young, M. J. Wakefield, G. K. Smyth, and A. Oshlack, Genome Biol 11(2): R14, 2010, https://doi.org/10.1186/gb-2010-11-2-r14). Functional categories among sets of protein-coding genes were significantly enriched if the adjusted P value (false discovery rate, FDR) was ≤ 0.1. Only FDR values below that threshold are shown; non-significant FDR values are indicated (NS). AS: anoxic-sulfidic; O: hypoxic + oxic.

**Table S4.** Top expressed *Ca*. T. oneisti proteins. Symbiont proteins under each condition were ranked by relative abundance (%cOrgNSAF), and the 30 proteins with functional annotation exhibiting the highest relative abundance per condition are shown. Proteins written in bold were only found in the top expressed proteins under the respective condition. Color gradients indicate relative abundance and are scaled between values 5.76 and 0.39. Relative abundance of symbiont proteins under two different conditions was determined as described in Supplementary Materials & Methods and is displayed as mean values of three individual replicates per condition. Data S1 specifies all proteins that were detected to be expressed (column “Proteome detection”).

**Table S5.** List of all ribulose-1,5-bisphosphate carboxylase (RuBisCO) protein sequences used for constructing the phylogenetic tree in Figure S8.

### Supplemental Dataset

**Data S1.** *Ca*. T. oneisti genes, functional annotations and expression.

