## Supplemental Data + Text for "Anaerobic sulfur oxidation underlies adaptation of a chemosynthetic symbiont to oxic-anoxic interfaces"

**Supplementary Materials & Methods**

Paredes et al.

**Sediment cores analysis.** To determine the habitat and spatial distribution of *L. oneistus*, we used cores of 60 cm length and 60 mm diameter (UWITEC, Mondsee, Austria) connected to rhizon samplers of a diameter of 2.5 mm and mean pore size of 0.15 µm (Rhizosphere Research Products, Wageningen, Netherlands). This set up allowed the collection of sand and interstitial pore water (the nematode habitat) down to a depth of 30 cm. In total, nine sediment cores were collected in July 2017 at ~1 m depth from a sand bar off Carrie Bow Caye, Belize (16°48′11.01″N, 88°4′54.42″W).

Immediately after collection, the pore-water sulfide content ($\sum H_2S$, i.e. the sum of $H_2S$, $HS^-$ and $S^{2-}$) was determined by the methylene-blue method [1]. In short, 670 µl of a 2% zinc acetate solution was mixed with 335 µl sample and subsequently 335 µl 0.5% *N*,*N*-dimethyl-*p*-phenylenediamine and 17 µl of 10% ferrous ammonium sulfate were added and incubated for 30 min in the dark. Surface seawater was used as a blank. Absorbance was measured at 670 nm and concentrations were quantified via calibration (measurement of $\sum H_2S$ standard solutions in the concentration range from 0 to 0.5 mM $\sum H_2S$). Samples for dissolved inorganic nitrogen (DIN: nitrate, nitrite, and ammonia) and dissolved organic carbon measurements (DOC) were stored and transported deep-frozen, and analyzed at the University of Vienna, Austria. Nitrate ($NO_3^-$) and nitrite ($NO_2^-$) concentrations were determined according to the Griess method [2] using $VCl_3$ [3], whereas the concentration of ammonium ($NH_4^+$) was measured according to Solórzano [4]. For the quantification of nitrate, nitrite, and ammonia, freshly prepared $KNO_3$, $NaNO_2$ and $NH_4Cl$ solutions ranging from 0 to 100 µM were used to create standard curves, respectively. Artificial seawater served as a blank (prepared according to [5]) and all measurements were performed in technical triplicates.

DOC was measured using a Shimadzu TOC-LCPH analyzer equipped with an ASI-L autosampler. After first acidifying the sample (pH 2 to 3) with hydrochloric acid, synthetic air (carbon dioxide free gas) was bubbled for 90 seconds through the sample to eliminate the inorganic carbon component. Next, the remaining total organic carbon was determined. Thereupon, 100 µL sample were injected into the combustion tube, which was filled with an oxidation platinum standard catalyst and heated to 720°C. The resulting combustion products were subsequently dehydrated, cooled and cleaned from chlorine and other halogens. Carbon dioxide was finally detected on a non-dispersive infrared (NDIR) gas analyzer. Each measurement constituted the mean from three 100 µL sample injections.

To determine the abundance of *L. oneistus*, the sand core was subdivided into 6 cm-thick layers and nematode were extracted from each sand layer by stirring the sand in seawater and pouring the supernatant through a 212 µm-mesh sieve. The retained material was transferred into a Petri dish, and single nematodes were handpicked using pipettes under a dissecting microscope. The number of *L. oneistus* nematodes and average $\sum H_2S$, nitrate and nitrite concentrations are shown in Figure S1A. All measurement data are listed in Table S1.

**Nanometer scale secondary ion mass spectrometry (NanoSIMS).** NanoSIMS analysis was performed to visualize and quantify the distribution and incorporation of the $^{13}C$ label into ectosymbiont and host biomass incubated in anoxic conditions without supplemented sulfide. The experimental set up of the incubations was identical with the incubations for EA-IRMS bulk analysis (see main Material and Methods), with the difference that here, we utilized batches of 30 worms in duplicates, and one replicate of 50 worms per incubation was used for EA-IRMS to verify the incorporation of the $^{13}C$ isotope prior to TEM/NanoSIMS sample preparation. EA-IRMS measurement values (δ13C) for the $^{13}C$-live, $^{13}C$-dead and $^{12}C$-live incubations were 403.9, -3.71, and -15.2 ‰, respectively. At the end of each incubation (24 h), the symbiotic nematodes were fixed and stored in 0.1 M Trump's fixative solution (0.1 M sodium cacodylate buffer, 2.5% glutaraldehyde, 2% paraformaldehyde, pH 7.2, 1 000 mOsm $L^{-1}$; [6]) at 4°C until further processing.

To obtain simultaneous information on the isotopic distribution and the site of incorporation, consecutive resin sections for TEM/NanoSIMS analysis were prepared as follows: the fixed samples were washed three times with sodium cacodylate buffer (0.1 M, pH 7.2, 1 000 mOsm $L^{-1}$), each for 10 min at room temperature (RT). Subsequently, the washing buffer was removed, and the samples were incubated in a solution of 1% osmium tetroxide for 1.5 h at RT in a shaker of low speed. Afterwards, the samples were rinsed two times with milli-Q water, each for 10 min at RT, and dehydrated stepwise by application of a concentration series of ethanol. The series consisted of 10 min incubations in 30%, 50%, 70% and 90% ethanol completed by three times 5 min incubations in 100% ethanol. Subsequently, ethanol was substituted by acetone via three times 10 min incubations in 100% acetone. Simultaneously, a fresh mixture of low viscosity resin was prepared (for 100 ml: 48 g LV resin, 8 g VH1 hardener, 44 g VH2 hardener, 2.5 g accelerator; Electron Microscopy Science). The dehydrated samples were then infiltrated stepwise by application of a resin/acetone concentration series: (i) 1:2 resin:acetone mixture for 15 min, (ii) 1:1 resin:acetone mixture for 30 min, (iii) 2:1 resin:acetone mixture for 2 h 30 min, and (iv) 100% resin for 1 h. The final step was conducted inside a vacuum desiccator. Samples were then polymerized in a laboratory oven at 60°C for 48 h. From the obtained resin blocks, thick sections (1-2 µm) were cut by a Leica Ultracut UCT microtome to assess the quality of the embedded samples and to identify appropriate regions for TEM/NanoSIMS analysis. Subsequently, consecutive sections of 70 nm (ultra-thin) and 120 nm (semi-thin) thickness were prepared using a Leica Ultracut UCT microtome and equipped with a diamond knife (Diatome, Bern, Switzerland). The ultra-thin sections (for TEM) were deposited onto previously coated (0.5% formvar solution) slot grids, and stained with 2.5% gadolinium acetate for 25 min, followed by staining with 3% lead citrate for 8 min. After each staining step, the samples were cleaned by gently dipping into milli-Q water for ten times. TEM imaging was conducted on a Zeiss Libra 120 transmission electron microscope (Carl Zeiss

AG, Oberkochen, Germany). The semi-thin sections (for NanoSIMS) were deposited onto antimony-doped silicon wafer platelets (7.1 x 7.1 x 0.7 mm; Active Business Company, Brunnthal, Germany) and analyzed on a NS 50L instrument (Cameca, Gennevilliers, France).

NanoSIMS data were recorded as multilayer image stacks by sequential scanning of a finely focused $Cs^+$ primary ion beam (approx. 80 nm probe size at 2 pA beam current) and simultaneous detection of negative secondary ions and secondary electrons. Recorded images had a 512 × 512 pixel resolution and a field-of-view ranging from 30 × 30 to 60 × 60 $\mu m^2$. The mass spectrometer was tuned for a achieving a mass resolving power of > 10 000 at mass 26 to separate $^{12}C^{14}N^-$ secondary ions from the isobaric species $^{13}C_2^-$. Prior to data acquisition, anaysis areas were pre-conditioned *in situ* by rastering of a high intensity, defocused $Cs^+$ ion beam in the following sequence of high and extreme low ion impact energies (HE / 16 keV and EXLIE / 50 eV, respectively): HE at 100 pA beam current to a fluence of 5.0E14 ions/$cm^2$; EXLIE at 400 pA beam current to a fluence of 5.0E16 ions/$cm^2$; HE at 100 pA to a fluence of 2.5E14 ions/$cm^2$. All images were recorded at a dwell time of 7.5 – 15 ms/pixel/cycle. Secondary ion signal intensities were corrected for detector dead time and quasi-simultaneous arrival (QSA) of secondary ions, using QSA sensitivity factors (“beta” values) of 1.10 for $C^-$ and 1.05 for $CN^-$ ions. Image data were evaluated using the WinImage software package v2.0.8 provided by Cameca. The carbon isotope composition is displayed as $^{13}C/(^{12}C + ^{13}C)$ isotope fraction, given in at%, calculated from $C_2^-$ secondary ion signal intensities via $^{13}C/(^{12}C + ^{13}C) = ^{12}C^{13}C^-/(2*^{12}C^{12}C^- + ^{12}C^{13}C^-)$. Numerical data evaluation was performed on manually defined regions of interest (ROI) (Figure S7B). Individual ROI values from samples of the $^{13}C$-live incubations were considered significantly enriched in $^{13}C$ if (i) the $^{13}C$ isotope fraction was above the 95th percent confidence interval of the corresponding ROI values determined on the negative control samples (i.e. $^{12}C$-live and $^{13}C$-dead: natural isotope abundance control and dead control, respectively) and (ii) the statistical counting error (5σ, Poisson) was smaller than the difference between the considered ROI and the mean value measured on each control.

**Preparation of *Ca.* T. oneisti pellets for proteomics.** 500 symbiotic *Laxus oneistus* were extracted from the sand as described in the main Materials & Methods, and incubated for 24 h in 13 ml of 0.2 µm filtered seawater in exetainers either in the presence of oxygen (mean concentration of dissolved oxygen at incubation start was 195.9 µM, and 183 µM after 24 h) or in anoxic conditions ($O_2$ was detected neither at incubation start, nor after 24 h; no sulfide was added). After the incubations, *Ca.* T. oneisti was dissociated from the nematodes by incubating each batch of 500 nematodes in 2 ml dd$H_2O$ for 1 min, then transferring them to 2 ml 0.2 µM-filtered seawater for 5 min. This osmotic shock causes *Ca.* T. oneisti to detach from the nematodes and move into the seawater, which was collected with a pipette under the dissecting microscope to exclude involuntary aspiration of nematode tissue (or fragments thereof). The 2 ml nematode-free, ectosymbiont suspension was then centrifuged for 1 min at 14 000 x g to obtain *Ca.* T. oneisti pellets. Ectosymbiont pellets and aposymbiotic nematodes were flash-frozen in liquid nitrogen and stored at -80°C until further processing. Only *Ca*. T. oneisti proteomic data are shown in this study. *L. oneistus* proteomics will be published separately.

**Protein extraction and 1D PAGE.** *Ca.* T. oneisti proteins were extracted as described previously [7]. Briefly, both samples, i.e. frozen ectosymbiont cell pellets from oxic and anoxic incubations, were resuspended in 1% (w/v) sodium deoxycholate (SDC), 4% (w/v) sodium dodecyl sulfate (SDS) in 50 mM triethylammonium bicarbonate buffer (lysis buffer). After boiling the samples for 5 min under agitation (600 rpm), they were incubated in an ultrasonic bath for 5 min at RT. After removal of cell debris by a 10 min centrifugation at RT (14 000 x g), protein concentrations in the supernatants were determined using the Pierce BCA (bicinchoninic acid) assay (Thermo Scientific Pierce) according to the manufacturer's instructions in a Tecan microtiter plate reader. For gel-based proteomic analysis (as previously described by [8]), 25 µg of protein per sample were mixed with loading buffer (2 % (w/v) SDS, 10 % glycerol, 12.5 mM dithiothreitol, 0.001 % (w/v) bromophenol blue in 0.06 M Tris-HCl) and separated in precast 4 – 20 % SDS mini gels (BioRad TGX). Per sample, three replicates (3 x 25 µg protein) were separated (giving a total of 6 samples). After staining with Coomassie Brilliant Blue, protein-containing gel lanes were excised and subdivided into 10 equal-sized pieces each, which were destained at 37 °C in 200 mM $NH_4HCO_3$ 30 % acetonitrile under agitation at 600 rpm and digested overnight at 37 °C with trypsin (sequencing grade, Promega). Finally, peptides were eluted in an ultrasonic bath and subjected to LC-MS/MS analysis.

**LC-MS/MS analysis.** Peptides were analyzed by reversed phase liquid chromatography (LC) electrospray ionization (ESI) MS/MS using an LTQ Orbitrap Velos (Thermo Fisher Scientific) according to [9]. Briefly, in-house self-packed nano-LC columns (100 µm x 20 cm) containing reverse-phase C18 material (3 µm, ReproSil-Pur 120-AQ; Dr. Maisch GmbH, Ammerbuch-Entringen, Germany) were used to perform LC with an Easy-nLC1000 system (Thermo Fisher Scientific). The peptides were loaded with solvent A (0.1% acetic acid (v/v)). Subsequently, the peptides were eluted by a non-linear binary gradient of 80 minutes from 5% to 99% solvent B (0.1% acetic acid (v/v), 99.9% acetonitrile (v/v)) in solvent A at a constant flow rate of 300 nl/min. MS data were acquired in data-dependent MS/MS mode for the 20 most abundant precursor ions. After a full scan in the Orbitrap (*m/z* 300 – 1 700) with a resolution of 30 000 at *m/z* 400, ions were fragmented via collision-induced dissociation (CID) and recorded in the linear trap quadrupole LTQ analyzer.

**Protein identification and quantification***.* For protein identification, a database was constructed, containing 18 364 *Laxus oneistus* host protein sequences (derived from a *de novo* assembled transcriptome; will be published separately), 5 169 *Ca*. T. oneisti protein sequences (JAAEFD000000000, see main Materials and Methods) and a set of 42 common laboratory contaminants. All sequences were reversed and appended to the database as decoys to allow for false-discovery rate (FDR) assessment. Mass spectra were searched against this target-decoy database using the Sorcerer SEQUEST algorithm (Sage-N Research) and filtered using Scaffold (version 4.8.4, http://www.proteomesoftware.com) applying the following thresholds: i) protein FDR and peptide FDR were set to 1% and ii) at least two unique peptides were required for a protein or protein group to be identified. Proteins were expressed if they were detected in at least two out of the three replicates in at least one condition. This way, 1 137 ectosymbiont proteins (22.0% of all predicted proteins in the database) were identified in total. Data S1 indicates all detected proteins in the column "Proteome detection". Relative abundance of identified proteins was calculated from total spectrum counts as normalized spectral abundance factor (%NSAF) values – giving the percentage of each protein relative to all proteins in the respective sample [10], and as %OrgNSAF, giving a protein's percentage relative to all ectosymbiont proteins in the respective sample [11]. As ectosymbiont protein identification rates varied substantially between oxic and anoxic samples, which may negatively affect comparability of relative abundances between samples, we included only such proteins in the final quantitation, which were detected under both conditions (824 proteins). This additional normalization step provided corrected %OrgNSAF values (%cOrgNSAF), which give a protein's percentage relative to all symbiont proteins that were expressed under both conditions.

**Intact polar lipid extraction and analysis.** Five batches of 100 freshly collected *Laxus oneistus* were incubated for 24 h in oxic or anoxic (no sulfide added) conditions as described in the main Materials and Methods (RNA-seq incubations). At the beginning of the incubations, mean concentrations of dissolved oxygen in the 0.2 µm filtered seawater were 180.9 µM (oxic) and 0.47 µM (anoxic). After 24 h, we measured on average 86 µM (oxic) and 0 µM (anoxic) oxygen, respectively. Sulfide ($\sum H_2S$) could not be detected in any of the incubations. At the end of the incubations, *Ca*. T. oneisti (from either the oxic or anoxic conditions) were dissociated from 500 nematodes, as described above (Proteomics). Symbiont pellets were flash-frozen in liquid nitrogen and stored at -80°C until further processing.

Lipids of the ectosymbionts were extracted using a modified Folch extraction [12] previously applied for lipid extraction from bacteria [13]. Briefly, pelleted bacteria were taken up in 1.6 ml 0.2 µm filtered seawater and 0.5 ml were transferred to 2 ml glass vials obtaining three analytical replicates. Bacteria were then pelleted by centrifugation, resuspended in 0.5 ml methanol and extracted using chloroform-methanol (all solvents LC-MS grade, Sigma-Aldrich). Extracted lipids were dried under nitrogen gas on a Techne Sample Concentrator and re-suspended in 1 ml of acetonitrile: 10 mM ammonium acetate at a 95:5 (v:v) ratio. Samples were analyzed by liquid chromatography mass spectrometry (LC-MS) as follows: lipids were separated on a Dionex UltiMate 3000RS UHPLC (Thermo Fisher Scientific) equipped with a XBridge BEH amide XP column (Waters, Milford, MA, USA) and coupled to an amaZon SL quadrupole ion trap MS (Bruker, Billerica, MA, USA) for detection. The column was maintained at 30°C with a flow rate of 150 µl $min^{-1}$. Samples were separated by a 15 min gradient from 95% (v:v) acetonitrile (Solvent A) to 30% (w:v) 10 mM ammonium acetate (pH 9.2, Solvent B) with 10 min equilibration between samples. Sample analysis was carried out in both positive and negative ion mode and fragmentation performed by the $autoMS^n$ function in Compass HyStar (Bruker). We used the Bruker Compass software package for lipid data analysis: DataAnalysis for peak detection and lipid identification, and QuantAnalysis for quantification of the relative abundances of lipids. Peak integration was manually corrected where necessary. Consecutively, for data normalization the peak area of each lipid was divided by the sum of the peak areas of all detected lipids in each sample. Statistical analysis of significant differences in ectosymbiont lipids between the anoxic and oxic condition was carried out using a Student's t-test.

**RuBisCO phylogenetic tree**. Amino acid sequences of the RuBisCO forms I-IV (Table S5) were obtained from GenBank and SwissProt databases, and aligned using mafft v7.397 [14]. Misaligned sequences were manually inspected. Gaps in more than 70% of the sequences were removed using TrimAl 1.4.rev15 [15]. The maximum-likelihood tree with SH-aLRT support values (10 000 replicates) was inferred using IQ-TREE v1.6.2 with automatic model selection [16, 17].

**Data availability**. The proteomics raw data and the combined *L. oneistus* host and ectosymbiont database used for proteomic analyses have been deposited to the

ProteomeXchange Consortium via the PRIDE [18] partner repository with the data set identifier PXD017709.

17. Kalyaanamoorthy S, Minh BQ, Wong TKF, Von Haeseler A, Jermiin LS. ModelFinder: Fast model selection for accurate phylogenetic estimates. *Nat Methods*

2017; **14**: 587–589.

18. Vizcaíno JA, Côté RG, Csordas A, Dianes JA, Fabregat A, Foster JM, et al. The Proteomics Identifications (PRIDE) database and associated tools: Status in 2013. *Nucleic Acids Res* 2013; **41**.

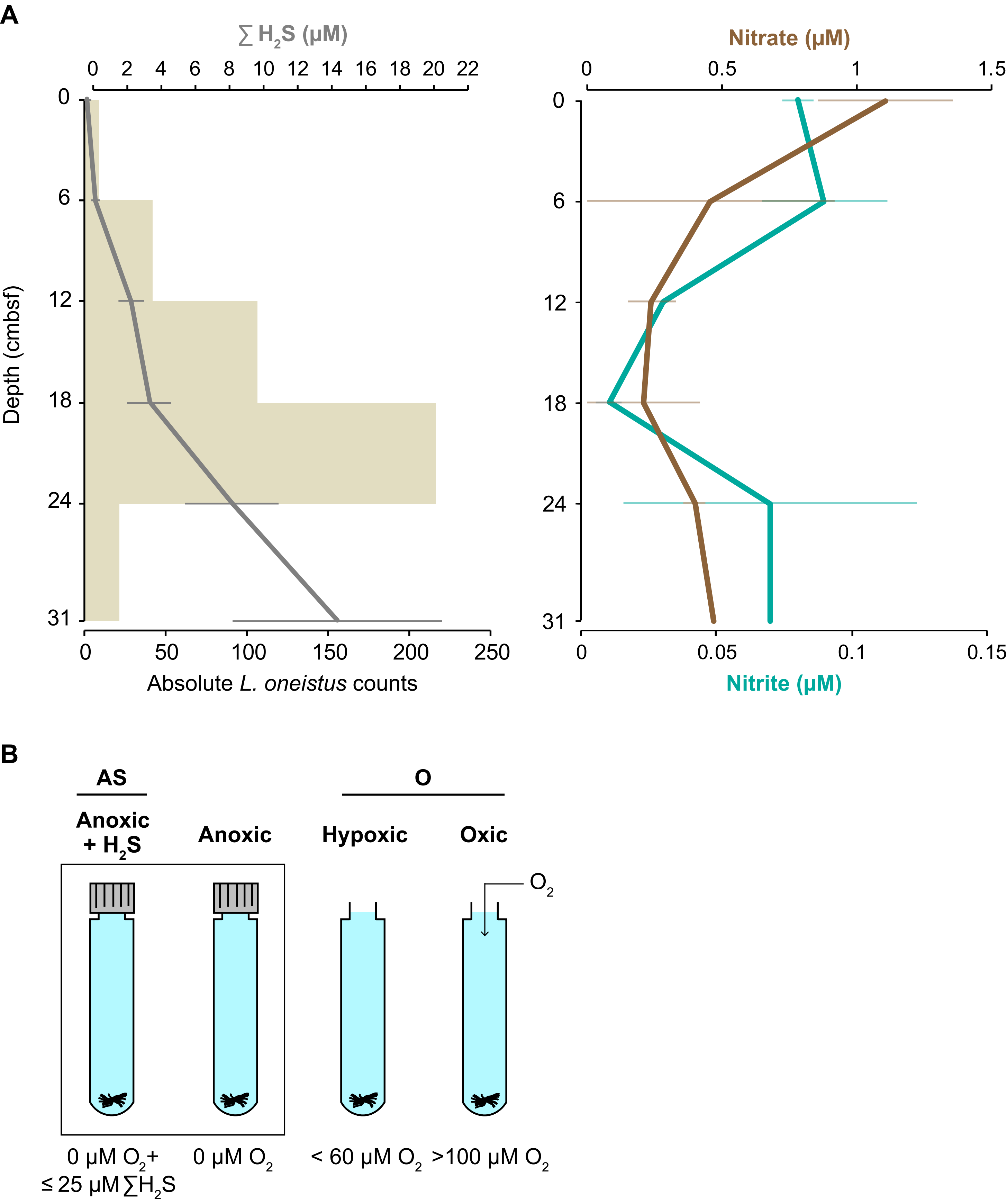
A
∑ $H_2S$ (μM)
0 2 4 6 8 10 12 14 16 18 20 22
Depth (cmbsf)
0
6
12
18
24
31
0 50 100 150 200 250
Absolute *L. oneistus* counts
Nitrate (μM)
0 0.5 1 1.5
0 0.05 0.1 0.15
Nitrite (μM)
B
AS
O
Anoxic + $H_2S$
Anoxic
Hypoxic
Oxic
$O_2$
0 μM $O_2$+ ≤ 25 μM ∑$H_2S$
0 μM $O_2$
< 60 μM $O_2$
>100 μM $O_2$

**Figure S1. Natural versus experimental conditions.** (A) The left panel shows *Laxus oneistus* total counts per 6 cm core subsection from all 8 sandbars (horizontal beige bars) and corresponding mean sulfide ($\sum H_2S$) concentrations (µM, grey line). The right panel shows mean nitrate and nitrate concentrations through depth in the same sandbars of Carrie Bow Cay, Belize. Error bars represent the standard error of the mean. Note that a few nitrate and nitrite data points are derived from only two or one sandbars (Table S1). (B) Experimental set up of incubations for RNA-Seq, EA-IRMS and Raman microspectroscopy. Batches of 50 *L. oneistus* were incubated under different oxygen concentrations: anoxic with sulfide (0 µM $O_2$, ≤ 25 µM sodium sulfide added), anoxic without sulfide (0 µM $O_2$), hypoxic (< 60 µM $O_2$ after 24 h), and oxic (> 100 µM $O_2$ after 24 h). The box around the anoxic incubation vials indicates that these incubations were carried out in a polyethylene isolation chamber. Given the similarity of gene expression profiles between the hypoxic and oxic samples (see Figure S2), most of the follow-up analyses were conducted by treating the hypoxic and oxic samples as biological replicates (O), and comparing the O condition to the anoxic-sulfidic (AS) condition. All incubations were performed in 0.2 µM-filtered seawater and in biological triplicates.

**A**

| Samples | Total number of reads | Number of reads mapped to symbiont genome | Number of reads mapped to symbiont genes | % reads mapped to symbiont genome | % reads mapped to symbiont genes | % reads mapped to rRNA genes |
|---|---|---|---|---|---|---|
| oxic-1 | 39,788,652 | 935,582 | 571,770 | 1.4 | 61.1 | 2.1 |
| oxic-2 | 35,270,773 | 1,308,826 | 795,913 | 2.3 | 60.8 | 1.6 |
| oxic-3 | 40,645,573 | 1,140,752 | 680,586 | 1.7 | 59.7 | 1.5 |
| oxic-4 | 25,823,639 | 1,422,342 | 786,363 | 3.0 | 55.3 | 1.9 |
| oxic-5 | 36,002,778 | 1,401,873 | 757,557 | 2.1 | 54.0 | 1.9 |
| oxic-6 | 61,503,518 | 2,065,340 | 1,178,912 | 1.9 | 57.1 | 2.5 |
| hypoxic-1 | 37,528,019 | 2,064,723 | 1,328,297 | 3.5 | 64.3 | 1.8 |
| hypoxic-2 | 36,704,147 | 2,246,543 | 1,430,114 | 3.9 | 63.7 | 0.5 |
| hypoxic-3 | 39,087,262 | 868,032 | 555,176 | 1.4 | 64.0 | 1.0 |
| anoxic-1 | 36,848,477 | 1,755,816 | 1,180,245 | 3.2 | 67.2 | 0.7 |
| anoxic-2 | 36,720,206 | 1,767,472 | 1,219,640 | 3.3 | 69.0 | 0.8 |
| anoxic-3 | 37,438,972 | 1,757,143 | 1,153,214 | 3.1 | 65.6 | 1.2 |
| anoxic-4 | 57,013,563 | 1,816,539 | 971,290 | 1.7 | 53.5 | 2.8 |
| anoxic-5 | 41,002,474 | 1,738,244 | 1,133,192 | 2.8 | 65.2 | 2.7 |
| anoxic-sulfidic-1 | 55,241,503 | 508,069 | 341,801 | 0.6 | 67.3 | 2.8 |
| anoxic-sulfidic-2 | 36,482,028 | 418,210 | 281,620 | 0.8 | 67.3 | 2.1 |
| anoxic-sulfidic-3 | 39,792,029 | 349,853 | 234,535 | 0.6 | 67.0 | 2.5 |

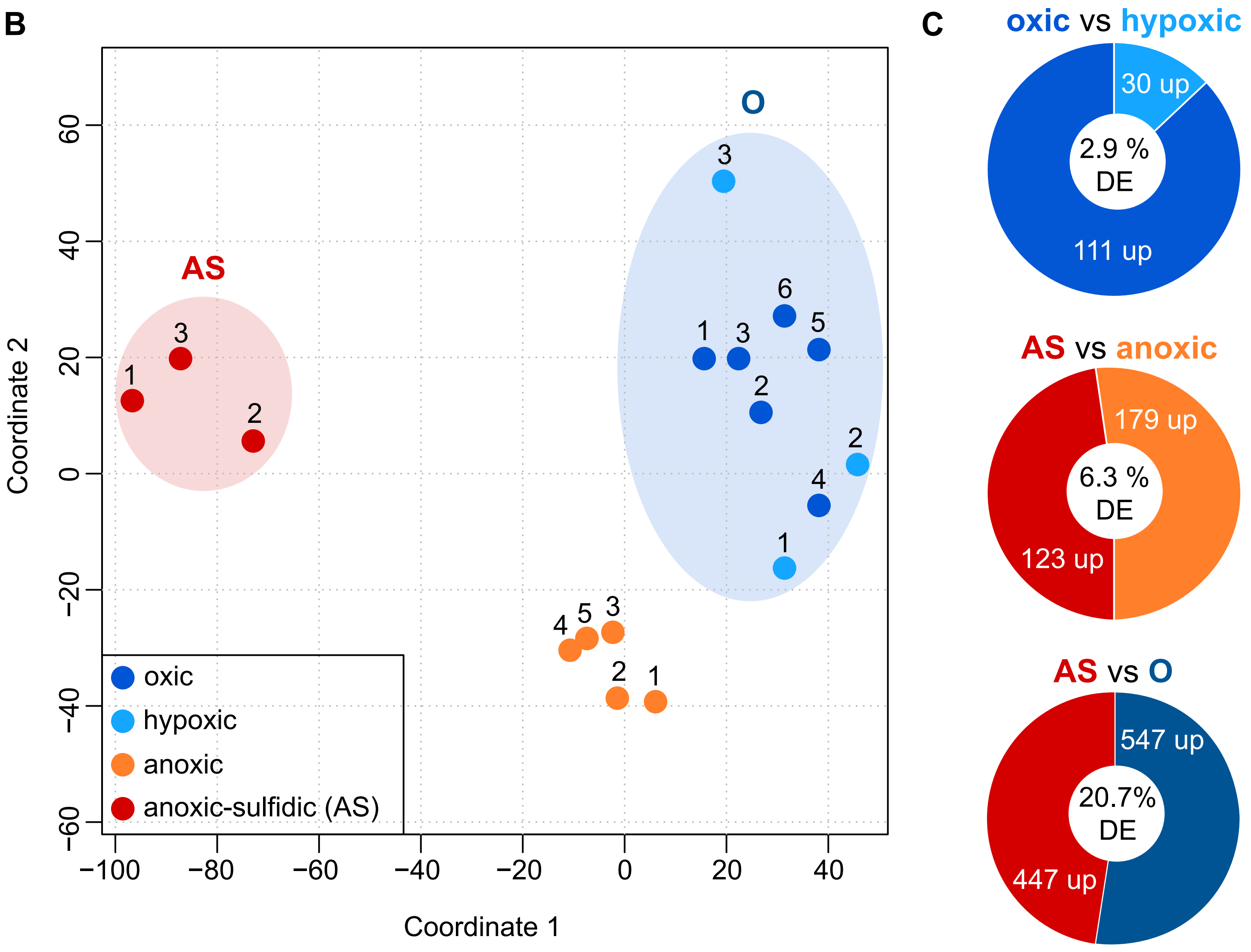

**Figure S2**. **RNA sequencing statistics, sample similarity and differential gene expression.** (A) RNA sequencing and mapping statistics. Sequencing reads were mapped to the symbiont genome assembly consisting of 401 contigs and 5 169 protein-coding genes. The total number of reads refers to the number of reads after quality filtering and trimming, and the number of reads mapped to the genome (i.e. genes, intergenic regions and antisense regions) only includes uniquely mapped reads. (B) Similarity between samples based on Euclidean distances between expression values ($log_2$TPMs), visualized by means of multidimensional scaling (MDS). A total of 4 797 protein-coding genes (92.8%) were detected to be expressed. Most of the follow-up analyses were conducted comparing the anoxic-sulfidic conditions (AS, red circle) to conditions in which oxygen was present (O, blue circle). (C) Differential expression (DE) analysis between hypoxic and oxic samples revealed that the number of DE genes was low (2.9% of all expressed genes), and thus hypoxic and oxic samples were treated as biological replicates. 20.7% of all expressed genes were differentially expressed between AS and O conditions. Genes were considered differentially expressed if their expression changed twofold with a false-discovery rate (FDR) $\leq 0.05$.

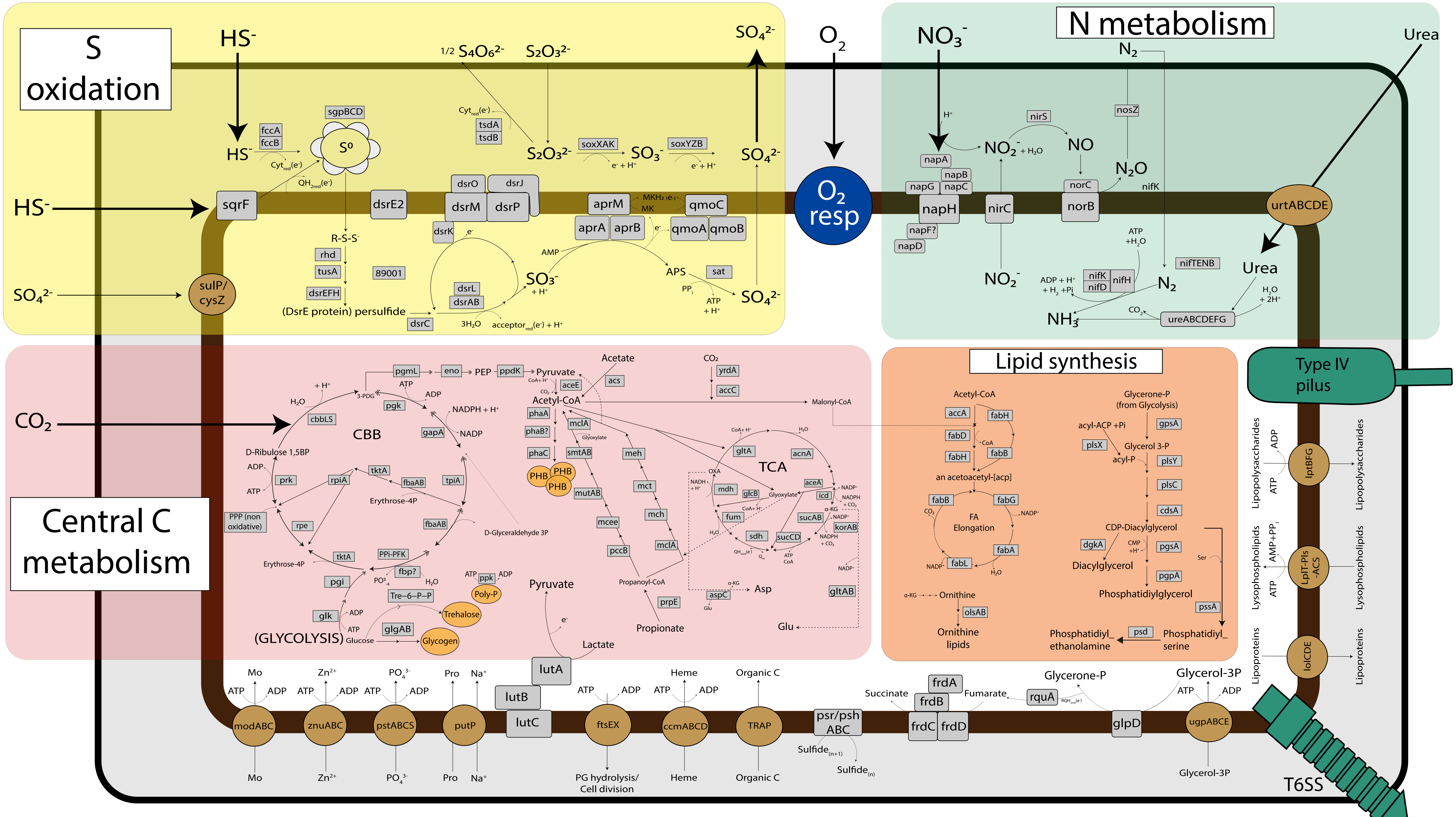


**Figure S3. Schematic representation of central metabolic pathways present in the *Ca.* T. oneisti genome.** All gene names are indicated in Data S1. Organic carbon compounds such as acetate, lactate, propionate and glycerol 3-phosphate (glycerol-3P) could be host-derived. The respiratory chain of oxygen respiration ($O_2$ resp., blue) consists of NADH dehydrogenase (*nuo* genes, complex I), succinate dehydrogenase (*sdh* genes, complex II), the cytochrome bc1 complex (*pet* genes, complex III) and an aa3-type cytochrome c oxidase (*cta* genes, complex IV). Note that *Ca.* T. oneisti only encodes a single complex IV enzyme. Grey: enzymes, brown: transporters, orange: storage compounds.

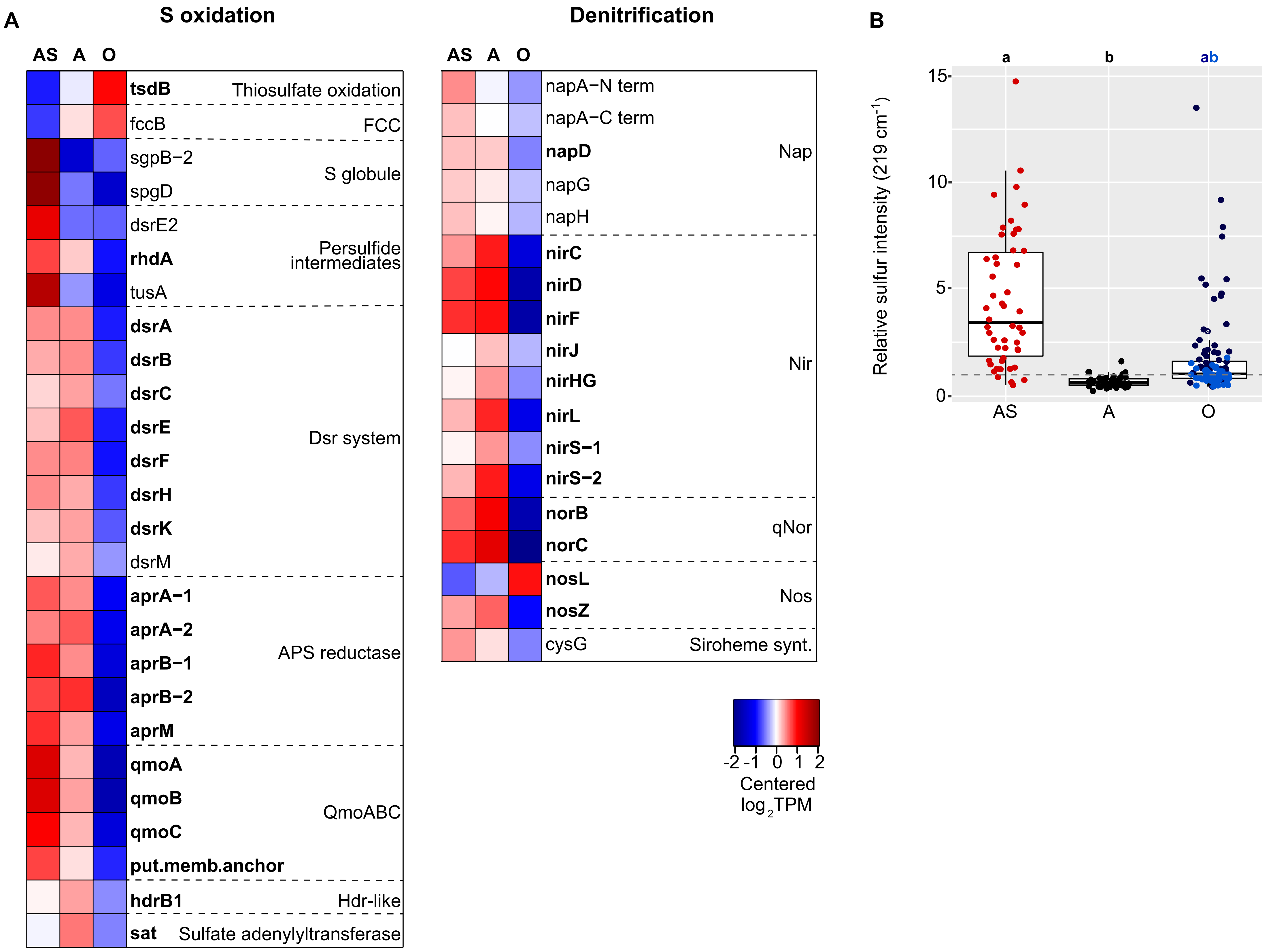

A
S oxidation
AS A O
tsdB Thiosulfate oxidation
fccB FCC
sgpB−2
spgD
S globule
dsrE2
rhdA
tusA
Persulfide intermediates
dsrA
dsrB
dsrC
dsrE
dsrF
dsrH
dsrK
dsrM
Dsr system
aprA−1
aprA−2
aprB−1
aprB−2
aprM
APS reductase
qmoA
qmoB
qmoC
put.memb.anchor
QmoABC
hdrB1 Hdr-like
sat Sulfate adenylyltransferase
Denitrification
AS A O
napA−N term
napA−C term
napD
napG
napH
Nap
nirC
nirD
nirF
nirJ
nirHG
nirL
nirS−1
nirS−2
Nir
norB
norC
qNor
nosL
nosZ
Nos
cysG Siroheme synt.
-2 -1 0 1 2
Centered log2TPM
B
a b ab
Relative sulfur intensity (219 cm-1)
15
10
5
0
AS A O

**Figure S4**. **Gene expression heatmaps for sulfur oxidation and denitrification including the anoxic condition without sulfide.** (A) Centered expression values of all genes that were differentially expressed between at least two conditions, are shown, with genes in bold that were both differentially expressed between both AS versus O and anoxic without sulfide versus O (twofold change, FDR ≤ 0.05). Genes are ordered by function in the respective metabolic pathways. Note that the expression of many of the genes follows a clear pattern depending on whether oxygen is present or not. (B) Relative elemental sulfur ($S^0$) content in ectosymbionts as determined by Raman microspectroscopy after 24 h incubations under anoxic-sulfidic (AS; red dots), anoxic without added sulfide (A, black dots) or in the presence of oxygen (O; dark blue: hypoxic, light blue: oxic). Each dot refers to the value obtained from measuring an individual ectosymbiont cell. 50 cells were measured per condition. Horizontal lines display medians, boxes show the interquartile ranges (25-75%), whiskers indicate minimum and maximum values, and different lower-case letters indicate significant differences among conditions ($p < 0.05$, Kruskal-Wallis test and Dunn post-hoc test for multiple pairwise comparisons; p (AS vs A) = 3.5E-17, p (AS vs hypoxic) = 0.011, p (AS vs oxic) = 7.4E-13, p (A vs hypoxic) = 3.2E-09, p (A vs oxic) = 0.188, p (hypoxic vs oxic) = 3.6e-06). Relative intensities below 1 (grey dashed line), indicate that elemental sulfur could not be detected.  Percentage of cells with sulfur detected: 92 % (AS), 14% (A), 76% (hypoxic), and 22% (oxic).

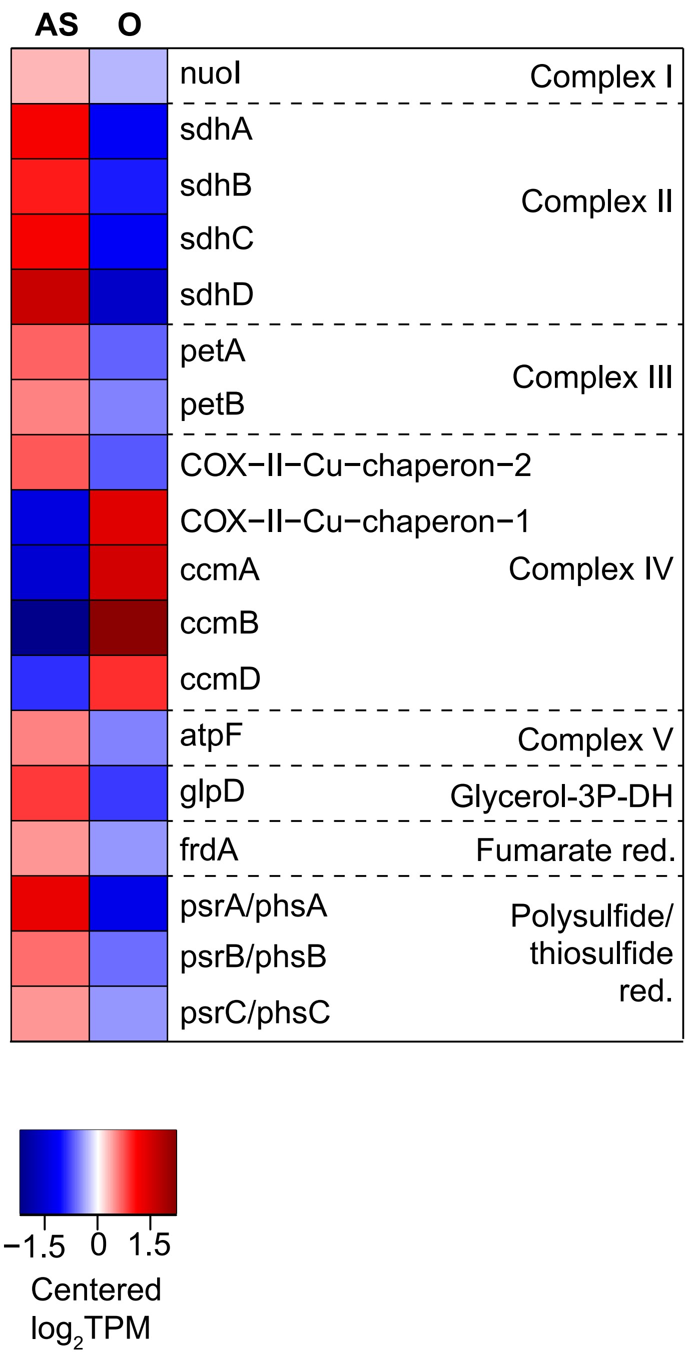


**Figure S5. Expression of genes involved in energy conservation other than sulfur oxidation and denitrification.** Only genes that were differentially expressed between AS and O are shown (twofold change, FDR ≤ 0.05). Genes are ordered by function. Glycerol-3P-DH: glycerol-3-phosphate dehydrogenase, Fumarate red.: fumarate reductase, Polysulfide/thiosulfate red.: Polysulfide/thiosulfate reductase.

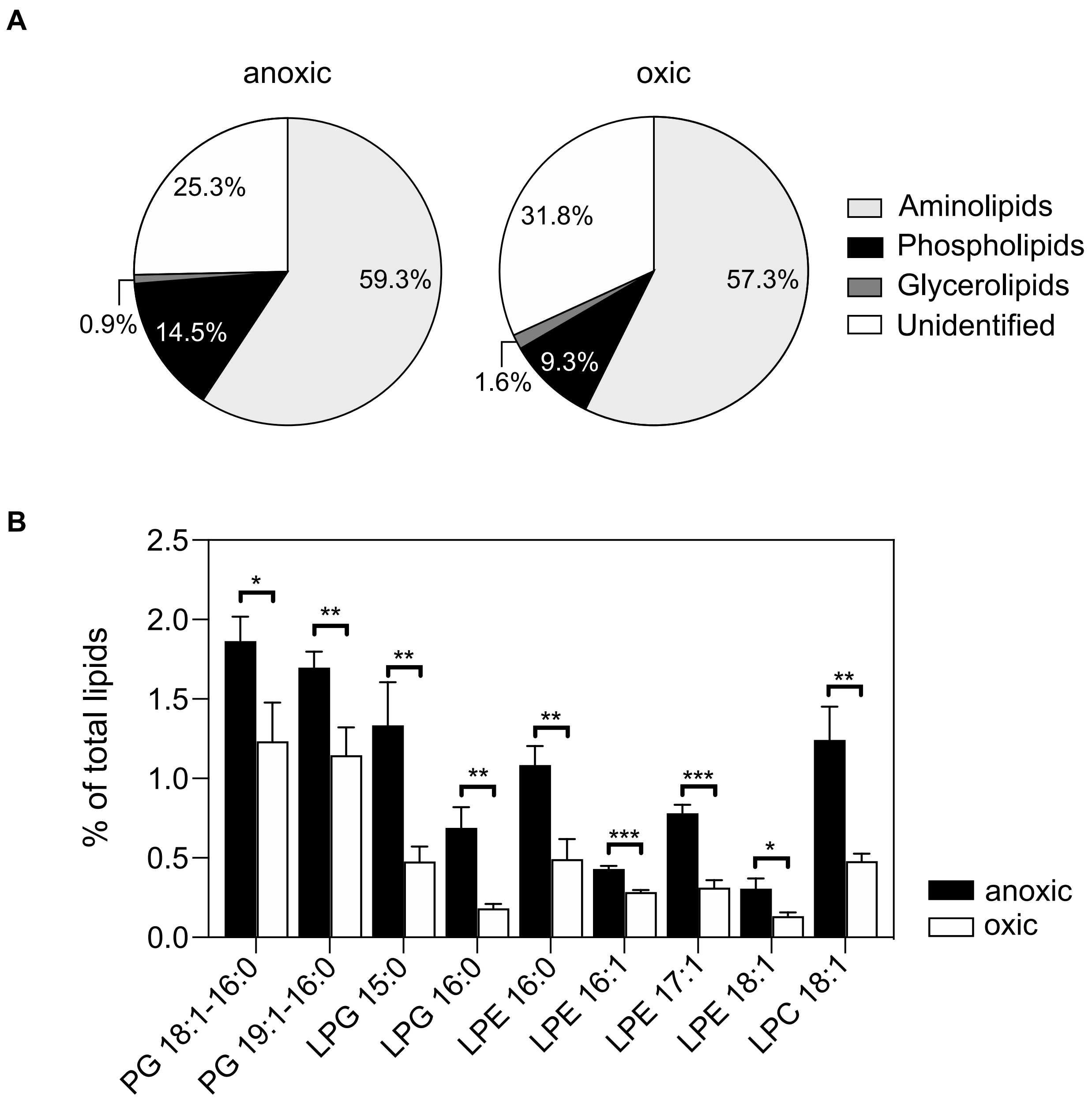


**Figure S6. Lipid composition of ectosymbionts after incubation of symbiotic nematodes in anoxic and oxic conditions.** (A) Major lipid classes and their abundance relative to all lipids detected. (B) Relative abundance of significantly changed glycerophospholipids. Lipid class, fatty acid chain length and saturation are depicted on the x-axis. Note that PG is composed of two fatty acids, while lyso-phospholipids (LPG, LPE and LPC) only contain one fatty acid. Bars show mean abundances relative to total lipids (%) and their standard deviations derived from three analytical replicates. The number of asterisks refers to the significance level (Student's t-test; * $P < 0.05$, ** $P < 0.01$, *** $P < 0.001$). Note that *Ca*. T. oneisti does not encode for any known phosphatidylcholine biosynthesis genes. PG: phosphatidylglycerol, LPG: lyso-phosphatidylglycerol, LPE: lyso-phosphatidylethanolamine, LPC: lyso-phosphatidylcholine. For details on methodology see Supplementary Materials & Methods.

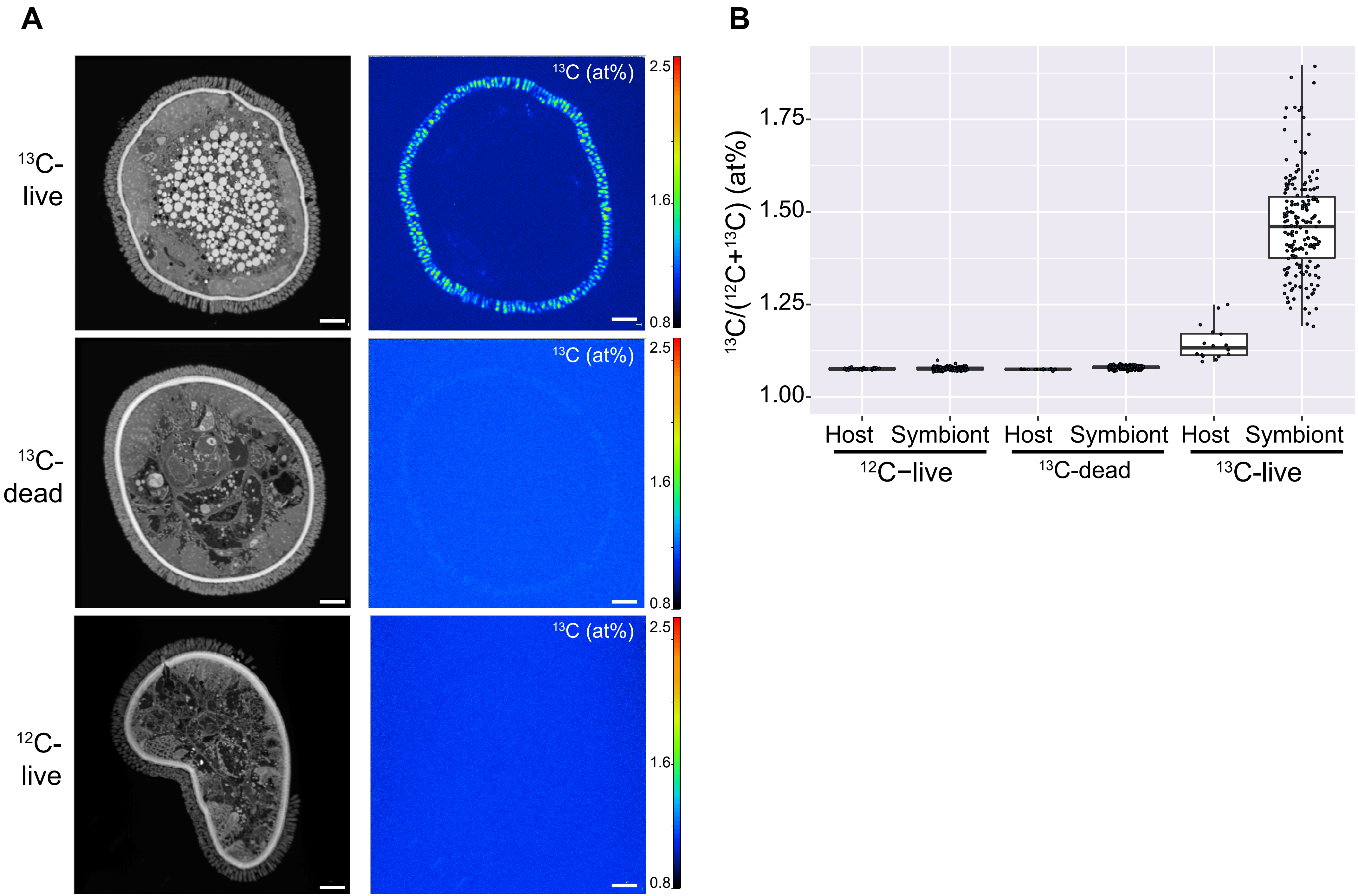
A
$^{13}$C-live
$^{13}$C-dead
$^{12}$C-live
$^{13}$C (at%)
2.5
1.6
0.8
B
$^{13}$C/($^{12}$C+$^{13}$C) (at%)
1.75
1.50
1.25
1.00
Host
Symbiont
$^{12}$C−live
$^{13}$C-dead
$^{13}$C-live

**Figure S7. NanoSIMS analysis of $^{13}C$ isotope incorporation in *L. oneistus* and its ectosymbiont after incubation in $^{13}C$-labeled bicarbonate for 24 h under anoxic conditions without sulfide.** The $^{13}C$ content is displayed as $^{13}C/(^{12}C + ^{13}C)$ isotope fraction, given in at%. (A) NanoSIMS images showing cellular ultrastructure, as displayed by the $^{12}C^{14}N^{-}$ secondary ion signal intensity (left), and isotope label distribution (right) in cross sections of *L. oneistus* after incubation of living worms in isotopically labeled ($^{13}C$-live, top row) and unlabeled ($^{12}C$-live, bottom row) bicarbonate. Incubation of 2% PFA-fixed worms under identical conditions in isotopically labeled bicarbonate ($^{13}C$-dead, central row) served as a control for exclusion of unspecific (non-metabolic) label uptake. Scale bars: 5 µm. (B) Region of interest (ROI)-specific evaluation of the isotopic label content, revealing significant $^{13}C$ enrichment both in the ectosymbiont cells and, within particular regions, also in the host tissue. For details on methodology see Supplementary Materials & Methods.

**Figure S8**

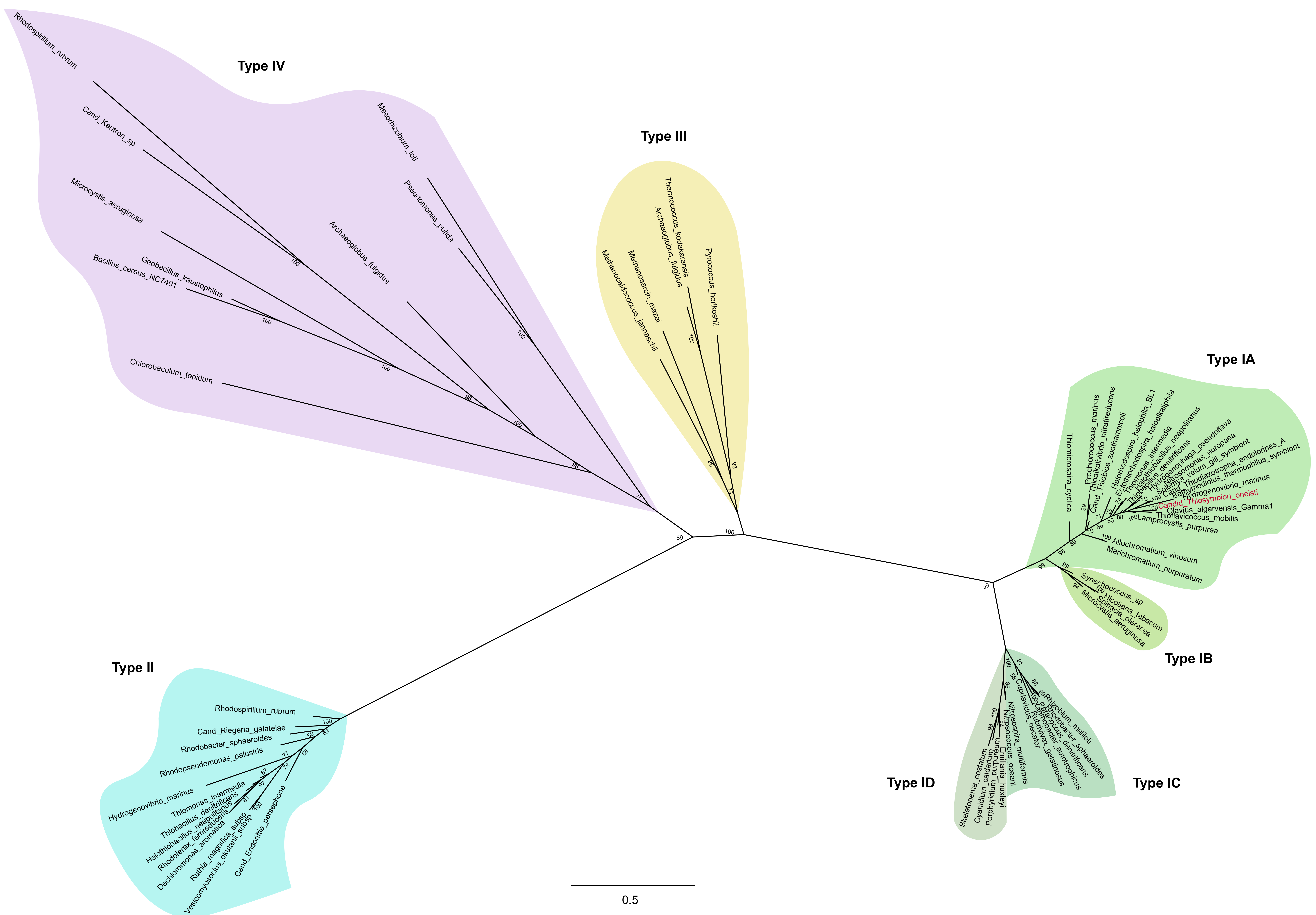
Type IV
Type III
Type IA
Type IB
Type IC
Type ID
Type II
Rhodospirillum_rubrum
Cand_Kentron_sp
Mesorhizobium_loti
Pseudomonas_putida
Microcystis_aeruginosa
Archaeoglobus_fulgidus
Geobacillus_kaustophilus
Bacillus_cereus_NC7401
Chlorobaculum_tepidum
Thermococcus_kodakarensis
Archaeoglobus_fulgidus
Pyrococcus_horikoshii
Methanosarcin_mazei
Methanocaldococcus_jannaschii
Candid_Thiosymbion_oneisti
Olavius_algarvensis_Gamma1
Thioflavicoccus_mobilis
Lamprocystis_purpurea
Allochromatium_vinosum
Marichromatium_purpuratum
Thiomicrospira_cyclica
Synechococcus_sp
Nicotiana_tabacum
Spinacia_oleracea
Microcystis_aeruginosa
Rhizobium_meliloti
Rhodobacter_sphaeroides
Paracoccus_denitrificans
Xanthobacter_autotrophicus
Rubrivivax_gelatinosus
Cupriavidus_necator
Nitrosospira_multiformis
Nitrosococcus_oceani
Skeletonema_costatum
Cyanidium_caldarium
Porphyridium_purpureum
Emiliania_huxleyi
Rhodospirillum_rubrum
Cand_Riegeria_galatelae
Rhodobacter_sphaeroides
Rhodopseudomonas_palustris
Hydrogenovibrio_marinus
Thiomonas_intermedia
Thiobacillus_denitrificans
Halothiobacillus_neapolitanus
Rhodoferax_ferrireducens
Dechloromonas_aromatica
Ruthia_magnifica_subsp
Vesicomyosocius_okutanii_subsp
Cand_Endoriftia_persephone
0.5

**Figure S8**. **Phylogenetic tree of the large subunit protein of the ribulose-1,5-bisphosphate carboxylase/oxygenase (RuBisCO).** Unrooted phylogenetic tree illustrating the four forms of RuBisCO from diverse organisms, such as plants, free-living and symbiotic bacteria. The CbbL protein of *Ca.* T. oneisti is highlighted in red. Type I (IA, IB, IC, ID): CbbL, type II: CbbM, type III, type IV: RuBisCO-like. The analysis is based on a MAFFT alignment of full-length amino acid sequences (Table S5) and was estimated under the LG+I+G4 model using Maximum Likelihood phylogeny (IQ-TREE) with node support calculated by SH-aLRT. The scale bar represents 0.5% estimated sequence divergence. SH-aLRT values at the nodes are based on 10 000 replicates. For details on methodology see Supplementary Materials & Methods.

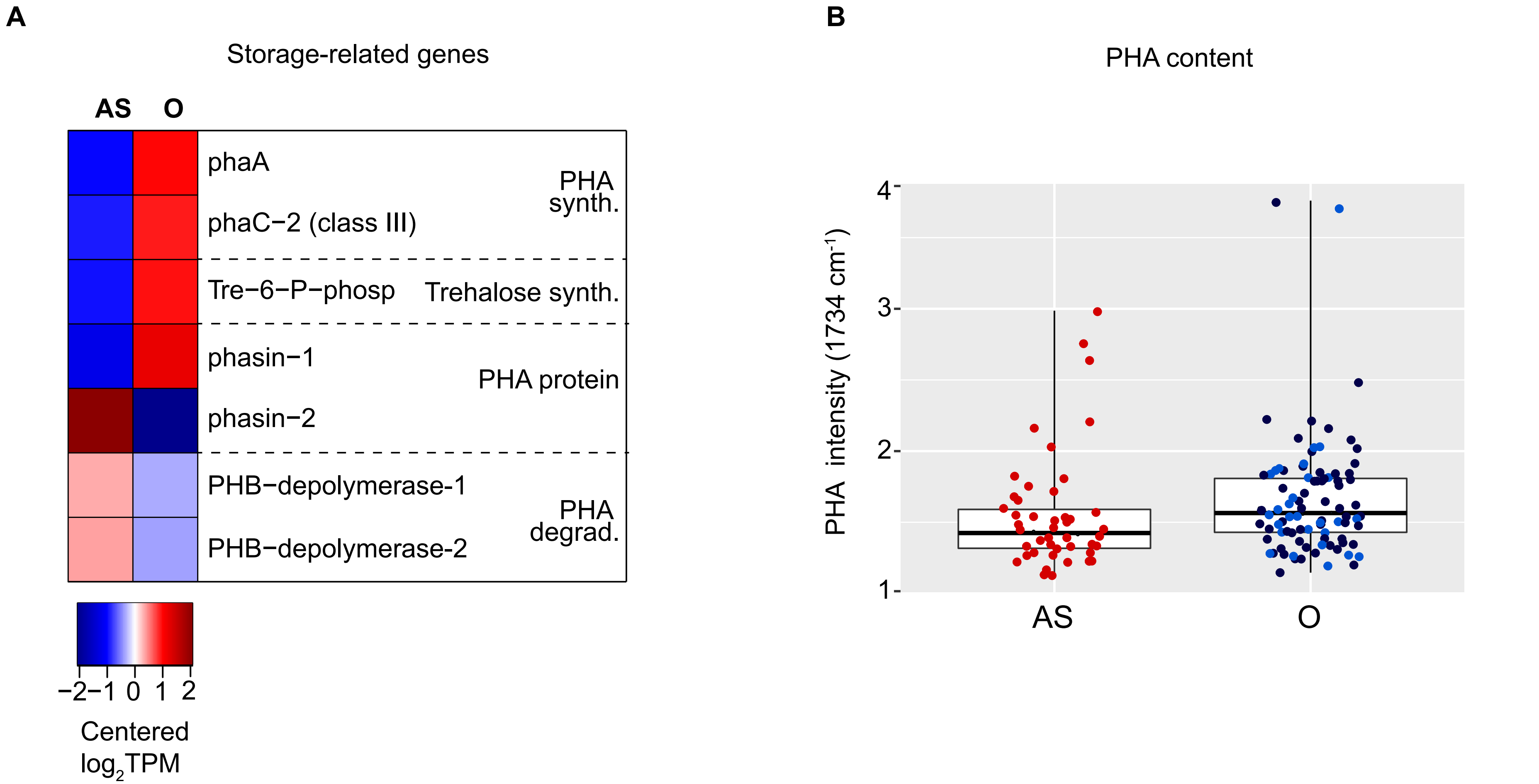


**Figure S9. Differentially expressed genes involved in biosynthesis and utilization of storage compounds and detection of PHA via Raman spectroscopy.** (A) Only differentially expressed genes involved in PHA and trehalose metabolism are shown (2-fold change, FDR ≤ 0.05). (B) Relative PHA content measurement of 50 ectosymbiont cells per condition after 24 h of incubation analyzed by Raman spectroscopy. Horizontal lines indicate medians, boxes show interquartile ranges (25-75%), and whiskers denote minimum and maximum measurements. Each dot represents a single ectosymbiont cell. Cells incubated in the presence of oxygen are depicted in one column (O) but using different colors to indicate the two different oxygenated conditions (light blue: oxic, dark blue: hypoxic). Measurements represented by red dots were obtained from anoxic-sulfidic (AS) incubations. PHA was detected in all conditions, although more PHA was detected in cells incubated in hypoxic incubations ($p < 0.05$, Kruskal Wallis test, pairwise comparisons; p (AS vs hypoxic) = 0.022, p (AS vs oxic) = 0.057, p (hypoxic vs oxic) = 0.054.

**Anaerobic sulfur oxidation underlies adaptation of a chemosynthetic symbiont to oxic-anoxic interfaces**

**Supplementary Tables**

Paredes et al.

**Table S1.** Sediment core nematode counts and chemical measurements

| | Sediment core | | | | | | | | | | |
|---|---|---|---|---|---|---|---|---|---|---|---|
| *L. oneistus* counts | | | | | | | | | | | |
| Depth (cmbsf) | A | B | C | D | E | F | G | H | I | Total | |
| 1 to 7 | 0 | 3 | 1 | 4 | 1 | 0 | 1 | 0 | 0 | 10 | |
| 7 to 13 | 0 | 4 | 2 | 22 | 9 | 1 | 1 | 3 | 0 | 42 | |
| 13 to 19 | 15 | 8 | 6 | 13 | 31 | 6 | 3 | 20 | 5 | 107 | |
| 19 to 25 | 27 | 26 | 2 | 10 | 9 | 21 | 4 | 67 | 50 | 216 | |
| 25 to 31 | ND | ND | 12 | ND | 10 | 0 | ND | ND | ND | 22 | |
| Relative *L. oneistus* abundance (%) | | | | | | | | | | | |
| Depth (cmbsf) | A | B | C | D | E | F | G | H | I | Mean | SE |
| 1 to 7 | 0.00 | 7.32 | 4.35 | 8.16 | 1.67 | 0.00 | 11.11 | 0.00 | 0.00 | 3.62 | 1.35 |
| 7 to 13 | 0.00 | 9.76 | 8.70 | 44.90 | 15.00 | 3.57 | 11.11 | 3.33 | 0.00 | 10.71 | 4.34 |
| 13 to 19 | 35.71 | 19.51 | 26.09 | 26.53 | 51.67 | 21.43 | 33.33 | 22.22 | 9.09 | 27.29 | 3.78 |
| 19 to 25 | 64.29 | 63.41 | 8.70 | 20.41 | 15.00 | 75.00 | 44.44 | 74.44 | 90.91 | 50.73 | 9.37 |
| 25 to 31 | ND | ND | 52.17 | ND | 16.67 | 0.00 | ND | ND | ND | 22.95 | 12.56 |
| $\sum H_2S$ (µM) | | | | | | | | | | | |

| Depth (cmbsf) | A | B | C | D | E | F | G | H | I | Mean | SE |
|---|---|---|---|---|---|---|---|---|---|---|---|
| 0 | 0.63 | 0.00 | 0.00 | 0.85 | 0.00 | 0.00 | ND | ND | ND | 0.25 | 0.14 |
| 6 | 0.84 | 0.42 | 0.63 | 1.48 | 1.90 | 0.00 | 0.00 | 0.11 | 0.95 | 0.70 | 0.21 |
| 12 | 1.69 | 0.63 | 2.95 | 1.48 | 6.55 | 3.70 | 6.34 | 0.32 | 1.58 | 2.80 | 0.73 |
| 18 | 2.11 | 1.90 | 2.11 | 1.69 | 13.10 | 0.95 | 6.34 | 2.22 | 4.33 | 3.86 | 1.20 |
| 24 | 5.30 | 1.90 | ND | 1.90 | 22.82 | 1.80 | 14.58 | 15.11 | 6.23 | 8.70 | 2.60 |
| 31 | ND | ND | 4.75 | ND | 16.90 | 6.44 | 31.38 | ND | ND | 14.87 | 5.30 |

Ammonium (µM)

| Depth (cmbsf) | A | B | C | D | E | F | G | H | I | Mean | SE |
|---|---|---|---|---|---|---|---|---|---|---|---|
| 0 | ND | 7.75 | ND | ND | 2.7 | 18.74 | ND | ND | ND | 9.73 | 3.87 |
| 6 | ND | 5.47 | ND | ND | 12.84 | 3.64 | ND | ND | ND | 7.32 | 2.30 |
| 12 | ND | 11.56 | ND | ND | 22.54 | 8.69 | ND | ND | ND | 14.26 | 3.45 |
| 18 | ND | 10.56 | ND | ND | 21.31 | 5.69 | 23.45 | ND | ND | 15.25 | 4.26 |
| 24 | ND | 29.73 | ND | ND | 20.44 | 13.95 | 20.28 | ND | ND | 21.11 | 2.81 |
| 31 | ND | ND | ND | ND | 5.33 | 11.7 | 37.34 | ND | ND | 18.12 | 7.99 |

Nitrate (µM)

| Depth (cmbsf) | A | B | C | D | E | F | G | H | I | Mean | SE |
|---|---|---|---|---|---|---|---|---|---|---|---|
| 0 | ND | ND | ND | ND | 1.41 | 1.13 | 0.79 | ND | ND | 1.11 | 0.25 |
| 6 | ND | 0.00 | ND | ND | 0.29 | ND | 1.08 | ND | ND | 0.46 | 0.46 |
| 12 | ND | ND | ND | ND | 0.15 | ND | 0.33 | ND | ND | 0.24* | 0.06* |
| 18 | ND | ND | ND | ND | 0.00 | ND | 0.42 | ND | ND | 0.21* | 0.14* |
| 24 | ND | ND | ND | ND | 0.38 | 0.48 | 0.34 | ND | ND | 0.38 | 0.48 |
| 31 | ND | ND | ND | ND | ND | ND | 0.47 | ND | ND | 0.47* | NA |

Nitrite (µM)

| Depth (cmbsf) | A | B | C | D | E | F | G | H | I | Mean | SE |
|---|---|---|---|---|---|---|---|---|---|---|---|
| 0 | ND | ND | ND | ND | 0.08 | 0.09 | 0.07 | ND | ND | 0.08 | 0.01 |
| 6 | ND | 0.14 | ND | ND | 0.07 | ND | 0.07 | ND | ND | 0.09 | 0.02 |
| 12 | ND | ND | ND | ND | 0.03 | ND | 0.03 | ND | ND | 0.03* | 0.0* |
| 18 | ND | ND | ND | ND | 0.01 | ND | 0.02 | ND | ND | 0.015* | 0.004* |
| 24 | ND | ND | ND | ND | 0.04 | 0.18 | 0.00 | ND | ND | 0.07 | 0.05 |

| | | | | | | | | | | | |
|---|---|---|---|---|---|---|---|---|---|---|---|
| 31 | ND | ND | ND | ND | ND | ND | 0.07 | ND | ND | 0.07* | NA |
| DOC (mg/L) | | | | | | | | | | | |
| Depth (cmbsf) | A | B | C | D | E | F | G | H | I | Mean | SE |
| 0 | ND | ND | 2.77 | 2.86 | 4.29 | 1.24 | ND | 6.52 | 2.38 | 3.34 | 0.75 |
| 6 | ND | 1.4 | 5.34 | 1.91 | 11.15 | 2.50 | ND | ND | 1.70 | 4.00 | 1.54 |
| 12 | ND | ND | 1.39 | 4.41 | 2.10 | 6.10 | ND | 1.99 | 16.44 | 5.41 | 2.32 |
| 18 | ND | 3.10 | 3.48 | 3.28 | 4.14 | ND | ND | 4.63 | 2.60 | 3.54 | 0.30 |
| 24 | ND | 5.56 | 4.75 | 4.53 | 1.42 | 9.51 | ND | 22.66 | 9.08 | 8.22 | 2.63 |
| 31 | ND | ND | 3.47 | ND | 1.78 | 20.40 | ND | ND | ND | 8.55 | 5.95 |

* Measurements that were taken in fewer than 3 sediment cores due to technical problems and thus means and SE were formed only to visualize the profile trends. Abbreviations: cmbsf, centimeter below seafloor; ND: not determined due to technical difficulties, SE: standard error of the mean, NA: not applicable due to just a single measurement.

**Table S2.** RNA-Seq and EA-IRMS incubation measurements. Note that RNA-Seq and EA-IRMS incubations were performed separately. Samples 1-3 were collected in July 2017, whereas samples 4-6 in March 2019.

RNA-Seq incubations

| Sample | $O_2$ (µM) | | $H_2S$ (µM) | |
|---|---|---|---|---|
| | T-0 | T-24 | T-0 | T-24 |
| anoxic-sulfidic -1 | 0.0 | 0.0 | 11.0 | 7.0 |
| anoxic-sulfidic -2 | 0.0 | 0.0 | 11.0 | 7.0 |
| anoxic-sulfidic -3 | 0.0 | 0.0 | 11.0 | 7.0 |
| anoxic-1 | 0.0 | 0.0 | 0.0 | 0.0 |
| anoxic-2 | 0.0 | 0.0 | 0.0 | 0.0 |
| anoxic-3 | 0.0 | 0.0 | 0.0 | 0.0 |
| anoxic-4 | 0.0 | 0.0 | 0.0 | 0.0 |
| anoxic-5 | 0.0 | 0.0 | 0.0 | 0.0 |
| hypoxic-1 | 111.6 | 31.4 | 0.0 | 0.0 |
| hypoxic-2 | 118.9 | 0.2 | 0.0 | 0.0 |
| hypoxic-3 | 114.8 | 17.4 | 0.0 | 0.0 |
| oxic-1 | 192.0 | 184.8 | 0.0 | 0.0 |
| oxic-2 | 196.0 | 185.0 | 0.0 | 0.0 |
| oxic-3 | 195.0 | 188.3 | 0.0 | 0.0 |
| oxic-4 | 198.3 | 192.4 | 0.0 | 0.0 |
| oxic-5 | 196.7 | 190.5 | 0.0 | 0.0 |
| oxic-6 | 196.9 | 189.7 | 0.0 | 0.0 |

EA-IRMS incubations

| Sample | EA-IRMS Incubation | Weight (mg) | δ13C | $O_2$ | | $H_2S$ | |
|---|---|---|---|---|---|---|---|
| | | | | T0 | T24 | T0 | T24 |
| anoxic-sulfidic (AS) | 13C live nematodes | 1.4 | 314.4 | 0 | 0 | 25 | 1.5 |
| | | 1.1 | 345 | 0 | 0 | 25 | 0.8 |
| | | 0.9 | 355.4 | 0 | 0 | 25 | 1.5 |
| | | 1.2 | 286.7 | 0 | 0 | 25 | 1.2 |
| | 13C dead nematodes | 1.1 | -18.8 | 0 | 0 | 25 | 0.7 |
| | | 0.8 | -22.3 | 0 | 0 | 25 | 0.5 |
| | | 0.9 | -21.5 | 0 | 0 | 25 | 1.5 |
| | 12 C live nematodes | 1.1 | -23.8 | 0 | 0 | 25 | 1.3 |
| | | 1.2 | -24 | 0 | 0 | 25 | 1 |
| | | 1.2 | -24.5 | 0 | 0 | 25 | 0.7 |
| | | 1 | -25.8 | 0 | 0 | 25 | 0.9 |
| hypoxic | 13C live nematodes | 1.3 | 281.7 | 60 | 45 | 0 | 0 |
| | | 1.2 | 414.2 | 60 | 45 | 0 | 0 |
| | | 1 | 458.7 | 59 | 45 | 0 | 0 |
| | 13C dead nematodes | 1.2 | -22.2 | 59 | 42 | 0 | 0 |
| | | 0.3 | -21.9 | 54 | 45 | 0 | 0 |
| | | 1.8 | -21.5 | 61 | 44 | 0 | 0 |
| | 12 C live nematodes | 1 | -25 | 57 | 50 | 0 | 0 |
| | | 1.7 | -24.1 | 55 | 49 | 0 | 0 |
| | | 1.4 | -24.4 | 59 | 52 | 0 | 0 |
| oxic | 13C live nematodes | 1 | 232.9 | 197 | 119 | 0 | 0 |
| | | 1.1 | 277.7 | 196 | 109 | 0 | 0 |
| | | 1.4 | 238.3 | 196 | 111 | 0 | 0 |

T0: before incubation, T24: after 24 h of incubation, δ13C: per mille (‰).

**Table S3.** Functional enrichments of selected gene sets. Statistical enrichment of functional categories was tested for GO terms (GO), Pfam domains (PF), KEGG metabolic maps (map), COG category (COG) and COG general category (uppercase letter) using the Bioconductor software package GOseq (M. D. Young, M. J. Wakefield, G. K. Smyth, and A. Oshlack, Genome Biol 11(2): R14, 2010, https://doi.org/10.1186/gb-2010-11-2-r14).). Functional categories among sets of protein-coding genes were significantly enriched if the adjusted P value (false discovery rate, FDR) was ≤ 0.1. Only FDR values below that threshold are shown; non-significant FDR values are indicated (NS). AS: anoxic-sulfidic; O: hypoxic + oxic.

| | | FDR (threshold 0.1) | | | | | | | | | | |
|---|---|---|---|---|---|---|---|---|---|---|---|---|
| | | anoxic vs AS[1] | | hypoxic vs oxic[1] | | anoxic vs O[1] | | AS vs O[1] | | unchanged over 4 conditions[2] | | |
| Category ID | Description | AS up n = 124 | anoxic up n = 179 | oxic up n = 111 | hypoxic up n = 30 | O up n = 433 | anoxic up n = 366 | O up n = 548 | AS up n = 448 | high n = 163 | low n = 2184 | medium n = 1128 |
| C | Energy production and conversion | NS | NS | NS | NS | NS | **2.9E-05** | NS | **2.4E-04** | NS | NS | NS |
| O | Post-translational modification, protein turnover, and chaperones | NS | NS | NS | NS | **0.009** | NS | **0.022** | NS | **0.085** | NS | NS |
| J | Translation, ribosomal structure and biogenesis | NS | NS | NS | NS | NS | NS | NS | NS | **0.018** | NS | **3.4E-05** |
| P | Inorganic ion transport and metabolism | NS | NS | NS | NS | NS | **0.015** | NS | NS | NS | NS | NS |
| K | Transcription | NS | NS | NS | NS | NS | **0.044** | NS | NS | NS | NS | NS |
| M | Cell wall/membrane/envelope biogenesis | NS | NS | NS | NS | NS | NS | NS | NS | NS | NS | **0.001** |
| T | Signal transduction mechanisms | NS | NS | NS | NS | NS | NS | NS | NS | **0.078** | NS | NS |
| E | Amino acid transport and metabolism | NS | NS | NS | NS | NS | NS | **0.087** | NS | NS | NS | NS |

| | | | | | | | | | | | | |
|---|---|---|---|---|---|---|---|---|---|---|---|---|
| H | Coenzyme transport and metabolism | NS | NS | NS | NS | NS | NS | **0.087** | NS | NS | NS | NS |
| map03010 | Ribosome | NS | NS | NS | NS | NS | NS | NS | NS | **0.019** | NS | **1.3E-05** |
| map00710 | Carbon fixation in photosynthetic organisms | NS | NS | NS | NS | NS | NS | NS | NS | NS | NS | **0.021** |
| map00920 | Sulfur metabolism | NS | NS | NS | NS | NS | **0.017** | NS | **0.066** | NS | NS | NS |
| map00550 | Peptidoglycan biosynthesis | NS | NS | NS | NS | NS | NS | NS | NS | NS | NS | **0.044** |
| map01230 | Biosynthesis of amino acids | NS | NS | NS | NS | NS | NS | NS | NS | NS | NS | **0.096** |
| COG1053 | Succinate dehydrogenase/fumarate reductase, flavoprotein subunit | NS | NS | NS | NS | NS | **0.023** | NS | **0.051** | NS | NS | NS |
| COG2010 | Cytochrome C, mono- and diheme variants | NS | NS | NS | NS | NS | **0.017** | NS | **0.046** | NS | NS | NS |
| COG1708 | Predicted nucleotidyltransferase | NS | NS | NS | NS | NS | **0.045** | NS | NS | NS | NS | NS |
| GO:0051912 | CoB—CoM heterodisulfide reductase activity | **0.041** | NS | NS | NS | NS | **0.021** | NS | **0.046** | NS | NS | NS |
| GO:0020037 | Heme binding | NS | NS | NS | NS | NS | **0.001** | NS | **0.046** | NS | NS | NS |
| GO:0015934 | Large ribosomal subunit | NS | NS | NS | NS | NS | NS | NS | NS | NS | NS | **0.007** |
| GO:0005840 | Ribosome | NS | NS | NS | NS | NS | NS | NS | NS | **0.053** | NS | **0.047** |
| GO:0019843 | rRNA binding | NS | NS | NS | NS | NS | NS | NS | NS | NS | NS | **9.0E-05** |
| GO:0003735 | Structural constituent of ribosome | NS | NS | NS | NS | NS | NS | NS | NS | **0.018** | NS | **9.0E-05** |
| GO:0006412 | Translation | NS | NS | NS | NS | NS | NS | NS | NS | **0.030** | NS | **9.0E-05** |
| GO:0006508 | Proteolysis | NS | NS | NS | NS | NS | NS | NS | NS | NS | NS | **0.015** |
| GO:0009055 | Electron transfer activity | | | | | | **0.081** | | | | | |
| PF13442 | Cytochrome C oxidase, cbb3-type, subunit III | NS | NS | NS | NS | NS | **0.024** | NS | **0.087** | NS | NS | NS |
| PF05168 | HEPN domain | NS | NS | NS | NS | NS | **0.023** | NS | **0.073** | NS | NS | NS |
| PF13609 | Gram-negative porin | NS | **0.100** | NS | NS | NS | NS | NS | NS | NS | NS | NS |

[1] These gene sets only comprise genes that were differentially expressed between indicated conditions (FDR ≤ 0.05, fold-change of 2).

[2] These gene sets comprise genes that were not significantly different between any of the four conditions (oxic, hypoxic, anoxic, and anoxic-sulfidic), and were furthermore classified based on expression level (low, medium, high) over all four conditions based on hierarchical clustering with Euclidean distances.

**Table S4.** Top expressed *Ca*. T. oneisti proteins. Symbiont proteins under each condition were ranked by relative abundance (%cOrgNSAF), and the 30 proteins with functional annotation exhibiting the highest relative abundance per condition are shown. Proteins written in bold were only found in the top expressed proteins under the respective condition. Color gradients indicate relative abundance and are scaled between values 5.76 and 0.39. Relative abundance of symbiont proteins under two different conditions was determined as described in Supplementary Materials & Methods and is displayed as mean values of three individual replicates per condition. Data S1 specifies all proteins that were detected to be expressed (column "Proteome detection").

| Locus tag | Description | Protein name | Mean % cOrgNSAF |
|---|---|---|---|
| Anoxic | | | |
| TONNANOP_v1_90031 | porin | NA | 5.76 |
| TONNANOP_v1_90030 | porin | NA | 4.09 |
| TONNANOP_v1_450027 | chaperone Hsp60 | GroEL | 3.98 |
| TONNANOP_v1_90020 | OmpA/MotB domain protein | NA | 2.93 |
| TONNANOP_v1_880016 | (R)-3-hydroxybutyryl-CoA dehydrogenase | PhaB | 2.07 |
| TONNANOP_v1_480022 | heat shock chaperone | IbpA | 1.39 |
| TONNANOP_v1_680020 | phasin family protein | NA | 1.26 |
| TONNANOP_v1_400015 | phasin family protein | NA | 1.15 |
| TONNANOP_v1_1690006 | adenylyl-sulfate reductase subunit alpha | AprA-1 | 1.12 |
| TONNANOP_v1_130028 | fructose-1,6-bisphosphate aldolase, class II | Fda | 1.01 |
| TONNANOP_v1_580017 | ribulose-1,5-bisphosphate carboxylase oxygenase large subunit | CbbL | 0.94 |
| TONNANOP_v1_390010 | short-chain dehydrogenase | NA | 0.79 |
| TONNANOP_v1_130050 | glyceraldehyde-3-phosphate dehydrogenase A | GapA | 0.74 |
| TONNANOP_v1_50022 | phosphoenolpyruvate carboxykinase (GTP) | PckG | 0.68 |
| TONNANOP_v1_670008 | **rubrerythrin** | **NA** | **0.67** |

| | | | |
|---|---|---|---|
| TONNANOP_v1_2780001 | elongation factor Tu | TufB | 0.63 |
| TONNANOP_v1_40089 | succinyl-CoA synthetase, beta subunit | SucC | 0.62 |
| TONNANOP_v1_380023 | respiratory polysulfide/ thiosulfate reductase, chain A | PsrA/PhsA | 0.61 |
| TONNANOP_v1_930011 | protease activity modulator | HflK | 0.58 |
| TONNANOP_v1_850028 | F0F1 ATP synthase subunit beta | AtpD | 0.53 |
| TONNANOP_v1_880015 | PHA synthase subunit E | PhaE | 0.51 |
| TONNANOP_v1_520016 | glutamate--ammonia ligase | GlnA | 0.50 |
| TONNANOP_v1_270021 | phosphate dikinase | PpdK | 0.46 |
| TONNANOP_v1_850016 | **F0F1 ATP synthase subunit B** | **AtpF** | **0.45** |
| TONNANOP_v1_1190006 | lipase | NA | 0.45 |
| TONNANOP_v1_1530001 | **malate dehydrogenase** | **Mdh** | **0.45** |
| TONNANOP_v1_1770003 | **adenylyl-sulfate reductase subunit alpha** | **AprA-2** | **0.42** |
| TONNANOP_v1_1690009 | **adenylylsulfate reductase membrane anchor** | **AprM** | **0.42** |
| TONNANOP_v1_1490013 | **50S ribosomal L14** | **RplN** | **0.41** |
| TONNANOP_v1_600015 | **malyl-CoA/beta-methylmalyl-CoA/citramalyl-CoA lyase** | **Mcl** | **0.40** |
| Oxic | | | |
| TONNANOP_v1_90031 | porin | NA | 4.80 |
| TONNANOP_v1_450027 | chaperone Hsp60 | GroEL | 4.03 |
| TONNANOP_v1_480022 | heat shock chaperone | IbpA | 3.83 |
| TONNANOP_v1_90030 | porin | NA | 3.42 |
| TONNANOP_v1_90020 | OmpA/MotB domain protein | NA | 1.89 |
| TONNANOP_v1_1690006 | adenylyl-sulfate reductase subunit alpha | AprA-1 | 1.60 |
| TONNANOP_v1_880016 | (*R*)-3-hydroxybutyryl-CoA dehydrogenase | PhaB | 1.56 |
| TONNANOP_v1_580017 | ribulose-1,5-bisphosphate carboxylase oxygenase large subunit | CbbL | 1.04 |
| TONNANOP_v1_680020 | phasin family protein | NA | 0.86 |
| TONNANOP_v1_130028 | fructose-1,6-bisphosphate aldolase, class II | Fda | 0.85 |
| TONNANOP_v1_400015 | phasin family protein | NA | 0.81 |
| TONNANOP_v1_50022 | phosphoenolpyruvate carboxykinase (GTP) | PckG | 0.70 |
| TONNANOP_v1_850028 | F0F1 ATP synthase subunit beta | AtpD | 0.70 |
| TONNANOP_v1_930011 | protease activity modulator | HflK | 0.65 |
| TONNANOP_v1_390010 | short-chain dehydrogenase | NA | 0.58 |

| | | | |
|---|---|---|---|
| TONNANOP_v1_380023 | respiratory polysulfide/ thiosulfate reductase, chain A | PsrA/PhsA | 0.58 |
| TONNANOP_v1_130050 | glyceraldehyde-3-phosphate dehydrogenase A | GapA | 0.58 |
| TONNANOP_v1_270021 | phosphate dikinase | PpdK | 0.56 |
| TONNANOP_v1_880015 | PHA synthase subunit E | PhaE | 0.54 |
| TONNANOP_v1_1530001 | lipase | NA | 0.49 |
| TONNANOP_v1_2780001 | elongation factor Tu | TufB | 0.46 |
| **TONNANOP_v1_850018** | **F0F1 ATP synthase subunit alpha** | **AtpA** | **0.45** |
| **TONNANOP_v1_20064** | **chaperone Hsp70** | **DnaK** | **0.45** |
| TONNANOP_v1_520016 | glutamate--ammonia ligase | GlnA | 0.44 |
| TONNANOP_v1_40089 | succinyl-CoA synthetase, beta subunit | SucC | 0.44 |
| **TONNANOP_v1_190055** | **cytochrome c oxidase subunit II** | **CtaC** | **0.44** |
| TONNANOP_v1_930010 | protease modulator | HflC | 0.44 |
| **TONNANOP_v1_840002** | **molecular chaperone Hsp90** | **HtpG** | **0.43** |
| **TONNANOP_v1_890008** | **sodium-translocating pyrophosphatase** | **HppA** | **0.42** |
| **TONNANOP_v1_100014** | **1,4-alpha-glucan branching enzyme** | **GlgB** | **0.39** |

NA: not applicable

**Table S5.** List of all ribulose-1,5-bisphosphate carboxylase (RuBisCO) protein sequences used for constructing the phylogenetic tree in Figure S8.

<table>
<tr><th>Organism</th><th>RuBisCO form[1]</th><th>Amino acid sequence</th></tr>
<tr><td>Thioflavicoccus mobilis</td><td>IA</td><td>>tr|L0GWD2|L0GWD2_9GAMM Ribulose bisphosphate carboxylase large chain OS=Thioflavicoccus mobilis 8321 OX=765912 GN=cbbL PE=3 SV=1<br>MAKTYNAGVKDYRETYWMPDYTPKDTDILACFKITPQPGVPREEAAAAVAAESSTGTWTT<br>VWTDLLTDLDYYKGRAYAIEDVPGDDERFYAFIAYPIDLFEEGSVVNVFTSLVGNVFGFK<br>AVRTLRLEDVRFPIAYVMTCNGPPHGIQVERDIMNKYGRPLLGCTIKPKLGLSAKNYGRA<br>VYECLRGGLDFTKDDENVNSQPFMRWRQRFDFVMEAIDKAESETGERKGHYLNVTAPTPD<br>EMFKRAEHAKELGAPIIMHDYITGGWCANTGLAQWCRDNGVLLHIHRAMHAVIDRHPHHG<br>IHFRVLAKLLRLSGGDHLHTGTVVGKLEGDRAATLGWIDLLRESYVEEDRSRGIFFDQDW<br>GSMPGVFAVASGGIHVWHMPALVTIFGDDAVLQFGGGTLGHPWGNAAGAAANRVALEACV<br>EARNRGVAIEKEGKEVLTAAANNSPELKAAMETWKEIKFEFDTVDKLDVAHK</td></tr>
<tr><td>Lamprocystis purpurea</td><td>IA</td><td>>WP_020507467.1 ribulose-bisphosphate carboxylase large subunit [Lamprocystis purpurea]<br>MAKSYSAGVKEYRETYWMPDYTPKDTDILACFKITPQPGVPREEAAAAVAAESSTGTWTTVWTDLLTDLDYYK<br>GRAYAIEDVPGDDSCYYAFIAYPIDLFEEGSVVNVFTSLVGNVFGFKAVRALRLEDVRFPIAYVMTCGGPPHGIQ<br>VERDMLNKYGRPLLGCTIKPKLGLSAKNYGRAVYECLRGGLDLTKDDENVNSQPFMRWRQRFDFVMDAIDKA<br>ERETGERKGHYLNVTAPTPEEMYKRAEYAKEIGAPIIMHDYITGGFCANTGLANWCRDNGILLHIHRAMHAVLD<br>RNPHHGIHFRVLAKILRLSGGDHLHSGTVVGKLEGDREATLGWIDIMRDKFIKEDRSRGIFFDQDWGSMPGVFP<br>VASGGIHVWHMPALVTIFGDDACLQFGGGTLGHPWGNAAGAAANRVALEACVQARNEGIAIEKEGKDVLTKAA<br>KHSPELKVAMETWKEIKFEFDTVDKLDVAHK</td></tr>
<tr><td>Allochromatium vinosum</td><td>IA</td><td>>sp|P22859|RBL1B_ALLVD Ribulose bisphosphate carboxylase large chain 2 OS=Allochromatium vinosum (strain ATCC 17899 / DSM 180 / NBRC 103801 / NCIMB 10441 / D) OX=572477 GN=cbbL2 PE=1 SV=4<br>MSTKTYDAGVKDYALTYWTPDYVPLDSDLLACFKVTPQAKVSREEAAAAVAAESSTGTWT<br>TVWSDLLTDLDYYKGRAYRIEDVPGDKESFYAFIAYPLDLFEEGSIVNVLTSLVGNVFGF<br>KAVRALRLEDIRFPLHYVKTCGGPPNGIQVERDRMDKYGRPFLGATVKPKLGLSAKNYGR<br>AVYEMLRGGLDFTKDDENVNSQPFMRWQNRFEFVSEAVRKAQEETGERKGHYLNVTAPTC<br>EEMFKRAEFAKECGAPIIMHDFLTGGFTANTSLANWCRDNGMLLHIHRAMHAVIDRNPKH<br>GIHFRVLAKCLRLSGGDHLHTGTVVGKLEGDRQSTLGFVDQLRESFIPEDRSRGLFFDQD<br>WGGMPGVMAVASGGIHVWHIPALVTIFGDDSVLQFGGGTQGHPWGNAAGAAANRVATEAC<br>VKARNEGVEIEKHAREVLSDAARHSPELAVAMETWKEIKFEFDVVDKLDAA</td></tr>
</table>

*Marichromatium purpuratum* IA

>tr|W0E2V6|W0E2V6_MARPU Ribulose bisphosphate carboxylase large chain OS=Marichromatium purpuratum 984 OX=765910 GN=rbcL PE=3 SV=1
MSTKTYDAGVKDYALTYWTPDYVPLDSDLLACFKVTPQANVPREEAAAAVAAESSTGTWT
TVWSDLLTDLDYYKGRAYRIEDVPGDKESFYAFVAYPIDLFEEGSIVNVLTSLVGNVFGF
KAVRHLRLEDIRFPLHYVKTCGGPPNGIQVERDRMDKYGRPFLGATVKPKLGLSAKNYGR
AVYEMLRGGLDFTKDDENVNSQPFMRWQNRFEFVMEAVRKAQEETGERKGHYLNVTAPTC
EEMFKRAEFAKELGAPIIMHDFLTGGFTANTSLANWCRDNGMLLHIHRAMHAVIDRHPKH
GIHFRVLAKCLRLSGGDHLHTGTVVGKLEGDRRSTLGFVDAIRESFVPEDRSRGLFFDQD
WGGMPGLLAVASGGIHVWHIPALVAIFGDDAVLQFGGGTQGHPWGNAAGAAANRVATEAC
IKARNEGVEVEQHAREILTEAARHSPELAAAMETWKEIKFEFDTVDKLDAA

*Solemya velum symbiont* IA

>sp|Q673V5|RBL_SOVGS Ribulose bisphosphate carboxylase large chain OS=Solemya velum gill symbiont OX=2340 GN=cbbL PE=3 SV=1
MAKTYDAGVKEYRETYWMPEYTPLDTDILACFKVTPQPGVPREEVAAAVAAESSTGTWTT
VWTDLLTDLDHYKGRAYAIEDVPGDDTCFYAFIAYPIDLFEEGSVVNVMTSLVGNVFGFK
ALRALRLEDIRFPIAYVMTCNGPPQGIQVERDLLNKYGRPLLGCTIKPKLGLSAKNYGRA
CYEGLRGGLDFTKDDENVNSQPFMRWRHRFDFVMEAIQKAEAETGERKGHYLNVTAPTSD
EMMKRAEYAKEIGAPIIMHDYITGGWSANTQLAQWCQDNGMLLHIHRAMHAVLDRNPHHG
IHFRVLTKILRLSGGDHLHSGTVVGKLEGDREATLGWIDIMRDSFNKEDRSRGIFFDQDW
GSMPGVLPVASGGIHVWHMPALVNIFGDDSVLQFGGGTLGHPWGNAAGAAANRVAVEACV
EARNNGRELEKEGKEILTTAAAHSPELKAAMETWKEIKFEFDTVDKLDVSHK

*Hydrogenophaga pseudoflava* IA

>sp|Q9ZB35|RBL_HYDPS Ribulose bisphosphate carboxylase large chain OS=Hydrogenophaga pseudoflava OX=47421 GN=cbbL PE=1 SV=3
MATKTYNAGVKEYRSTYWEPHYTPKDTDILACFKITPQPGVDREEVAAAVAAESSTGTWT
TVWTDLLTDLDYYKGRAYRIEDVPGDDTCFYAFVAYPIDLFEEGSVVNVLTSLVGNVFGF
KALRALRSEDVRFPIAYVKTCGGPPHGIQVERDIMNKYGRPLLGCTIKPKLGLSGKNYGR
AVYECLRGGLDFTKDDENVNSQPFMRWPQRFDFEQEAIEKAHGETGERKVHYLNVTAPTP
GEMYKRAEYAKELGAPIIMHDYLTGGLCANTGLANWCRDNGMLLHIHRAMHAELDRNPHH
GIHFRVLTKVLRLSGRDHLHSGTVVGKLEGDRASTLGWIDIMRDTFIKEDRSRGIFFDQD
FGSMPGVMPVASGGIHVWHMPALVNIFGDDSVLQFGGGTVGHPWGNAPGATANRVELEAC
VKARNEGIAVEKEGKAVLTEAANDSPELKIAMETWKEIKFEFDTVDKLDIAHK

*Nitrosomonas europaea* IA

>sp|Q82TG6|RBL_NITEU Ribulose bisphosphate carboxylase large chain OS=Nitrosomonas europaea (strain ATCC 19718 / CIP 103999 / KCTC 2705 / NBRC 14298) OX=228410 GN=cbbL PE=3 SV=1
MSAKTYNAGVKEYRHTYWEPHYNVQDTDILACFKIVPQPGVDREEAAAAVAAESSTGTWT
TVWTDLLTDLDYYKGRSYRIEDVPGDDSSFYAFIAYPIDLFEEGSVVNVLTSLTGNVFGF
KAVRSLRLEDVRFPIAYVKTCGGPPNGIQVERDILNKYGRAYLGCTIKPKLGLSAKNYGR

| | | |
|---|---|---|
| | | AVYECLRGGLDFTKDDENVNSQPFMRWRQRFDFVMEAIHKAERETGERKGHYLNVTAPTP<br>EEMFKRAEYAKELKAPIIMHDYIAGGFCANTGLANWCRDNGILLHIHRAMHAVIDRNPHH<br>GIHFRVLAKMLRLSGGDHLHSGTVVGKLEGDREATLGWIDIMRDSFIKEDRSRGIMFDQD<br>WGSMPGVVPVASGGIHVWHMPALVTIFGDDACLQFGGGTLGHPWGNAAGAAANRVALEAC<br>VEARNRGVPIEKEGKAILTEAAKHSPELKIAMETWKEIKFEFDTVDKLDVAHK |
| *Halorhodospira halophila* | IA | >sp\|A1WVW0\|RBL_HALHL Ribulose bisphosphate carboxylase large chain OS=Halorhodospira halophila (strain DSM 244 / SL1) OX=349124 GN=cbbL PE=3 SV=1<br>MASKTYTAGVKDYRETYWEPDYKIKDSDLLAVFKVTPQPGVDREEAAAAVAAESSTGTWT<br>TVWTDLLTDLEHYKGRAYKVEDVPGDDEAFYAFIAYPIDLFEEGSIVNVFTSLVGNVFGF<br>KAVRALRLEDVRFPLHFVMTCPGPPNGIQVERDKMNKYGRPLLGCTIKPKLGLSAKNYGR<br>AVYECLRGGLDFTKDDENVNSQPFMRWRDRFEFVMEAIQKAEEETGERKGHYLNVTAPTP<br>EEMYKRAEFAKELGAPIIMHDYITAGFCAHQGLANWCRDNGMLLHIHRAMHAVLDRNPNH<br>GIHFRVLTKILRLMGGDQLHTGTVVGKLEGDRQSTLGWIDLLRKPYIEEDRSRGLFFDQD<br>WGAMPGAFAVASGGIHVWHMPALLSIFGDDAVFQFGGGTLGHPWGNAAGAAANRVALEAC<br>VKARNEGRELEKEGKEILTEAAKSSPELKAAMETWKEIKFEFDTVDKLDTAHR |
| *Prochlorococcus marinus* | IA | >sp\|Q7VD33\|RBL_PROMA Ribulose bisphosphate carboxylase large chain OS=Prochlorococcus marinus (strain SARG / CCMP1375 / SS120) OX=167539 GN=cbbL PE=3 SV=1<br>MSKKYDAGVKEYRDTYWTPDYVPLDTDLLACFKCTGQEGVPREEVAAAVAAESSTGTWST<br>VWSELLTDLEFYKGRCYRIEDVPGDKESFYAFIAYPLDLFEEGSITNVLTSLVGNVFGFK<br>ALRHLRLEDIRFPMAFIKTCGGPPQGIVVERDRLNKYGRPLLGCTIKPKLGLSGKNYGRV<br>VYECLRGGLDLTKDDENINSQPFQRWRDRFEFVAEAVKLAQQETGEVKGHYLNCTATTPE<br>EMYERAEFAKELDMPIIMHDYITGGFTANTGLANWCRKNGMLLHIHRAMHAVIDRHPKHG<br>IHFRVLAKCLRLSGGDQLHTGTVVGKLEGDRQTTLGYIDNLRESFVPEDRTRGNFFDQDW<br>GSMPGVFAVASGGIHVWHMPALLAIFGDDSCLQFGGGTHGHPWGSAAGAAANRVALEACV<br>KARNAGREIEKESRDILMEAAKHSPELAIALETWKEIKFEFDTVDKLDVQ |
| *Thioalkalivibrio nitratireducens* | IA | >tr\|L0DYE9\|L0DYE9_THIND Ribulose bisphosphate carboxylase large chain OS=Thioalkalivibrio nitratireducens (strain DSM 14787 / UNIQEM 213 / ALEN2) OX=1255043 GN=cbbL [H] PE=3 SV=1<br>MAVKTYEAGVKEYREKYWTPDYVPLDTDLLACFKVTGQPGVPREEVAAAVAAESSTGTWS<br>TVWSELLTDLEYYKGRAYRIEDVPGDKESFYAFVAYPLDLFEEGSIVNVLTSLVGNVFGF<br>KALKHLRLEDIRFPIAYIKTCMGPPSGIQVERDKLNKYGRPMLGATIKPKLGLSAKNYGR<br>AVYECLRGGLDLTKDDENVNSQPFMRWQNRFEFVAEAVMKAQAETGERKGHYLNVTAPDP<br>EQMYERAEFAKELGMPIVMHDFLTGGFTANTGLAKWCRKNGILLHIHRAMHAVIDRHPKH<br>GIHFRVLAKCLRLSGGDHLHTGTVVGKLEGDRNSTLGFVDQLREAFVPEDRARGVFFDQD<br>WGSMPGVFAVASGGIHVWHMPALVAIFGDDSVLQFGGGTQGHPWGNAAGAAANRVALEAC |

<table>
<tr><td></td><td></td><td>VKARNEGRELEREAREILTDAARHSPELAIAMETWKEIKFEFETVDKLDVG</td></tr>
<tr><td>Ectothiorhodospira haloalkaliphila</td><td>IA</td><td>>tr|W8KKV1|W8KKV1_9GAMM Ribulose bisphosphate carboxylase large chain OS=Ectothiorhodospira haloalkaliphila OX=421628 GN=rbcL PE=3 SV=1<br>MAVKTYSAGVKDYRQTYWTPEYTPRDTDILAVFKITPQPGVDREEAAAAVAAESSTGTWT<br>TVWTDLLTDMDYYKGRAYQIEDVPGDDECFYAFIAYPIDLFEEGSVVNVFTSLVGNVFGF<br>KAVRTLRLEDVRFPIAYVKTCGGPPHGIQVERDIMNKYGRGLLGCTIKPKLGLSAKNYGR<br>AVYECLRGGLDFTKDDENVNSQPFMRWRQRFDFVMEAIDKAEQETGERKGHYLNVTAPTP<br>EEMYKRAEYAKEIGAPIIMHDYITGGFCANTGLANWCRENGMLLHIHRAMHAVMDRHPRH<br>GIHFRVLAKILRLSGGDHLHTGTVVGKLEGDRDATLGWIDLLREDYVKEDRSRGIFFDQD<br>WGSMPGVFAVASGGIHVWHMPALVSIFGDDAVFQFGGGTLGHPWGNAAGATANRVALEAC<br>VQARNEGRELEKEGKDILTAAAGHSPELKVAMETWKEIKFEFDTVDKLDVQHR</td></tr>
<tr><td>Ca. Thiodiazotropha endoloripes A</td><td>IA</td><td>>tr|A0A1E2V426|A0A1E2V426_9GAMM Ribulose bisphosphate carboxylase large chain OS=Candidatus Thiodiazotropha endoloripes OX=1818881 GN=cbbL PE=3 SV=1<br>MAKKYDAGVKEYRETYWMPEYTPLDTDILACFKVTPQPGVPREEVAAAVAAESSTGTWTT<br>VWTDLLTDLDHYKGRAYAIEDVPGDDTCFYAFVAYPIDLFEEGSVVNVMTSLVGNVFGFK<br>ALRALRLEDIRFPIAYVMTCNGPPQGIQVERDMLNKYGRPLLGCTIKPKLGLSAKNYGRA<br>CYEGLRGGLDFTKDDENVNSQPFMRWKHRFDFVMEAIQKAEAETGERKGHYLNVTAPTSD<br>EMMKRAEYAKEIGAPIIMHDYITGGWSANTQLAQWCQDNGMLLHIHRAMHAVLDRNPHHG<br>IHFRVLTKILRLSGGDHLHSGTVVGKLEGDRDATLGWIDIMRDSFVKEDRSRGIFFDQDW<br>GSMPGVLPVASGGIHVWHMPALVNIFGDDSVLQFGGGTLGHPWGNAAGAAANRVAVEACV<br>EARNQGRELEKEGKDILTNAAASSPELKAAMETWKEIKFEFDTVDKLDVSHK</td></tr>
<tr><td>Ca. Thiosymbion oneisti</td><td>IA</td><td>>TONNANOP_v1_580017|ID:49048881|cbbL| Ribulose bisphosphate carboxylase large chain 1 [Candidatus Thiosymbion oneisti nanopore BZ 1385A run1run2]<br>MAKTYSAGVKEYRETYWMPDYTPKDTDILACFKITPQPGVPREEAAAAVAAESSTGTWTT<br>VWTDLLTDLDHYKGRAYAIEDVPGDDECYYAFIAYPIDLFEEGSVVNVFTSLVGNVFGFK<br>AVRTLRLEDVRFPIAYVMTCNGPPHGIQVERDKFNKYGRALLGCTIKPKLGLSAKNYGRA<br>VYECLRGGLDLTKDDENVNSQPFMRWRNRFEFVMEAIHKAEKETGERKGHYLNVTAPTPE<br>EMYKRAEFAKELGAPIIMHDFLTGGFTANTGLAQWCRDNGILLHIHRAMHAVLDRHPRHG<br>IHFRVLAKALRLSGGDHLHSGTVVGKLEGDRDATLGWIDIMRDSFIKEDRSRGIFFDQDW<br>GSMPGVFPVASGGIHVWHMPALVTIFGDDACLQFGGGTLGHPWGNAAGAAANRVALEACV<br>EARNQGIPVEKEGKEVLTKAAASSPELKAAMETWKEIKFEFDTVDKLDVAHK</td></tr>
<tr><td>Olavius algarvensis γ1 symbiont</td><td>IA</td><td>>tr|B5QSJ5|B5QSJ5_9GAMM Ribulose-1,5-bisphosphate large subunit (Fragment) OS=Olavius algarvensis Gamma 1 endosymbiont OX=260705 GN=cbbL PE=3 SV=1<br>TWTTVWTDLLTDLDHYKGRAYAIEDVPGDDECYYAFVAYPIDLFEEGSVVNVFTSLVGNV<br>FGFKAIRTLRLEDVRFPIAYVMTCNGPPNGIQVERDKFNKYGRALLGCTIKPKLGLSAKN</td></tr>
</table>

| | | |
|---|---|---|
| | | YGRAVYECLRGGLDLTKDDENINSQPFMRWRNRFEFVMEAIEKAEKETGERKGHYLNVTA<br>PTPEEMYKRAEFAKELGAPIIMHDFLTGGFTANTGLAQWCRDNGMLLHMHR |
| *Thioalkalimicrobium cyclicum* | IA | >tr\|F6DAZ0\|F6DAZ0_THICA Ribulose bisphosphate carboxylase large chain OS=Thiomicrospira cyclica (strain DSM 14477 / JCM 11371 / ALM1) OX=717773 GN=cbbL PE=3 SV=1<br>MANQTFNAGVQDYKLTYWTPDYTPLDTDLLACFKVIPQAGVPREEAAAAVAAESSTGTWT<br>TVWTDLLTDMEFYKGRCYRIEDVPGNKDAFYAFIAYPLDLFEEGSVVNVLTSLVGNVFGF<br>KAVRSLRLEDIRFPVAFIKTCGGPPSGIQVERDKLNKYGRPMLGCTIKPKLGLSAKNYGR<br>AVYECLRGGLDLTKDDENINSQPFQRWRDRFSFVADAINKAEAETGEVKGHYLNVTAATC<br>EDMMERAEYAKELGVRIVMHDFLTGGFTANTSLANWCRKNGMLLHIHRAMHAVIDRNPNH<br>GIHFRVLAKCLRLSGGDHLHTGTVVGKLEGDRASTLGFVDQLREAFVPEDRSRGVFFDQD<br>WGSMPGVMAVASGGIHVWHMPALVTIFGDDSVLQFGGGTQGHPGGNAAGAAANRVALEAC<br>VKARNEGRDLEREGGDILRDAARHSPELAVALETWKEIKFEFDTVDKLDA |
| *Ca.* Thiobios zoothamnicoli | IA | >tr\|B8QCU8\|B8QCU8_9GAMM Ribulose-1,5-bisphosphate carboxylase/oxygenase large subunit form I (Fragment) OS=Candidatus Thiobios zoothamnicoli OX=374667 PE=4 SV=1<br>DFTKDDENVNSQPFMRWRDRFEFVGEAIQKAEQETGEKKGHYLNVTAATPEEMYKRAEFA<br>KEVGSPILMHDFITGGFTANTGLANWCRDNGMLLHIHRAMHAVIDRHPKHGIHFRVLAKC<br>LRLSGGDHLHTGTVVGKLEGDRQSTLGFVDQLRESFVPEDRSRGVFFDQDWGSMPGVFAV<br>ASGGIHVWHMPALVTIFGDDSMLQFGGGTQGHPWGNAAGAAANRVALEASVKARNEGREI<br>EKEARDILTEAAKHSPELAIAMETWKEIKF |
| *Bathymodiolus thermophilus* | IA | >tr\|A0A1J5TXR2\|A0A1J5TXR2_9GAMM Ribulose bisphosphate carboxylase large chain OS=Bathymodiolus thermophilus thioautotrophic gill symbiont OX=2360 GN=cbbL PE=3 SV=1<br>MAKVYDAGVKDYRETYWMPDYTPKETDILACFKVTPQDGVPREEVAAAVAAESSTGTWTT<br>VWTDLLTDLDYYKGRCYAIEDVPGDDTCFYAFIAYPIDLFEEGSVVNIMTSLVGNVFGFK<br>ALRALRLEDIRFPIAYVMTCNGPPQGIQLERDILNKYGRPLLGCTIKPKLGLSAKNYGRA<br>CYEGLRGGLDFTKDDENVNSQPFMRWRARFDFVQEAIEKAEAETGERKGHYLNVTAPTSD<br>EMMKRAEYAKEIGSPIIMHDYITGGWSANTQLAQWCQDNGMLLHIHRAMHAVLDRNPHHG<br>IHFRVLTKILRLSGGDHLHSGTVVGKLEGDRDATLGWIDIMRDSYIKEDRSRGIFFDQDW<br>GAMPGVIPVASGGIHVWHMPALVNIFGDDSCLQFGGGTLGHPWGNAAGAAANRVAVEACV<br>EARNTGRELEKEGKDILTTAAKHSPELAIAMETWKEIKFEFDTVDKIDVAHK |
| *Thiomonas intermedia* | IA | >sp\|Q9ZHZ1\|RBL1_THIK1 Ribulose bisphosphate carboxylase large chain OS=Thiomonas intermedia (strain K12) OX=75379 GN=cbbL PE=3 SV=1<br>MAVKTYQAGVKEYRQTYWMPEYTPLDTDLLACFKITPQAGVDREEAAAAVAAESSTGTWT<br>TVWTDLLTDMDYYKGRAYRIEDVPGDDTCFYAFIAYPIDLFEEGSVVNVFTSLVGNVFGF<br>KAIRALRLEDIRFPIAYVKTCNGPPNGIQVERDVINKYGRPLLGCTIKPKLGLSGKNYGR<br>AVYECLRGGLDFTKDDENINSQPFMRWKQRFDFVQEATLKAEQETGERKGHYLNVTAPTP |

| | | |
|---|---|---|
| | | DEMFKRAEYAKEIGAPIIMHDYITGGFCANTGLAQWCRDNGMLLHIHRAMHAVLDRNPHH<br>GIHFRVLTKILRLSGGDHLHTGTVVGKLEGDRASTLGWIDLLRESYVPEDRSRGIFFDQD<br>WGSMPGAFAVASGGIHVWHMPALVTIFGDDSVLQFGGGTLGHPWGNAAGAAANRVALEAC<br>VQARNEGRQVEKEGREILTAAAQHSPELKIAMETWKEIKFEFDTVDKLDVTNK |
| *Halothiobacillus neapolitanus* | IA | >sp\|O85040\|RBL1_HALNC Ribulose bisphosphate carboxylase large chain OS=Halothiobacillus neapolitanus (strain ATCC 23641 / c2) OX=555778 GN=cbbL PE=1 SV=1<br>MAVKKYSAGVKEYRQTYWMPEYTPLDSDILACFKITPQPGVDREEAAAAVAAESSTGTWT<br>TVWTDLLTDMDYYKGRAYRIEDVPGDDAAFYAFIAYPIDLFEEGSVVNVFTSLVGNVFGF<br>KAVRGLRLEDVRFPLAYVKTCGGPPHGIQVERDKMNKYGRPLLGCTIKPKLGLSAKNYGR<br>AVYECLRGGLDFTKDDENINSQPFMRWRDRFLFVQDATETAEAQTGERKGHYLNVTAPTP<br>EEMYKRAEFAKEIGAPIIMHDYITGGFTANTGLAKWCQDNGVLLHIHRAMHAVIDRNPNH<br>GIHFRVLTKILRLSGGDHLHTGTVVGKLEGDRASTLGWIDLLRESFIPEDRSRGIFFDQD<br>WGSMPGVFAVASGGIHVWHMPALVNIFGDDSVLQFGGGTLGHPWGNAAGAAANRVALEAC<br>VEARNQGRDIEKEGKEILTAAAQHSPELKIAMETWKEIKFEFDTVDKLDTQNR |
| *Hydrogenovibrio marinus* | IA | >sp\|Q59458\|RBL1A_HYDMR Ribulose bisphosphate carboxylase large chain 1 OS=Hydrogenovibrio marinus OX=28885 GN=cbbL1 PE=1 SV=1<br>MAKTYNAGVKEYRETYWMPEYEPKDSDFLACFKVVPQPGVPREEIAAAVAAESSTGTWTT<br>VWTDLLTDLDYYKGRAYRIEDVPGDDSAFYAFIAYPIDLFEEGSIVSVMTSLVGNVFGFK<br>ALRSIRLEDIRFPLAYVMTCGGPPHGIQVERDKMDKYGRPMLGCTIKPKLGLSAKNYGRA<br>VYECLRGGLDFTKDDENVTSQPFMRWRDRFLFCQDAIEKAQDETGERTGHYLNATAGTPE<br>EMYERAEFAKEIGSPIVMHDFLTGGLTANTGLANYCRKNGLLLHIHRAMHGVIDRNPLHG<br>IHFRVLSKVLRLSGGDHLHSGTVVGKLEGDRGSDLGWIDIMRDSFIAEDRSRGIMFDQDF<br>GEMPGVIPVASGGIHVWHMPALVAIFGDDSVLQFGGGTIGHPWGNAVGAAVNLVALEACV<br>QARNEGQEIEKNGKEILTNDGKHSPELKIAMETWKEIKFEFDTVDKLDLSHK |
| *Synechococcus* | IB | >sp\|Q02518\|RBL_SYNSP Ribulose bisphosphate carboxylase large chain OS=Synechococcus sp. OX=1131 GN=cbbL PE=3 SV=1<br>MAYTQSKSQKVGYQAGVKDYRLTYYTPDYTPKDTDILAAFRVTPQPGVPFEEAAAAVAAE<br>SSTGTWTTVWTDLLTDLDRYKGRCYDIEPLPGEDNQFIAYIAYPLDLFEEGSVTNMLTSI<br>VGNVFGFKALKALRLEDLRIPVAYLKTFQGPPHGIQVERDKLNKYGRPLLGCTIKPKLGL<br>SAKNYGRAVYECLRGGLDFTKDDENINSQPFQRWRDRFLFVADAIHKAQAETGEIKGHYL<br>NVTAPTCEEMLKRAEFAKDWNAIIMHDFLTAGFTANTTLSKGCRDNGMLLHIHRAMHAVM<br>DRQKNHGIHFRVLAKCLRMSGGDHIHTGTVVGKLEGDKAVTLGFVDLLRENYIEQDRSRG<br>IYFTQDWASMPGVMAVASGGIHVWHMPALVDIFGDDAVLQFGGGTLGHPWGNAPGATANR<br>VALEACIQARNEGRDLMREGGDIIREAARWSPELAAACELWKEIKFEFEAQDTI |

| | | |
|---|---|---|
| *Thiobacillus denitrificans* | IA | >sp\|Q56259\|RBL1_THIDA Ribulose bisphosphate carboxylase large chain OS=Thiobacillus denitrificans (strain ATCC 25259) OX=292415 GN=cbbL PE=1 SV=2<br>MAVKTYSAGVKEYRQTYWMPEYTPLDTDILACFKITPQAGVDREEAAAAVAAESSTGTWT<br>TVWTDLLTDLDYYKGRAYAIEDVPGDDTCFYAFIAYPIDLFEEGSVVNVFTSLVGNVFGF<br>KAVRALRLEDVRFPIAYVKTCGGPPHGIQVERDVMNKYGRPLLGCTIKPKLGLSAKNYGR<br>AVYECLRGGLDFTKDDENVNSQPFMRWRQRFDFVMEAIQKSERETGERKGHYLNVTAPTP<br>EEMYKRAEYAKEIGAPIIMHDYITGGFCANTGLANWCRDNGMLLHIHRAMHAVLDRNPHH<br>GIHFRVLTKILRLSGGDHLHSGTVVGKLEGDREATLGWIDMMRDSFVKEDRSRGIFFDQD<br>WGSMPGVFPVASGGIHVWHMPALVTIFGDDSVLQFGGGTLGHPWGNAAGAAANRVALEAC<br>VEARNKGVAIEKEGKTVLTEAAKNSPELKIAMETWKEIKFEFDTVDKLDVAHK |
| *Nicotiana tabacum* | IB | >sp\|P00876\|RBL_TOBAC Ribulose bisphosphate carboxylase large chain OS=Nicotiana tabacum OX=4097 GN=rbcL PE=1 SV=2<br>MSPQTETKASVGFKAGVKEYKLTYYTPEYQTKDTDILAAFRVTPQPGVPPEEAGAAVAAE<br>SSTGTWTTVWTDGLTSLDRYKGRCYRIERVVGEKDQYIAYVAYPLDLFEEGSVTNMFTSI<br>VGNVFGFKALRALRLEDLRIPPAYVKTFQGPPHGIQVERDKLNKYGRPLLGCTIKPKLGL<br>SAKNYGRAVYECLRGGLDFTKDDENVNSQPFMRWRDRFLFCAEALYKAQAETGEIKGHYL<br>NATAGTCEEMIKRAVFARELGVPIVMHDYLTGGFTANTSLAHYCRDNGLLLHIHRAMHAV<br>IDRQKNHGIHFRVLAKALRMSGGDHIHSGTVVGKLEGERDITLGFVDLLRDDFVEQDRSR<br>GIYFTQDWVSLPGVLPVASGGIHVWHMPALTEIFGDDSVLQFGGGTLGHPWGNAPGAVAN<br>RVALEACVKARNEGRDLAQEGNEIIREACKWSPELAAACEVWKEIVFNFAAVDVLDK |
| *Spinacia oleracea* | IB | >sp\|P00875\|RBL_SPIOL Ribulose bisphosphate carboxylase large chain OS=Spinacia oleracea OX=3562 GN=rbcL PE=1 SV=1<br>MSPQTETKASVEFKAGVKDYKLTYYTPEYETLDTDILAAFRVSPQPGVPPEEAGAAVAAE<br>SSTGTWTTVWTDGLTNLDRYKGRCYHIEPVAGEENQYICYVAYPLDLFEEGSVTNMFTSI<br>VGNVFGFKALRALRLEDLRIPVAYVKTFQGPPHGIQVERDKLNKYGRPLLGCTIKPKLGL<br>SAKNYGRAVYECLRGGLDFTKDDENVNSQPFMRWRDRFLFCAEALYKAQAETGEIKGHYL<br>NATAGTCEDMMKRAVFARELGVPIVMHDYLTGGFTANTTLSHYCRDNGLLLHIHRAMHAV<br>IDRQKNHGMHFRVLAKALRLSGGDHIHSGTVVGKLEGERDITLGFVDLLRDDYTEKDRSR<br>GIYFTQSWVSTPGVLPVASGGIHVWHMPALTEIFGDDSVLQFGGGTLGHPWGNAPGAVAN<br>RVALEACVQARNEGRDLAREGNTIIREATKWSPELAAACEVWKEIKFEFPAMDTV |
| *Microcystis aeruginosa PCC 9806* | IB | >ARI84004.1 ribulose-1,5-bisphosphate carboxylase/oxygenase large subunit [Microcystis aeruginosa PCC 7806SL]<br>MVQAKSKGFQAGVKDYRLTYYTPDYTPKDTDLLACFRVTPQPGVPPEEAGAAVAAESSTGTWTTVWTDNL<br>TDLDRYKGRCYDIEPVPNEDNQFFCFVAYPLDLFEEGSVTNILTSIVGNVFGFKALRGLRLEDIRFPVAL<br>IKTFQGPPHGITVERDKLNKYGRPLLGCTIKPKLGLSAKNYGRAVYECLRGGLDFTKDDENINSQPFMRW |

| | | |
|---|---|---|
| | | RDRFLFVQEAIVKSQAETNEVKGHYLNVTAPTCEQMMQRAEFAAEIKTPIIMHDYLTGGFTANTTLAKFC<br>RDKGLLLHIHRAMHAVIDRQKNHGIHFRVLAKCLRLSGGDHLHSGTVVGKLEGERGITMGFVDLMREDYV<br>EEDRARGIFFTQDYASLPGVMPVASGGIHVWHMPALVEIFGDDSCLQFGGGTLGHPWGNAPGATANRVAL<br>EACIQARNEGRSLAREGNDVIREACRWSPELAAACELWKEIKFEFEAMDTL |
| *Sinorhizobium meliloti* | IC | >sp\|P58348\|RBL1_RHIME Ribulose bisphosphate carboxylase large chain OS=Rhizobium meliloti (strain 1021) OX=266834 GN=cbbL PE=3 SV=1<br>MNADAKTEIKGRERYKAGVLKYAQMGYWNGDYEPKDTDLIALFRITPQDGVDPIEAAAAV<br>AGESSTATWTVVWTDRLTACDQYRAKAYRVDPVPGTPGQYFCYVAYDLILFEEGSIANLT<br>ASIIGNVFSFKPLKAARLEDMRLPVAYVKTFRGPPTGIVVERERLDKFGKPLLGATTKPK<br>LGLSGKNYGRVVYEGLKGGLDFMKDDENINSQPFMHWRDRYLYCMEAVNHASAVTGEVKG<br>HYLNITAGTMEEMYRRAEFAKELGSVIVMVDLIVGWTAIQSISEWCRQNDMILHMHRAGH<br>GTYTRQKNHGISFRVIAKWLRLAGVDHLHAGTAVGKLEGDPPTVQGYYNVCREMKNEVDL<br>PRGLFFEQDWADLKKVMPVASGGIHAGQMHQLLDLFGDDVVLQFGGGTIGHPMGIQAGAT<br>ANRVALEAMVLARNEGRDIAHEGPEILRAAAKWCKPLEAALDIWGNISFNYTPTDTSDFV<br>PSVTAA |
| *Xanthobacter autotrophicus* | IC | >sp\|A7IGM0\|RBL_XANP2 Ribulose bisphosphate carboxylase large chain OS=Xanthobacter autotrophicus (strain ATCC BAA-1158 / Py2) OX=78245 GN=cbbL PE=3 SV=1<br>MGADAAIGQIKDAKKRYAAGVLKYAQMGYWDGDYQPKDTDVLALFRITPQDGVDAVEAAA<br>AVAGESSTATWTVVWTDRLTAADMYRAKAYKVEPVPGQPGQYFCWVAYDLDLFEEGSIAN<br>LTASIIGNVFSFKPLKACRLEDMRLPVAYVKTFRGPPTGIVVERERLDKFGRPLLGATTK<br>PKLGLSGKNYGRVVYEGLKGGLDFVKDDENINSQPFMHWRDRFLYCMEAVNKAQAETGEV<br>KGHYLNITAGTMEEMYRRAEFAKELGSVVVMVDLIVGWTAIQSISNWCRENDVLLHMHRA<br>GHGTYTRQKGHGISFRVIAKWLRLAGVDHLHTGTAVGKLEGDPMTVQGYYNVCRETVTKT<br>DYTRGIFFDQDWAGLRKVMPVASGGIHAGQMHQLIDLFGEDVVLQFGGGTIGHPDGIQAG<br>AIANRVALETMILARNEGRDIKNEGPEILIEAAKWCRPLRAALDTWGEVTFNYASTDTSD<br>FVPTASVA |
| *Paracoccus denitrificans* | IC | >sp\|A1B2Q2\|RBL_PARDP Ribulose bisphosphate carboxylase large chain OS=Paracoccus denitrificans (strain Pd 1222) OX=318586 GN=cbbL PE=3 SV=1<br>MNEMSKSEITDKKKRYAAGVLKYAQMGYWDGDYQPKDTDILALFRITPQDGVDPIEAAAA<br>VAGESSTATWTVVWTDRLTACDQYRAKAYKVEPVPGQEGQYFCYVAYDLILFEEGSIANV<br>TASIIGNVFSFKPLLAARLEDMRFPVAYMKTFAGPPTGIVVERERLDKFGRPLLGATTKP<br>KLGLSGKNYGRVVYEGLKGGLDFMKDDENINSQPFMHWRDRFLYCMEAVNKATAVTGEVK<br>GHYLNITAGTMEEMYRRAELAKELGSVIVMVDLIVGWTAIQSISNWCRQNDMILHMHRAG<br>HGTYTRQKNHGISFRVIAKWLRMAGVDHLHCGTAVGKLEGDPLTVQGYYNTCREMVNEVD<br>LPRGIFFEQDWGNLKKVMPVASGGIHAGQMHQLLDLFGDDVVLQFGGGTIGHPMGIQAGA |

| | | |
|---|---|---|
| | | TANRVALEAMVLARNEGVDLKTEGPEVLRRAAKWCKPLEAALDVWGNITFNYTSTDTSDF<br>VPTASVS |
| *Cupriavidus necator* | IC | >sp|P0C2C2|RBL1C_CUPNE Ribulose bisphosphate carboxylase large chain, chromosomal OS=Cupriavidus necator OX=106590 GN=cbbL1 PE=1 SV=1<br>MNAPETIQAKPRKRYDAGVMKYKEMGYWDGDYVPKDTDVLALFRITPQDGVDPVEAAAAV<br>AGESSTATWTVVWTDRLTACDMYRAKAYRVDPVPNNPEQFFCYVAYDLSLFEEGSIANLT<br>ASIIGNVFSFKPIKAARLEDMRFPVAYVKTFAGPSTGIIVERERLDKFGRPLLGATTKPK<br>LGLSGRNYGRVVYEGLKGGLDFMKDDENINSQPFMHWRDRFLFVMDAVNKASAATGEVKG<br>SYLNVTAGTMEEMYRRAEFAKSLGSVIIMVDLIVGWTCIQSMSNWCRQNDMILHLHRAGH<br>GTYTRQKNHGVSFRVIAKWLRLAGVDHMHTGTAVGKLEGDPLTVQGYYNVCRDAYTQTDL<br>TRGLFFDQDWASLRKVMPVASGGIHAGQMHQLIHLFGDDVVLQFGGGTIGHPQGIQAGAT<br>ANRVALEAMVLARNEGRDILNEGPEILRDAARWCAPLRAALDTWGDITFNYTPTDTSDFV<br>PTASVA |
| *Rubrivivax gelatinosus* | IC | >tr|I0HVA8|I0HVA8_RUBGI Ribulose bisphosphate carboxylase large chain OS=Rubrivivax gelatinosus (strain NBRC 100245 / IL144) OX=983917 GN=cbbL PE=3 SV=1<br>MNKPHEPGATGDQILDKKQRYSAGVLKYRQMGYWDSDYVPKATDVVCLFRITPQEGVDPI<br>EAAAAVAGESSTATWTVVWTDRLTACDSYRAKAYKVEPVPGRPGEYFAWVAYDLILFEEG<br>SIANMTASLIGNVFSFKPLKAARLEDIQIPVAYVKTFKGPPTGLIVERERLDKFGRPLLG<br>ATTKPKLGLSGRNYGRVIYEGLKGGLDFMKDDENINSQPFMHWRDRFLYVMDGVNKASAA<br>TGEVKGSYLNVTGATMEDIYERAEFAKELGSVVIMVDLIIGWSAIQSIANWARKNDMIVH<br>MHRAGHGTYTRQKNHGVSFRVMAKWLRLAGVDHLHTGTAVGKLEGDPLTVQGYYNVCRDA<br>YTKQDLPRGLFFDQDWADLRKVMPVASGGIHAGQMHQLIDLFGDDVILQFGGGTIGHPAG<br>IQAGAVANRVALEAMVKARNEGRDIKNEGPEILQKAAQFCTPLKQALDTWKDVSFNYAST<br>DQSDYAVTPATSA |
| *Nitrosospira multiformis* | IC | >sp|Q2YB78|RBL_NITMU Ribulose bisphosphate carboxylase large chain OS=Nitrosospira multiformis (strain ATCC 25196 / NCIMB 11849 / C 71) OX=323848 GN=cbbL PE=3 SV=1<br>MSEAITGAERYKSGVIPYKKMGYWEPDYVPKDTDIIAMFRITPQAGVEPEEAAAAVAGES<br>STATWTVVWTDRLTACELYRAKAFRTDPVPNTGEGTKTEQQYFAYIAYDLDLFEPGSIAN<br>LTASIIGNVFGFKAVKALRLEDMRIPVAYLKTFQGPATGIIVERERLDKFGRPLLGATTK<br>PKLGLSGRNYGRVVYEGLKGGLDFMKDDENINSQPFMHWRDRFLYCMEAVNKASAATGEV<br>KGHYLNVTAGTMEEMYERAEFAKSLGSVIIMIDLVIGYTAIQSMAKWARKNDMILHLHRA<br>GNSTYSRQKNHGMNFRVICKWMRMAGVDHIHAGTVVGKLEGDPLMIKGFYDTLRDRHTPV<br>SLEHGLFFEQDWASLNKVMPVASGGIHAGQMHQLLDYLGEDVILQFGGGTIGHPQGIQAG<br>AVANRVALEAMIMARNEGRDYVKEGPQILEEAAKWCTPLKLALDTWKDITFNYESTDTAD<br>FVPSETASV |

| | | |
|---|---|---|
| *Nitrosococcus oceani* | IC | >sp\|Q3JE87\|RBL_NITOC Ribulose bisphosphate carboxylase large chain OS=Nitrosococcus oceani (strain ATCC 19707 / BCRC 17464 / NCIMB 11848 / C-107) OX=323261 GN=cbbL PE=3 SV=1<br>MGKSETIAEGKDRYQAGVIPYKKMGYWEPDYQPKDTDIIAMFRITPQPGVDPEEAAAAVA<br>GESSTATWTVVWTDRLTDCELYRAKAYRADLVPNTGEGTKNEAQYFAYIAYDLDLFEPGS<br>IANLTASIIGNVFGFKAVKALRLEDMRIPVAYLKTFQGPATGVVVERERLDKFGRPLLGA<br>TTKPKLGLSGRNYGRVVYEALKGGLDFVKDDENINSQPFMHWRDRFLYCMEAVNKASAAT<br>GEVKGHYLNVTAATMEDMYERAEFAKSLGSVIIMIDLVVGYTAIQSMAKWARKNDMILHL<br>HRAGNSTYSRQKNHGMNFRVICKWMRMAGVDHIHAGTVVGKLEGDPLMIKGFYDTLLDSH<br>TPTSLEHGLFFDQDWASLNKVMPVASGGIHAGQMHQLIQYLGEDVILQFGGGTIGHPQGI<br>QAGAVANRVALEAMILARNEGRDYVKEGPQILQDAAKWCSPLKAALDTWKDVTFNYESTD<br>TADFVPTATASV |
| *Rhodobacter sphaeroides* | IC | >sp\|P27997\|RBL1_RHOSH Ribulose bisphosphate carboxylase large chain OS=Rhodobacter sphaeroides OX=1063 GN=cbbL PE=1 SV=1<br>MDTKTTEIKGKERYKAGVLKYAQMGYWDGDYVPKDTDVLALFRITPQEGVDPVEAAAAVA<br>GESSTATWTVVWTDRLTACDSYRAKAYRVEPVPGTPGQYFCYVAYDLILFEEGSIANLTA<br>SIIGNVFSFKPLKAARLEDMRFPVAYVKTYKGPPTGIVGERERLDKFGKPLLGATTKPKL<br>GLSGKNYGRVVYEGLKGGLDFMKDDENINSQPFMHWRDRFLYVMEAVNLASAQTGEVKGH<br>YLNITAGTMEEMYRRAEFAKSLGSVIVMVDLIIGYTAIQSISEWCRQNDMILHMHRAGHG<br>TYTRQKNHGISFRVIAKWLRLAGVDHLHCGTAVGKLEGDPLTVQGYYNVCREPFNTVDLP<br>RGIFFEQDWADLRKVMPVASGGIHAGQMHQLLSLFGDDVVLQFGGGTIGHPMGIQAGATA<br>NRVALEAMVLARNEGRNIDVEGPEILRAAAKWCKPLEAALDTWGNITFNYTSTDTSDFVP<br>TASVAM |
| *Rhodopseudomonas palustris* | IC | >sp\|Q219P7\|RBL_RHOPB Ribulose bisphosphate carboxylase large chain OS=Rhodopseudomonas palustris (strain BisB18) OX=316056 GN=cbbL PE=3 SV=2<br>MNESVTIRGKDRYKSGVMEYKKMGYWEPDYEPKDTDIIALFRVTPQDGVDPTEASAAVAG<br>ESSTATWTVVWTDRLTAAEKYRAKCYRVDPVPNSPGQFFAYIAYDLDLFENGSIANLSAS<br>IIGNVFGFKPLKALRLEDMRLPVAYVKTFQGPATGIVVERERMDKFGRPLLGATVKPKLG<br>LSGRNYGRVVYEALKGGLDFTKDDENINSQPFMHWRERFLYCMEAVNKAQAASGEIKGTY<br>LNVTAGTMEEMYERAEFAKQLGSVIIMIDLVIGYTAIQSMAKWARRNDMILHLHRAGHST<br>YTRQRNHGVSFRVIAKWMRLAGVDHIHAGTVVGKLEGDPATTKGYYDICREDYNPMQLEH<br>GIFFEQNWASLNKLMPVASGGIHAGQMHQLLDHLGEDVVLQFGGGTIGHPMGIQAGATAN<br>RVALEAMIMARNEGRDYLHEGEEILAKAALTCTPLKAALETWKNVTFNYESTDMPDYAPT<br>PSVSM |
| *Emiliania huxleyi* | ID | >sp\|Q4G3F4\|RBL_EMIHU Ribulose bisphosphate carboxylase large chain OS=Emiliania huxleyi OX=2903 GN=rbcL PE=3 SV=1 |

| | | |
|---|---|---|
| | | MSQAVESRTRIKSERYESGVIPYAKMGYWDPEYVIKDTDILALFRCTPQPGVDPVEAAAA<br>LAGESSTATWTVVWTDLLTACDLYRAKAFRVDPVPSAADTYFCYIAYDIDLFEEGSLANL<br>TASIIGNIFGFKAVKALRLEDMRFPVALLKTYQGPATGVVVERERMDKFGRPLLGATVKP<br>KLGLSGKNYGRVVFEGLKGGLDFLKDDENINSQPFMRYRERFLYSMEGVNHAACLTGEVK<br>GHYLNTTAATMEDMYERANFARDLGSVIVMIDLVIGYTAIQSMGKWSRDNDVILHLHRAG<br>NSTYSRQKNHGMNFRVICKWMRMSGCDHIHAGTVVGKLEGDPLMIKGFYNTLLDTKTEVN<br>LPQGLFFAQDWASLRKCVPVASGGIHCGQMHQLINYLGDDVVLQFGGGTIGHPDGIQAGA<br>TANRVALECMVLARNEGRDYIAEGPQILRDAAKTCGPLQTALDLWKDITFNYASTDTADF<br>VETATANV |
| *Cyanidium caldarium* | ID | >sp\|P37393\|RBL_CYACA Ribulose bisphosphate carboxylase large chain OS=Cyanidium caldarium OX=2771 GN=rbcL PE=3 SV=2 |
| | | MAQSVQERTRLKNKRYESGVIPYAKMGYWDPNYVVKDTDILALFRVTPQPGVDPIEASAA<br>VAGESSTATWTVIWCDLLTACDVYRAKAYRVDQVPNSPDQYFAYIAYDLDLFEEGSIANL<br>TASIIGNVFGFKALAALRLEDMRIPIGYLKTFQGPATGVVVERERLNMFGKPFLGATVKP<br>KLGLSSKNYGRVVYEGLKGGLNFLKDDENINSQPFMRWRERFLYVMEGVNRASAATGEIK<br>GSYLNVTAATMEEMYNRAACAKEVGSIIIMIDLVIGYTAIQSMAIWARENNMILHLHRAG<br>NSTYARQKNHGINFRVICKWMRMAGVDHIHAGTVVGKLEGDPIIVKGFYNTLLLPKLDVN<br>LPQGLFFEMDWASLRKTVPVASGGIHAGQMHLLLKYLGDDVVLQFGGGTLGHPDGIQAGA<br>TANRVALEAIVLARNEGRDYVNEGPQILKEAARTCGPLQTSLDLWKDISFNFTSTDTADF<br>VETPTANV |
| *Skeletonema costatum* | ID | >tr\|A0A097IV41\|A0A097IV41_SKECO Ribulose bisphosphate carboxylase large chain (Fragment) OS=Skeletonema costatum OX=2843 GN=rbcL PE=3 SV=1 |
| | | LGYWDASYTVKDTDVLALFRITPQPGVDPVEAAAAVAGESSTATWTVVWTDLLTACERYR<br>AKAYRVDPVPNSADVFFAFIAYECDLFEEASLANLTASIIGNVFGFKAVSALRLEDMRIP<br>HSYLKTFQGPATGIIVERERLNKYGTPLLGATVKPKLGLSGKNYGRVVYEGLKGGLDFLK<br>DDENINSQPFMRWRERFLNCMEGINRASAATGEVKGSYLNITAATMEEVYKRAEYAKAVG<br>SIVVMIDLVMGYTAIQSIAYWARENDMLLHLHRAGNSTYARQKNHGINFRVICKWMRMSG<br>VDHIHAGTVVGKLEGDPLMIKGFYDILRLTELEVNLPFGIFFEMDWASLRRCMPVASGGI<br>HCGQMHQLIHYLGDDVVLQFGGGTIGHPDGIQAGATANRVALESMVLARNEGVDYFDQQV<br>GPQILRDAAKTCGPLQTALDLWKDISFDYTSTDTADFAETPTAN |
| *Porphyridium purpureum* | ID | >tr\|A0A343KNV1\|A0A343KNV1_PORPP Ribulose bisphosphate carboxylase large chain OS=Porphyridium purpureum OX=35688 GN=rbcL PE=3 SV=1 |
| | | MSQSVEERTRIKNERYESGVIPYAKMGYWDADYAIKETDVLALFRVTPQPGVDPVEAAAA<br>IAGESSTATWTVVWTDLLTACDLYRAKAYRVDPVPNSPDQFFAYIAYDIDLFEEGSIANL<br>TASIIGNVFGFKAVKALRLEDMRLPIAYLKTFQGPATGVIVERERMNNFGRPVLGATVKP |

| | | |
|---|---|---|
| | | KLGLSGKNYGRVVYEGLKGGLDFLKDDENINSQPFMRWRERFLFCIEGTNRAVAASGEVK<br>GHYLNVTAATQEDMYERAEFAKEVGSIICMIDLVIGYTAIQTMAKWARKNDMILHLHRAG<br>NSTYSRQKNHGMNFRVICKWMRMAGVDHIHAGTVVGKLEGDPLMIKGFYNTLLDSHLPIN<br>LPQGLFFEQNWASLRKVMPVASGGIHCGQMHQLINYLGDDVVLQFGGGTIGHPDGIQAGA<br>TANRVALECMVQARNEGRDYISEGPQILRDAAKTCGPLRTALDLWKDISFNYTSTDTADF<br>LETATANI |
| *Ca.* Endoriftia persephone | II | >tr\|Q0PQU1\|Q0PQU1_9GAMM Ribulose-15-bisphosphate carboxylase/oxygenase form II large subunit (Fragment) OS=Candidatus Endoriftia persephone str. Hot96_1+Hot96_2 OX=394104 PE=3 SV=1<br>VESIGRDLAVSHAQSTQSTRYGVDQTMALDQTNRYSDLSLTEDELIASGDYVLCAYLMKP<br>KSGYGYLEAAAHFAAESSTGTNVEVSTTDDFTKGVDALVYEIDEAKELMKIAYPVDLFDI<br>NIIDGRAMLASFLTLTIGNNQGMGDIEYAKMLDFYMPPKYLRLYDGPAVNIQDMWRILGR<br>PIENGGYIAGTIIKPKLGLRPEPFAEAAYQF |
| *Ca.* Vesicomyosocius okutanii | II | >tr\|A5CWB0\|A5CWB0_VESOH Ribulose-bisphosphate carboxylase form II OS=Vesicomyosocius okutanii subsp. Calyptogena okutanii (strain HA) OX=412965 GN=cbbM PE=3 SV=1<br>MDQSNRYADLSLDEDTLLAEGGHILVAYTMNIMPNFGGYLETAAHFAAESSTGTNVEVST<br>TDDFTKDLDAMVYEIDEVKGIMKIAYPCGLFDRNLIDGRAMVVSFLTLAIGNNQGMGDVK<br>CAQMIDFHVPKQMLDIFDGPSVDITDLWNLLGRDRKNGGYIAGTIIKPKLGLRPQPFAEA<br>AYQFWLGGDFIKNDEPQGNQVYARMKDVMPLVSDAMKRAQDETGEAKIFSANITADDYHE<br>MIARGEYILEAFGENAHHVAFLVDGYVGGCGMVTTARRNFPGQYLHYHRAGHGAITSPSS<br>VRGYTALVLAKLSRLMGASGIHVGTMGYGKMEGGADDRNIAYMIERDSADGPVYHQEWFG<br>MKPTTPIISGGMNALRLPGFFENLGHGNVINTSGGGSYGHIDSPSAGAKSLRQAYDCWMA<br>KADPIEFARDNNEFARAFESFPGDADSLYPGWRDKLGVHK |
| *Ca.* Ruthia magnifica | II | >tr\|A1AWY1\|A1AWY1_RUTMC Ribulose-1,5-bisphosphate carboxylase/oxygenase large subunit OS=Ruthia magnifica subsp. Calyptogena magnifica OX=413404 GN=Rmag_0701 PE=3 SV=1<br>MDQSNRYADLLLDEETLIKEGNHFLVAYTMTPMPGFGGYLETAAHFAAESSTGTNVEVST<br>TDDFTKDLDAMVYEIDEAKGTMKIAYPNNLFDRNLIDGRAMVVSLLTLIIGNNQGMGDVQ<br>CAQIQDFWISRKFLEIFDGPSLDITDLWSILGRSRTDGGYIAGTIIKPKLGLRPKPFSEA<br>AYQFWLGGDFIKNDEPQGNQVYARMKDVTPLVADAMRRAQDETSEAKIFSANITADDHHE<br>MCARADYILETFAENAHHVAFLVDGYVGGCGMITTARRNYPNQYLHYHRAGHGAITSPSS<br>VRGYTAFVLGKLSRLMGASGIHVGTMGYGKMEGGADDRNIAYMIERDSADGPVYHQEWFG<br>MKPTTPIISGGMNALRLPGFFENLGHGNVINTSGGGSYGHIDSPAAGAKSLRQAYDCWMA<br>KADPIEFAKDHNEFARAFESFPNDADSLYPGWRDKLGIHK |
| *Dechloromonas aromatica* | II | >sp\|Q479W5\|RBL2_DECAR Ribulose bisphosphate carboxylase OS=Dechloromonas aromatica (strain RCB) OX=159087 GN=cbbM PE=3 SV=1<br>MDQSNRYADLSLTEAELIAGGQHILCAYKMKPKAGHRYLEAAAHFAAESSTGTNVEVCTT |

| | | |
|---|---|---|
| | | DEFTKGVDALVYHIDEASEDMRIAYPLDLFDRNMTDGRMMMASFLTLTIGNNQGMGDIEH<br>AKMVDFYVPRRGIELFDGPSKDISDLWRMLGRPVKDGGYIAGTIIKPKLGLRPEPFARAA<br>YEFWLGGDFIKNDEPQGNQVFAPLKKVIPLVYDSMKRAMDETGEAKLFSMNITADDHFEM<br>CARADFALEAFGPDADKLAFLVDGYVGGPGMITTARRQYPNQYLHYHRAGHGAVTSPSSK<br>RGYTAYVLAKMSRLQGASGIHVGTMGYGKMEGDKDDRACAYIIERDSYTGPVYHQEWYGM<br>KPTTPIISGGMNALRLPGFFENLGHGNVINTAGGGAYGHIDSPAAGARSLRQAYDCWKAG<br>ADPVEWARDHYEFARAFESFPQDADQLYPGWRHKLRPAA |
| *Rhodoferax ferrireducens* | II | >sp\|Q21YM9\|RBL2_RHOFT Ribulose bisphosphate carboxylase OS=Rhodoferax ferrireducens (strain ATCC BAA-621 / DSM 15236 / T118) OX=338969 GN=cbbM PE=3 SV=1<br>MDQSKRYADLSLQEAALIAGGQHILCAYKMAPKDGLNYLEAAAHFAAESSTGTNVEVCTT<br>DDFTRDVDALVYYVNEATEDMRIAYPLALFDRNITDGRFMLVSFLTLAVGNNQGMGDIKH<br>AKMIDFYVPERVIQMFDGPAKDISDLWRILGRPVKDGGFIVGTIIKPKLGLRPEPFAQAA<br>YQFWLGGDFIKNDEPQGNQVFSPIKKTLPLVYDALKRAQDETGQAKLFSMNITADDHFEM<br>CARADFALETFGADADKLAFLVDGFVGGPGMVTTARRQYPNQYLHYHRGGHGMVTSPSSK<br>RGYTALVLAKMSRLQGASGIHVGTMGHGKMEGAGDDRVMAYMIERDECQGPVYFQKWYGI<br>KPTTPIVSGGMNALRLPGFFDNLGHGNIINTAGGGSYGHLDSPAAGAVSLRQAYECWKAG<br>ADPIEWAKEHREFARAFESFPQDADRLFAGWRDKLGVGA |
| *Ca.* Riegeria galatelae | II | >tr\|G0YWF8\|G0YWF8_9PROT Ribulose-1,5-bisphosphate carboxylase/oxygenase form II (Fragment) OS=Candidatus Riegeria galatelae OX=1045002 GN=cbbM PE=4 SV=1<br>FWLGGDFIKNDEPQGNQTFAPMKKTIPLVADSMKRAQDETGAAKLFSANITADDPFEMIA<br>RGEYILETFGENAHHLAFLVDGFAAGPTAVTTCRRYFPNTFLHYH |
| *Rhodobacter sphaeroides* | II | >sp\|P29278\|RBL2_RHOSH Ribulose bisphosphate carboxylase OS=Rhodobacter sphaeroides OX=1063 GN=cbbM PE=3 SV=3<br>MDQSNRYARLDLQEADLIAGGRHVLCAYVMKPKAGYGYLETAAHFAAESSTGTNVEVSTT<br>DDFTRGVDALVYEIDPEKEIMKIAYPVELFDRNIIDGRAMLCSFLTLTIGNNQGMGDVEY<br>AKMHDFYVPPCYLRLFDGPSMNIADMWRVLGRDVRNGGMVVGTIIKPKLGLRPKPFADAC<br>HEFWLGADFIKNDEPQGNQTFAPLKETIRLVADAMKRAQDETGEAKLFSANITADDHYEM<br>VARGEYILETFGENADHVAFLVDGYVTGPAAITTARRQFPRQFLHYHRAGHGAVTSPQSM<br>RGYTAFVLSKMARLQGASGIHTGTMGYGKMEGEAADKIMAYMLTDEAAEGPFYRQTGWGS<br>KATTPIISGGMNALRLPGFFDNLGHSNVIQTSGGGAFGHLDGGTAGAKSLRQSHEAWMAG<br>VDLVTYAREHRELARAFESFPADADKFYPGWRDRLHRAA |
| *Rhodopseudomonas palustris* | II | >sp\|Q6N0W9\|RBL2_RHOPA Ribulose bisphosphate carboxylase OS=Rhodopseudomonas palustris (strain ATCC BAA-98 / CGA009) OX=258594 GN=cbbM PE=1 SV=1<br>MDQSNRYANLNLKESELIAGGRHVLCAYIMKPKAGFGNFIQTAAHFAAESSTGTNVEVST<br>TDDFTRGVDALVYEVDEANSLMKIAYPIELFDRNVIDGRAMIASFLTLTIGNNQGMGDVE |

| | | |
|---|---|---|
| | | YAKMYDFYVPPAYLKLFDGPSTTIKDLWRVLGRPVINGGFIVGTIIKPKLGLRPQPFANA<br>CYDFWLGGDFIKNDEPQGNQVFAPFKDTVRAVADAMRRAQDKTGEAKLFSFNITADDHYE<br>MLARGEFILETFADNADHIAFLVDGYVAGPAAVTTARRAFPKQYLHYHRAGHGAVTSPQS<br>KRGYTAFVLSKMARLQGASGIHTGTMGFGKMEGEAADRAIAYMITEDAADGPYFHQEWLG<br>MNPTTPIISGGMNALRMPGFFDNLGHSNLIMTAGGGAFGHVDGGAAGAKSLRQAEQCWKQ<br>GADPVEFAKDHREFARAFESFPQDADKLYPNWRAKLKPQAA |
| *Thiomonas intermedia* | II | >sp\|O84917\|RBL2_THIK1 Ribulose bisphosphate carboxylase OS=Thiomonas intermedia (strain K12) OX=75379 GN=cbbM PE=3 SV=1<br>MAHDQSSRYANLDLKESDLIAGGKHILVAYKMKPKAGYDYLATAAHFAAESSTGTNVEVS<br>TTDDFTKGVDALVYFIDEATEDMRIAYPIELFDRNVIDGRFMIVSFLTLVIGNNQGMGDV<br>EYGKMIDFYVPERAIQMFDGPATDISNLWRILGRPIKDGGYIAGTIIKPKLGLRPEPFAQ<br>AAYQFWLGGDFIKNDEPQGNQVFAPVKKVIPLVYDAMKRAQDETGEAKLFSMNITADDYH<br>EMCARADFALEVFGPDADKLAFLVDGYVGGPGMVTTARRQYPNQYLHYHRAGHGAITSPS<br>SKRGYTAFVLAKMSRLQGASGIHVGTMGYGKMEGEGDDRNIAYMIERDECQGPVYFQKWY<br>GMKPTTPIISGGMNALRLPGFFENLGHGNVINTAGGGSYGHIDSPAAGAKSLRQAYECWK<br>AGADPIEYAKEHKEFARAFESFPGDADKLFPGWRDKLGVHK |
| *Halothiobacillus neapolitanus* | II | >sp\|Q9ZHZ4\|RBL2_HALNC Ribulose bisphosphate carboxylase OS=Halothiobacillus neapolitanus (strain ATCC 23641 / c2) OX=555778 GN=cbbM PE=2 SV=1<br>MDQSARYADLSLKEEDLIAGGKHILVAYKMKPKAGHGYLEASAHFAAESSTGTNVEVSTT<br>DDFTKGVDALVYYIDEATEDMRIAYPMDLFDRNVTDGRMMLVSVLTLIIGNNQGMGDIEH<br>AKIHDIYFPERAIQLFDGPSKDISDMWRILGRPIENGGYIAGTIIKPKLGLRPEPFAAAA<br>YQFWLGGDFIKNDEPQGNQVFCPLKKVLPLVYDSMKRAQDETGQAKLFSMNITADDHYEM<br>MARADFGLETFGPDADKLAFLVDGFVGGPGMITTARRQYPNQYLHYHRAGHGMITSPSAK<br>RGYTAFVLAKISRLQGASGIHVGTMGYGKMEGEGDDRNIAYMIERDEAQGPVYFQKWYGM<br>KPTTPIISGGMNALRLPGFFENLGHGNVINTAGGGSYGHIDSPAAGAISLKQAYECWKAG<br>ADPIEFAKEHKEFARAFESFPKDADAIFPGWREKLGVHK |
| *Hydrogenovibrio marinus* | II | >sp\|Q59462\|RBL2_HYDMR Ribulose bisphosphate carboxylase OS=Hydrogenovibrio marinus OX=28885 GN=cbbM PE=1 SV=3<br>MDQSNRYADLTLTEEKLVADGNHLLVAYRLKPAAGYGFLEVAAHVAAESSTGTNVEVSTT<br>DDFTRGVDALVYEIDEAAFGDKGGLMKIAYPVDLFDPNLIDGHYNVSHMWSLILGNNQGM<br>GDHEGLRMLDFLVPEKMVKRFDGPATDISDLWKVLGRPEVDGGYIAGTIIKPKLGLRPEP<br>FAKACYDFWLGGDFIKNDEPQANQNFCPMEVVIPKVAEAMDRAQQATGQAKLFSANVTAD<br>FHEEMIKRGEYVLGEFAKYGNEKHVAFLVDGFVTGPAGVTTSRRAFPDTYLHFHRAGHGA<br>VTSYKSPMGMDPLCYMKLARLMGASGIHTGTMGYGKMEGHNDERVLAYMLERDECQGPYF<br>YQKWYGMKPTTPIISGGMDALRLPGFFENLGHGNVINTCGGGSFGHIDSPAAGGISLGQA |

| | | |
|---|---|---|
| | | YACWKTGAEPIEAPREFARAFESFPGDADKIFPGWREKLGVHK |
| *Thiobacillus denitrificans* | II | >sp\|Q60028\|RBL2_THIDA Ribulose bisphosphate carboxylase OS=Thiobacillus denitrificans (strain ATCC 25259) OX=292415 GN=cbbM PE=1 SV=3<br>MDQSARYADLSLKEEDLIKGGRHILVAYKMKPKSGYGYLEAAAHFAAESSTGTNVEVSTT<br>DDFTKGVDALVYYIDEASEDMRIAYPLELFDRNVTDGRFMLVSFLTLAIGNNQGMGDIEH<br>AKMIDFYVPERCIQMFDGPATDISNLWRILGRPVVNGGYIAGTIIKPKLGLRPEPFAKAA<br>YQFWLGGDFIKNDEPQGNQVFCPLKKVLPLVYDAMKRAQDDTGQAKLFSMNITADDHYEM<br>CARADYALEVFGPDADKLAFLVDGYVGGPGMVTTARRQYPGQYLHYHRAGHGAVTSPSAK<br>RGYTAFVLAKMSRLQGASGIHVGTMGYGKMEGEGDDKIIAYMIERDECQGPVYFQKWYGM<br>KPTTPIISGGMNALRLPGFFENLGHGNVINTAGGGSYGHIDSPAAGAISLRQSYECWKQG<br>ADPIEFAKEHKEFARAFESFPKDADKLFPGWREKLGVHK |
| *Rhodospirillum rubrum ATCC 11170* | II | >sp\|P04718\|RBL2_RHORU Ribulose bisphosphate carboxylase OS=Rhodospirillum rubrum OX=1085 GN=cbbM PE=1 SV=1<br>MDQSSRYVNLALKEEDLIAGGEHVLCAYIMKPKAGYGYVATAAHFAAESSTGTNVEVCTT<br>DDFTRGVDALVYEVDEARELTKIAYPVALFHRNITDGKAMIASFLTLTMGNNQGMGDVEY<br>AKMHDFYVPEAYRALFDGPSVNISALWKVLGRPEVDGGLVVGTIIKPKLGLRPKPFAEAC<br>HAFWLGGDFIKNDEPQGNQPFAPLRDTIALVADAMRRAQDETGEAKLFSANITADDPFEI<br>IARGEYVLETFGENASHVALLVDGYVAGAAAITTARRRFPDNFLHYHRAGHGAVTSPQSK<br>RGYTAFVHCKMARLQGASGIHTGTMGFGKMEGESSDRAIAYMLTQDEAQGPFYRQSWGGM<br>KACTPIISGGMNALRMPGFFENLGNANVILTAGGGAFGHIDGPVAGARSLRQAWQAWRDG<br>VPVLDYAREHKELARAFESFPGDADQIYPGWRKALGVEDTRSALPA |
| *Methanocaldococcu* | III | MDYINLNYRPNEGDLLSCMVIKGENLEKLANEIAGESSIGTWTKVQTMKSDIYEKLRPKV<br>YEIKEIGEENGYKVGLIKIAYPLYDFEINNMPGVLAGIAGNIFGMKIAKGLRILDFRFPA<br>EFVKAYKGPRFGIEGVRETLKIKERPLLGTIVKPKVGLKTEEHAKVAYEAWVGGVDLVKD<br>DENLTSQEFNKFEDRIYKTLEMRDKAEEETGERKAYMPNITAPYREMIRRAEIAEDAGSE<br>YVMIDVVVCGFSAVQSFREEDFKFIIHAHRAMHAAMTRSRDFGISMLALAKIYRLLGVDQ<br>LHIGTVVGKMEGGEKEVKAIRDEIVYDKVEADNENKFFNQDWFDIKPVFPVSSGGVHPRL<br>VPKIVEILGRDLIIQAGGGVHGHPDGTRAGAKAMRAAIEAIIEGKSLEEKAEEVAELKKA<br>LEYWK |
| *Archaeoglobus fulgidus* | III | >sp\|O28635\|RBL_ARCFU Ribulose bisphosphate carboxylase OS=Archaeoglobus fulgidus (strain ATCC 49558 / VC-16 / DSM 4304 / JCM 9628 / NBRC 100126) OX=224325 GN=rbcL PE=1 SV=1<br>MAEFEIYREYVDKSYEPQKDDIVAVFRITPAEGFTIEDAAGAVAAESSTGTWTSLHPWYD<br>EERVKGLSAKAYDFVDLGDGSSIVRIAYPSELFEPHNMPGLLASIAGNVFGMKRVKGLRL<br>EDLQLPKSFLKDFKGPSKGKEGVKKIFGVADRPIVGTVPKPKVGYSAEEVEKLAYELLSG<br>GMDYIKDDENLTSPAYCRFEERAERIMKVIEKVEAETGEKKSWFANITADVREMERRLKL |

| | | |
|---|---|---|
| | | VAELGNPHVMVDVVITGWGALEYIRDLAEDYDLAIHGHRAMHAAFTRNAKHGISMFVLAK<br>LYRIIGIDQLHIGTAGAGKLEGQKWDTVQNARIFSEVEYTPDEGDAFHLSQNFHHIKPAM<br>PVSSGGLHPGNLEPVIDALGKEIVIQVGGGVLGHPMGAKAGAKAVRQALDAIISAIPLEE<br>HAKQHPELQAALEKWGRVTPI |
| *Thermococcus kodakaraensis* | III | >sp\|O93627\|RBL_THEKO Ribulose bisphosphate carboxylase OS=Thermococcus kodakarensis (strain ATCC BAA-918 / JCM 12380 / KOD1) OX=69014 GN=rbcL PE=1 SV=5<br>MVEKFDTIYDYYVDKGYEPSKKRDIIAVFRVTPAEGYTIEQAAGAVAAESSTGTWTTLYP<br>WYEQERWADLSAKAYDFHDMGDGSWIVRIAYPFHAFEEANLPGLLASIAGNIFGMKRVKG<br>LRLEDLYFPEKLIREFDGPAFGIEGVRKMLEIKDRPIYGVVPKPKVGYSPEEFEKLAYDL<br>LSNGADYMKDDENLTSPWYNRFEERAEIMAKIIDKVENETGEKKTWFANITADLLEMEQR<br>LEVLADLGLKHAMVDVVITGWGALRYIRDLAADYGLAIHGHRAMHAAFTRNPYHGISMFV<br>LAKLYRLIGIDQLHVGTAGAGKLEGGKWDVIQNARILRESHYKPDENDVFHLEQKFYSIK<br>AAFPTSSGGLHPGNIQPVIEALGTDIVLQLGGGTLGHPDGPAAGARAVRQAIDAIMQGIP<br>LDEYAKTHKELARALEKWGHVTPV |
| *Pyrococcus horikoshii* | III | >sp\|O58677\|RBL_PYRHO Ribulose bisphosphate carboxylase OS=Pyrococcus horikoshii (strain ATCC 700860 / DSM 12428 / JCM 9974 / NBRC 100139 / OT-3) OX=70601 GN=rbcL PE=1 SV=1<br>MMVLRMKVEWYLDFVDLNYEPGRDELIVEYYFEPNGVSPEEAAGRIASESSIGTWTTLWK<br>LPEMAKRSMAKVFYLEKHGEGYIAKIAYPLTLFEEGSLVQLFSAVAGNVFGMKALKNLRL<br>LDFHPPYEYLRHFKGPQFGVQGIREFMGVKDRPLTATVPKPKMGWSVEEYAEIAYELWSG<br>GIDLLKDDENFTSFPFNRFEERVRKLYRVRDRVEAETGETKEYLINITGPVNIMEKRAEM<br>VANEGGQYVMIDIVVAGWSALQYMREVTEDLGLAIHAHRAMHAAFTRNPRHGITMLALAK<br>AARMIGVDQIHTGTAVGKMAGNYEEIKRINDFLLSKWEHIRPVFPVASGGLHPGLMPELI<br>RLFGKDLVIQAGGGVMGHPDGPRAGAKALRDAIDAAIEGVDLDEKAKSSPELKKSLREVG<br>LSKAKVGVQH |
| *Methanosarcina mazei* | III | >sp\|Q8PXG9\|RBL_METMA Ribulose bisphosphate carboxylase OS=Methanosarcina mazei (strain ATCC BAA-159 / DSM 3647 / Goe1 / Go1 / JCM 11833 / OCM 88) OX=192952 GN=rbcL PE=3 SV=1<br>MRRDYIDIGYSPKETDLVCEFHIEPTAGVNFEEAATHLAGESSIDSWTEIATLSPELAEK<br>LKPHVFYADEGAQTVRVAYSEELFELGSVPQVLSAVAGNILSMKIVDNVRLQDIAFPKSM<br>INEFKGPNFGLPGIRKLVGVQDRPLIGTIVKPKVGLNSEKHAEVAYNSFVGGCDLVKDDE<br>NLSDQKFNSFEKRAELTLKLAEKAESETGEKKMYLCNVTAPTCREMIRRMNFLKDLGASY<br>VMVDIVPAGWTAIQTLREEAEDAGLALHAHRCMHSAYTRNPRHGISMLVVAKLCRLIGLD<br>QLHIGTVVGKMHGEKHEVLNLRDQCVLDKVPADESQHILAQDWRGLKPMFPVASGGLAPT<br>MIPDLYSIFGKDVIMQFGGGIHAHPMGTVAGATACRQALEASLEGISLQDYAKNHKELET<br>ALGKWLKK |

| | | |
|---|---|---|
| *Chlorobaculum tepidum* | IV | >sp\|Q8KBL4\|RBLL_CHLTE Ribulose bisphosphate carboxylase-like protein OS=Chlorobaculum tepidum (strain ATCC 49652 / DSM 12025 / NBRC 103806 / TLS) OX=194439 GN=CT1772 PE=1 SV=1<br>MNAEDVKGFFASRESLDMEQYLVLDYYLESVGDIETALAHFCSEQSTAQWKRVGVDEDFR<br>LVHAAKVIDYEVIEELEQLSYPVKHSETGKIHACRVTIAHPHCNFGPKIPNLLTAVCGEG<br>TYFTPGVPVVKLMDIHFPDTYLADFEGPKFGIEGLRDILNAHGRPIFFGVVKPNIGLSPG<br>EFAEIAYQSWLGGLDIAKDDEMLADVTWSSIEERAAHLGKARRKAEAETGEPKIYLANIT<br>DEVDSLMEKHDVAVRNGANALLINALPVGLSAVRMLSNYTQVPLIGHFPFIASFSRMEKY<br>GIHSKVMTKLQRLAGLDAVIMPGFGDRMMTPEEEVLENVIECTKPMGRIKPCLPVPGGSD<br>SALTLQTVYEKVGNVDFGFVPGRGVFGHPMGPKAGAKSIRQAWEAIEQGISIETWAETHP<br>ELQAMVDQSLLKKQD |
| *Bacillus cereus* | IV | >BAL19741.1 ribulose bisphosphate carboxylase, putative [Bacillus cereus NC7401]<br>MSGIIATYLIHDDSHNLEKKAEQIALGLTIGSWTHLPHLLQEQLKQHKGNVIHVEELAEHEHTNSYLRKK<br>VKRGIIKIEYPLLNFSPDLPAILTTTFGKLSLDGEVKLIDLTFSDELKKHFPGPKFGIDGIRNLLQVHDR<br>PLLMSIFKGMIGRNIGYLKTQLRDQAIGGVDIVKDDEILFENALTPLTKRIVSGKEVLQSVYETYGHKTL<br>YAVNLTGRTFDLKENAKRAVQAGADILLFNVFSYGLDVLQSLAEDDEIPVPIMAHPAVSGAYSASKLYGV<br>SSPLLLGKLLRYAGADFSLFPSPYGSVALEKEEALAISKYLTEDDAFFKKSFSVPSAGIHPGFVPFIVRD<br>FGKDVVINAGGGIHGHPNGAQGGGKAFRAAIDATLQNKPLHEVDDINLHSALQIWGNPSHEVKL |
| *Ca.* Kentron sp. H | IV | >tr\|A0A450V0T7\|A0A450V0T7_9GAMM Ribulose-bisphosphate carboxylase large chain OS=Candidatus Kentron sp. H OX=2126337 GN=BECKH772A_GA0070896_1002221 PE=4 SV=1<br>MPPTDRLQLTGERIRAVYRLTGAVGEASARAEDICVEQTVEFPKDLIDREDILGGIVGKV<br>VSLNETGGRTVEATIAFPREAACSELTQLLNVLFGNISLKPGIRLVGLTLPDGLLSAYRG<br>PRFGRTGLRQILDVPERPLLATALKPLGLEAKELANLAGRFALGGLDMIKDDHGLTDQPF<br>CRFRDRVARCADAVREANVRTGGNCLYFANITAPAGELEARAYFAKTSGTGGVVIAPGLA<br>GFDAMRALADDDALALPILSHPAFLGSFTVHPASGIAHGVLHGRINRLAGADACIFPSYG<br>GRFSFTEAECRDIAEAVASPMGNIKTILPAPAGGMNLERVPELIRFYGNECILLIGGDLH<br>RHEGDLVKGCRKFVDIVEKMV |
| *Microcystis aeruginosa PCC 9806* | IV | >WP_002783691.1 2,3-diketo-5-methylthiopentyl-1-phosphate enolase [Microcystis aeruginosa]<br>MTIIVDYRFPPAINAEKQAKTIAIGQTAGTWSDRHSHRQEQLQQHLAEVVGIREEADGYKVARVRFPQIN<br>VENDIASLLTMIFGKYSMAGAGKVVGVYLPESYGTKAKLGITGIRQRLGVYDRPLVMAIFKPALGLSAQD<br>HADILREVAFAGLDVIKDDEIMADIPVAPTHERLDCCRRVLEEVRQQTGRNVLYAVNVTGKADELQRKAR<br>LLVKHGANALLLNVLTYGFSVLEALASDPAIDVPIFAHPAFAGAMCAGSDTGLAYSVVLGTMMAHAGADA<br>VLYPAAYGSLPFDPQEEGKIRDILRDRNVFPVPSAGIRPGIVPQVLRDYGRNVILNAGTGIMDHPSGPAS<br>GVRAFFEALARIEAGDSFDPANLPEGALKQAILEWG |

| | | |
|---|---|---|
| *Rhodospirillum rubrum ATCC 11170* | IV | >WP_011389751.1 ribulose-bisphosphate carboxylase [Rhodospirillum rubrum]<br>MTDRLRATYRVKATAASIEARAKGIAVEQSVEMPLSAIDDPAVLDGIVGVVEEITERGEDCFEVRLALST<br>ATIGGDAGQLFNMLFGNTSLQDDTVLLDIDLPDDLLASFGGPNIGAAGLRARVGASADRALTCSALKPQG<br>LPPDRLADLARRMALGGLDFIKDDHGMADQAYAPFASRVGAVAAAVDEVNRQTGGQTRYLPSLSGHLDQL<br>RSQVRTGLDHGIDTFLIAPMIVGPSTFHAVVREFPGAAFFAHPTLAGPSRIAPPAHFGKLFRLLGADAVI<br>FPNSGGRFGYSRDTCQAVAEAALGPWGGLHASLPVPAGGMSLARVPEMIATYGPDVIVLIGGNLLEARDR<br>LTEETAAFVASVAGAASRGCGLAP |
| *Geobacillus kaustophilus HTA426* | IV | >WP_011230454.1 2,3-diketo-5-methylthiopentyl-1-phosphate enolase [Geobacillus kaustophilus]<br>MSAVMATYLLHDETDIRKKAEGIALGLTIGTWTDLPALEQEQLRKHKGEVVAIEELGESERVNAYFGKRL<br>KRAIVKIAYPTVNFSADLPALLVTTFGKLSLDGEVRLLDLEFPDEWKRQFPGPRFGIDGIRDRVGVHNRP<br>LLMSIFKGMIGRDLAYLTSELKKQALGGVDLVKDDEILFDSELLPFEKRITEGKAALQEVYEQTGKRTLY<br>AVNLTGKTFALKDKAKRAAELGADVLLFNVFAYGLDVLQALREDEEIAVPIMAHPAFSGAVTPSEFYGVA<br>PSLWLGKLLRLAGADFVLFPSPYGSVALEREQALGIARALTDDQEPFARAFPVPSAGIHPGLVPLIIRDF<br>GLDTIVNAGGGIHGHPDGAIGGGRAFRAAIDAVLAGRPLRAAAAENEALQKAIDRWGVVEVEA |
| *Archaeoglobus fulgidus* | IV | >KUJ93818.1 Ribulose bisphosphate carboxylase-like protein [Archaeoglobus fulgidus]<br>MQLGVRLRFQKFEHPEANPEALPEGIDPEEYIIGTYYMSFPKGMNPFEITQVLALEQSTGTWLPVPGETP<br>EVRRKHVAKVVGVYEIPDYEIMVPQEVDWRNFIVQIAFPWRNIGSKLSMLFSTVVGNISMAPKLKLLDLR<br>FPKEFVKGFKGPKFGIEGVRDVLGVKDRPLLNNMIKPDVYSPPDLGAKLAYEVARGGVDIIKDDELLANP<br>EFNRIEERVPKFMEAIDRADEEKGEKTLYAVNVTADLPEVLENAERAIELGANCLLVNYLATGFPVLRAL<br>AEDESIKVPIMAHMDVAGAYYVSPISGVRSTLILGKLPRLAGADIVVYPAPYGKAPMLEEKYVEVAKNHR<br>YPFYNIKPCFPMPSGGIAPIMVPKLVNTLGKDFVVAAGGGIHAHPEGPAAGARAFRQAIDAAMQGYTDLR<br>KYAEENNLQELLKALQL |
| *Pseudomonas putida* | IV | >WP_064491904.1 ribulose 1,5-bisphosphate carboxylase [Pseudomonas putida]<br>MVDHKAGREEFTARFFVESSYPIERVSEVIAGEQSSGTFLSLPGESAELKERSRARVVAIEPLPSVSSPS<br>LHSDYLARHNPSETYHRGEVTIAFPTANIGTNIPALLTTIAGNLFEIGEVSGLRVLDLELPERFASGFIG<br>PQFGIEGTRALAGVHDRPIIGTIVKPSIGLTPEQTASVVDELCAGGIDFIKDDELLIDPTYASFDARLGA<br>VMPVLHKHADRLGRMPMYAINISGSIDEMMRRQDAVLEAGGTCVMLCLNWVGHSAVEHIRKHAQLPIHGH<br>RNGWGALTRYPQLGFSFEVYQKIWRLAGIDHLHVNGIQGKFYEADDSVISSAKSCLAPFAGMRPLMPVFS<br>SGQWAGQAPELFNRLQSVDLMHLAGGGIIGHPMGIEAGVQSMREGWEAAVQHMPLERYAEGRPALSAALQ<br>KFGGK |
| *Mesorhizobium loti R7ANSxAA22* | IV | >WP_032929074.1 MULTISPECIES: ribulose 1,5-bisphosphate carboxylase [Mesorhizobium]<br>MITLTYRIETSESIEALAAKIASDQSTGTFVALPGETEELKARVAARVLAIRHLPDAVQPSIPEASNGPF |

| | | |
|---|---|---|
| | | KRADVDIAFPFDAIGTDLSALMTIAIGGTYSIKGLSGIRIVDMKLPDDYRGAHPGPQFGVAGSRKLTGVE<br>GRPIIGTIVKPALGLRPYETAEMVGELIAAGVDFIKDDEKLMSPAYSPLAERVKAIMPLVRDHEQKTGKK<br>VMYAFGISHADPDEMMRNHDLVLKAGGNCAVININSIGFGGMAYLRKRSSLVLHAHRNGWDILTRHPGLG<br>MDFKVWQQFWRLLGVDQFQINGIASKYWEPDASFIESFKAVTTPIFSPDDCALPVAGSGQWGGQAPETYQ<br>RTGRTVDLLYLCGGGIVSHPDGPGAGVRAVRQAWQAAVDGIPLAKYARSHAELARSIEKFGDGKAA |

[1] According to literature.
